# An evolutionarily conserved role for separase in the regulation of nuclear lamins

**DOI:** 10.1101/2025.02.19.638993

**Authors:** Francesca Cipressa, Gaëlle Pennarun, Giuseppe Bosso, Serena Rosignoli, Liliana Tullo, Schiralli Nadia, Chiara Borghi, Alessandro Paiardini, Michael Lewis Goldberg, Pascale Bertrand, Giovanni Cenci

**Author notes:** **Corresponding authors:** Francesca Cipressa, Giovanni Cenci.

## Abstract

Separase is a well conserved endopeptidase that facilitates sister chromatid separation at the metaphase-anaphase transition by cleaving cohesins. Beyond its role in chromosome segregation, Separase also participates in various biological processes, including chromatin organization and replication, centrosome disengagement and duplication, cytokinesis, and telomere capping. Here, we report that the loss of *Drosophila* separase (*Sse*) function induces significant changes in global protein expression and affects the protein levels of both A/C-type lamin C (*LamC*) and B-type lamin Dm0 (*Dm0*). We further demonstrate that SSE physically interacts with lamins and colocalizes with them at the nuclear envelope during interphase. Additionally, loss of SSE activity disrupts nuclear organization in larval muscles and impairs locomotion in adult flies. Notably, similar to SSE in flies, depletion of human separase (*ESPL1*) in SV40 fibroblasts leads to misshapen nuclei and increased levels of lamin A. Moreover, we show that ESPL1 interacts with lamin A in human fibroblasts, suggesting that the functional interaction between Separase and lamins is evolutionarily conserved across different organisms.

## INTRODUCTION

Separase is a conserved cysteine protease that facilitates sister chromatid separation at the metaphase-anaphase transition by cleaving the cohesin subunit Scc1/Rad21/Mcd1 ^1^. For most of the cell cycle, separase is bound to its chaperone, securin, which inhibits its protease activity ^2^. In addition to securin, separase is regulated through multiple mechanisms, including phosphorylation, autocleavage, and isomerization ^2, 3, 4, 5^.

Beyond its role in chromatid cohesion resolution, separase participates in various cellular functions. It cleaves kendrin, also known as pericentrin, a centrosomal scaffold protein required for recruiting centrosomal components, and is involved in proper spindle assembly and elongation in budding yeast ^6, 7, 8^. Studies in *C. elegans* embryos have demonstrated that separase localizes to the ingressing furrow and midbody during cytokinesis, where it regulates RAB-11-positive vesicle trafficking at the cleavage furrow and midbody ^9^. Moreover, separase is essential for double-strand break repair, where it locally cleaves cohesins to facilitate homology-directed repair (HDR), thereby preventing oncogenic transformation ^10, 11^.

The *Drosophila* genome contains a single separase-encoding gene, *Sse*. However, unlike separase in other model organisms, *Drosophila* SSE lacks the extensive N-terminal regulatory domain found in non-dipteran species. This region appears to have evolved into *thr*, a gene encoding the protein Thr, which binds to SSE and is essential for sister chromatid separation during mitosis ^12, 13^. Previous studies on *Drosophila* separase mutants revealed that its inhibition leads to extensive endoreduplication in mitotic cells, consistent with its role in regulating sister chromatid cohesion ^13^. Further research has shown that SSE is also required for chromosome separation, homolog and sister chromatid disjunction during male meiosis ^14, 15^, polar body formation in female meiosis ^16^, epithelial reorganization ^17^, and telomere capping—an evolutionarily conserved function also observed in human cells ^18^.

Lamins are type V intermediate filament proteins located on the inner surface of the nuclear envelope in metazoan cells, where they contribute to nuclear architecture and interact with chromatin. In addition to maintaining nuclear shape, lamins play key roles in nuclear genome organization, epigenetic histone mark maintenance and distribution, transcriptional regulation and DNA synthesis ^19, 20{Etourneaud, 2021^ ^#59, 21, 22, 23^}. Human lamins are classified into A- and B-types: A-type lamins (A,C and C2) arise from alternative splicing of the *LMNA* gene and are expressed in differentiating cells, whereas B-type lamins (lamins B1, B2 and B3) are encoded by separate genes and exhibit either ubiquitous expression (lamins B1 and B2) or are expressed only in spermatids (Lamin B3) ^24, 25^}. Mutations in nuclear lamin genes lead to a diverse group of human disorders known as laminopathies, which manifest in phenotypes such as premature aging, cardiomyopathy, neuropathy, and lipodystrophy ^26^.

In *Drosophila*, the single A-type lamin, lamin C (*LamC*), is encoded by the *LamC* gene, while the B-type lamin, lamin Dm0 (*Dm0*), is encoded by *LamDm0*. Dm0 is ubiquitously expressed, and its loss results in lethality at the pupal stage, with rare sterile escapers exhibiting locomotion impairments and reduced lifespan ^27^. *Drosophila* LamC is expressed later in embryogenesis, and mutations in *LamC* are lethal, with rare survivors displaying severe muscle defects ^20, 28^. Expression of human mutant pathogenic lamins A/C in *Drosophila* leads to muscle homeostasis defects, including alterations in nuclear shape, nuclear lamina component mislocalization, reduced larval mobility, and disruptions in the nucleus-cytoskeleton connection ^28, 29, 30^.

Here, we demonstrate for the first time that separase physically interacts with lamins in *Drosophila* and regulates their expression. Furthermore, the loss of separase affects nuclear morphology and impairs muscle function in *Drosophila* larvae and adults. Additionally, we show that the relationship between separase and A-type lamins is conserved in human cells, suggesting that this endopeptidase has evolved an evolutionarily conserved function as a lamin-binding factor and regulator.

## RESULTS

### Sse Regulates Global Protein Expression and Nuclear Lamina Component Levels

Previous studies in *Drosophila* have shown that the loss of separase activity affects the expression and localization of several proteins required for chromatin organization and segregation at the post-translational level ^14, 18^. While the SSE-dependent regulation of some of these factors was expected, as they are direct targets of SSE peptidase activity, the mechanism by which SSE influences the levels of other proteins remains unclear.

To further investigate the role of SSE in global protein expression, we performed mass spectrometry (MS) analysis on third instar larval brain extracts from the *diplo -fused telomeres* (*dft*) null allele of the *Sse* gene (*Sse^dft^*) and the *Oregon-R* (Or-R) strain as a wild-type control ^18^. To distinguish protein expression changes specifically dependent on Sse function from those resulting from general mitotic defects, we simultaneously analyzed protein extracts from a trans-heteroallelic mutant combination of the *diamond (dind)* gene, an essential *Drosophila* gene required for mitosis ^31^. *dind* mutant neuroblasts exhibit severe mitotic defects, including abnormalities in chromosome morphology and number, centrosome and mitotic spindle disorganization, chromosome segregation failure, and metaphase delay, all of which collectively impair mitotic progression.

After filtering for cell cycle regulators that were also differentially expressed in *dind* mutants, we identified 4,622 proteins with altered expression in *Sse^dft^* mutants compared to controls. Of these, 402 were upregulated (log2FC > 0.4) and 482 were downregulated (log2FC < -0.4) (Supplementary Figure 1A and B). Western blot analysis of two upregulated factors identified by MS, *Hu-li tai shao* (Hts), the *Drosophila* homolog of Adducin, and Pericardin (Prc), a heart collagen, confirmed a two-fold increase in their expression in *Sse^dft^* larval brain extracts compared to Or-R controls, validating our MS results (Supplementary Figure 2A and B).

Functional enrichment analysis of the upregulated proteins revealed significant enrichment for factors involved in muscle system function (Figure 1A and Supplementary Table 1). Conversely, as expected given the role of SSE in cell cycle regulation and its interactions with DNA, the downregulated proteins were enriched for factors involved in mitotic cell cycle regulation and chromatin organization (Supplementary Table 2). Consistent with our previous findings ^18^, we identified HP1a among the downregulated proteins, further validating our MS analysis in identifying protein modulation specifically dependent on SSE function.

**Figure 1.**
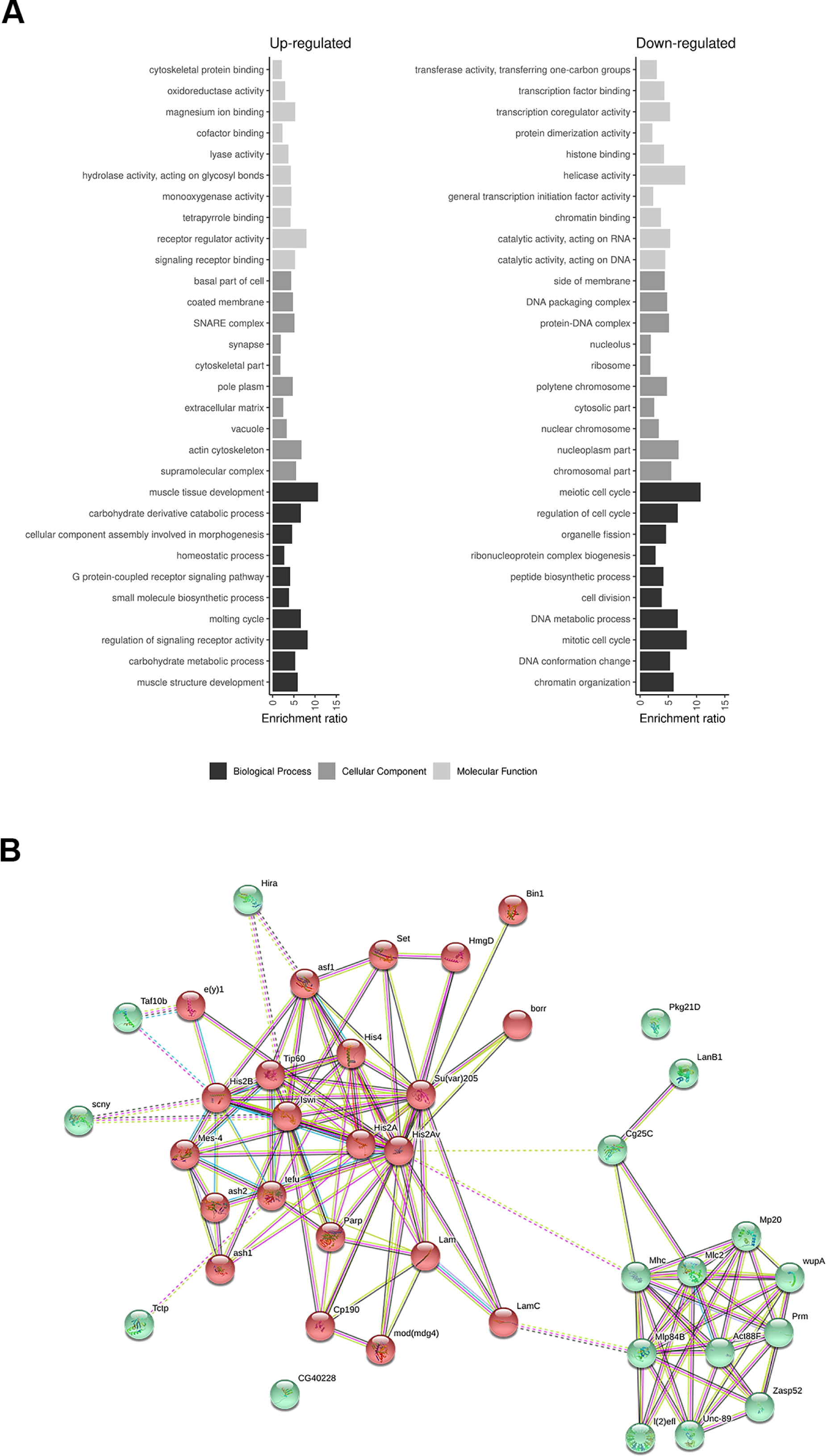
Analysis of protein expression in the absence of separase activity. (A) Bar plot displaying the top 10 enriched Gene Ontology (GO) categories among differentially expressed proteins. The x-axis represents the ratio of observed to expected proteins for each GO category (y-axis). Only categories with an FDR < 0.05 are shown. A complete list of differentially expressed proteins can be found in Supplementary Tables 1 and 2. (B) Interaction network analysis of proteins associated with the enriched GO terms “Muscle structure development” (green spheres) and “Chromatin Organization” (red spheres). A k-means clustering algorithm was applied to group proteins into distinct clusters. Dashed lines indicate interactions between proteins of different clusters. Known interactions retrieved from curated databases or experimental studies are depicted in light blue and light purple, whereas green, red, and blue lines represent predicted interactions based on genomic neighborhood co-occurrence, fusion events, and gene co-occurrence patterns across genomes, respectively. Yellow lines highlight proteins frequently mentioned together in literature, black lines indicate co-expression relationships, and violet lines denote protein homology.

An additional functional enrichment analysis (Supplementary Figure 3 and Supplementary Tables 1 and 3) further confirmed the presence of proteins involved in muscle structure and fiber organization. These proteins are classified under the following Gene Ontology (GO) categories: Muscle System Process (GO:0003012), Muscle Structure Development (GO:0061061), and Supramolecular Fiber Organization (GO:0097435).

To explore potential functional correlations among differentially expressed proteins, we performed a network pathway enrichment analysis using the STRING tool (Figure 1B). This analysis focused on proteins associated with the GO categories Muscle Structure Development (GO:0061061) and Chromatin Organization (GO:0006325), which represented the most significantly enriched categories for up- and down-regulated proteins, respectively. A k-means clustering algorithm was applied to the network to better distinguish distinct pathways. Interestingly, LamC emerged as a central node connecting these two pathways. On one hand, LamC functionally interacts with the Muscle LIM protein (Mlp84B), while on the other, it binds Dm0, highlighting a possible regulatory link between chromatin organization and muscle structure development.

### SSE Regulates Nuclear Lamin Protein Levels

To validate the MS data suggesting a functional relationship between separase (SSE) and lamins, we performed Western blot (WB) analysis on total protein extracts from the *Sse^dft^* mutant. Compared to the Or-R control strain, loss of SSE in mutant neuroblast cells resulted in a 60% reduction in Dm0 protein expression (Figure 2A, C). Conversely, LamC protein levels showed a threefold increase in the same extracts (Figure 2B, D).

**Figure 2.**
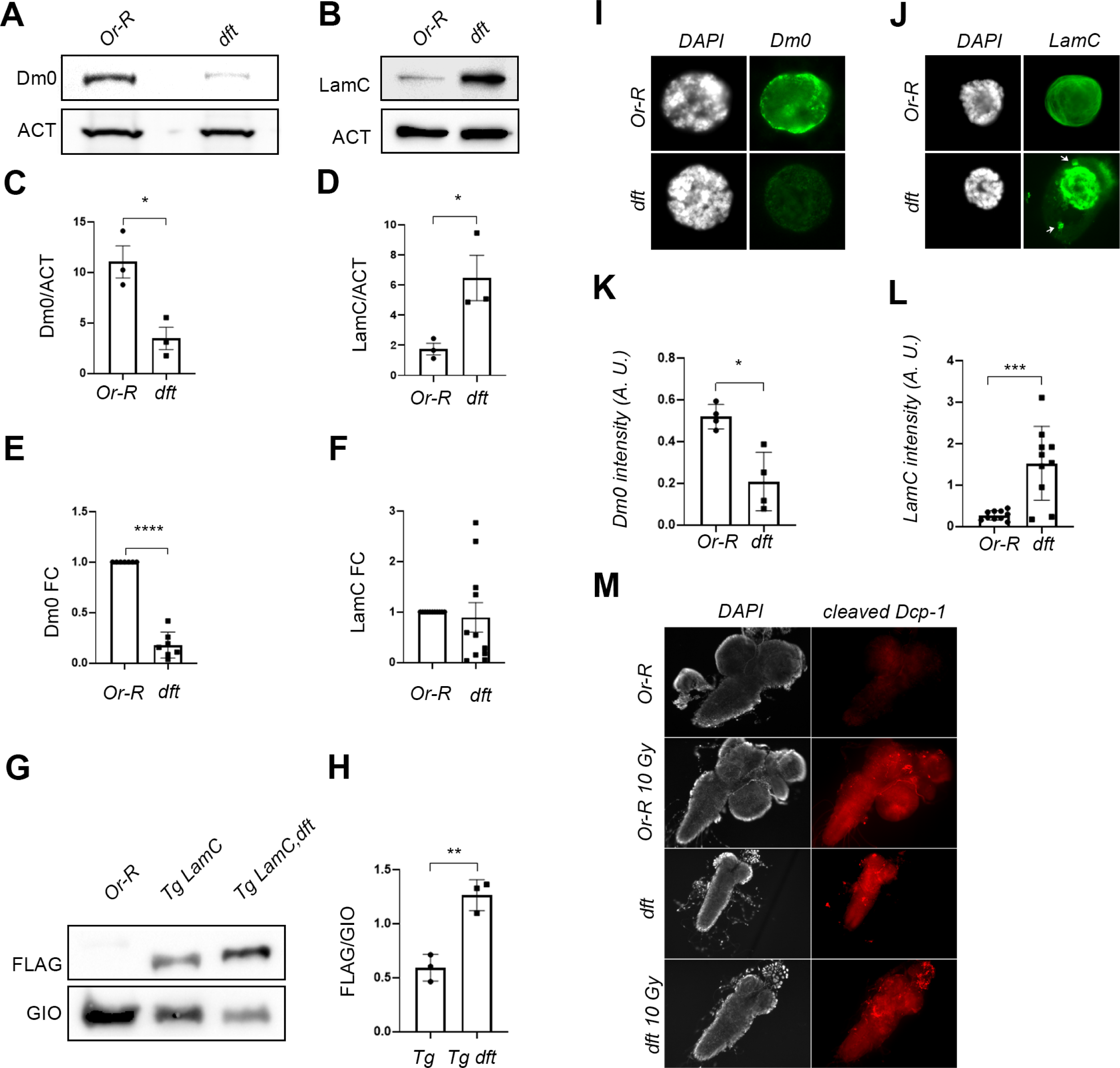
Separase regulates nuclear lamin protein levels. (A, B) Western blot analysis of total protein extracts from wild-type Oregon-R (Or-R) and *Sse^dft^* mutant (*dft*) third instar larval neuroblasts, probed with anti-Dm0 (A) and anti-LamC (B) antibodies. Anti-Actin antibody was used as a loading control. (C, D) Quantification of Dm0 (C) and LamC (D) protein signals from western blot analysis. Data were collected from three independent experiments (*p<0.05, **p<0.01; Student’s t-test). (E, F) qPCR quantification of Dm0 (E) and LamC (F) mRNA expression. At least three independent experiments were conducted (*p<0.05, **p<0.01; Student’s t-test). (G, H) Western blot analysis of FLAG-tagged LamC expression in wild-type (Tg LamC) and *Sse^dft^* mutant (Tg LamC dft) backgrounds (G), with quantification of transgenic LamC expression levels (H). Three independent experiments were performed (*p<0.05, **p<0.01; Student’s t-test). anti-Giotto (GIO) was used as a loading control. (I, J) Immunofluorescence staining with anti-Dm0 (I) and anti-LamC (J) antibodies (green) in intact nuclei from Or-R and *Sse^dft^* third instar larval salivary glands. DAPI was used to stain DNA. Note the presence of LamC aggregates in the *Sse^dft^* nuclei (arrows). (K, L) Quantification of Dm0 (K) and LamC (L) fluorescence intensity from salivary gland immunostaining (***p<0.001; Student’s t-test). (M) Immunostaining with anti-cleaved caspase Dcp-1 (red) in larval brains from untreated and IR-treated (10 Gy) Oregon-R controls, and from untreated (dft) and IR-treated (10 Gy dft) *Sse^dft^* mutant brains.

To determine whether the increase in LamC protein levels was due to a transcriptional regulation, we performed qPCR analysis. Our results showed that LamC mRNA levels in *Sse^dft^* mutants were comparable to those in the Or-R control (Figure 2F), suggesting that the increase in LamC occurs at the post-transcriptional level. In contrast, Dm0 transcript levels were significantly reduced in *Sse^dft^* mutants (Figure 2E), indicating that SSE regulates lamins through distinct mechanisms.

Immunofluorescence analysis of salivary gland nuclei further confirmed the regulatory role of SSE on both Dm0 and LamC. SSE depletion led to a strong reduction in perinuclear Dm0 localization (Figure 2I), whereas LamC staining was significantly increased (Figure 2I–L). Interestingly, in addition to the overall LamC up-regulation, *Sse^dft^* mutant salivary gland nuclei exhibited extranuclear LamC aggregates that were absent in control cells. This suggests that LamC may form misfolded filaments in the absence of SSE. Furthermore, expression of a catalytically inactive SSE (SSE^DH^) in an *Sse^dft^* mutant background failed to completely restore wild-type LamC protein levels, suggesting that SSE-mediated LamC regulation requires its proteolytic activity (Supplementary Figure 4).

To rule out the possibility that the increased LamC levels in *Sse^dft^* mutants resulted from a second-site mutation, we generated transgenic flies expressing N-terminal V5- and C-terminal Flag-tagged LamC under the control of the *Drosophila* tubulin promoter. WB and immunofluorescence analyses confirmed that transgenic LamC was correctly expressed, localized to the nuclear envelope and functionally incorporated into the salivary gland nuclei. Furthermore, these flies were viable and fertile, suggesting that the molecular tags did not alter LamC function (Supplementary Figure 5A, B). We then crossed V5-LamC-Flag transgenic flies with *Sse^dft^* mutants to analyze the transgenic LamC expression (using an anti-V5 antibody) in the absence of *Sse*. As expected, LamC levels were significantly higher in *Sse^dft^* ; *[V5-LamC-Flag]* recombinants than in flies carrying only the transgene, further confirming SSE-mediated regulation of LamC (Figure 2G, H).

### SSE Does Not Regulate LamC via Apoptosis

Lamins are typically cleaved in a caspase-dependent manner during apoptosis to facilitate nuclear lamina disassembly ^32, 33^. Recent studies have demonstrated that premature separase activation can induce Death in Mitosis (DiM), triggering separase-mediated cleavage of pro-survival factors MCL1 and BCL-X, leading to apoptosis ^34^.

Our MS analysis identified the apoptotic initiator caspase Dronc, along with the effector caspases Dcp-1 and Drice, as downregulated proteins in *Sse^dft^* mutants (Table 1). This raised the question of whether the observed upregulation of LamC resulted from reduced apoptosis upon loss of SSE function.

To investigate this, we immunostained wild-type and *Sse^dft^* mutant neuroblasts with an anti-cleaved Dcp-1 antibody, which detects the 22 kDa active fragment of Dcp-1. In untreated control larval brains, cleaved Dcp-1 was almost undetectable, whereas IR-treated cells showed a robust Dcp-1 signal, indicating apoptosis induction (Figure 2M). *Sse^dft^* mutant brains exhibited basal levels of cleaved Dcp-1 staining, which significantly increased after IR treatment, suggesting that apoptosis still occurs in mutant cells. These findings rule out the possibility that LamC upregulation results from reduced apoptosis.

### SSE Physically Interacts with Lamins

To further investigate the relationship between SSE and nuclear lamins, we tested whether SSE physically interacts with lamins using GST pull-down assays. These experiments showed that bacterially purified GST-SSE, but not its interacting partner PIM, successfully precipitated both endogenous Dm0 and LamC proteins from total neuroblast extracts (Figure 3A, B).

**Figure 3.**
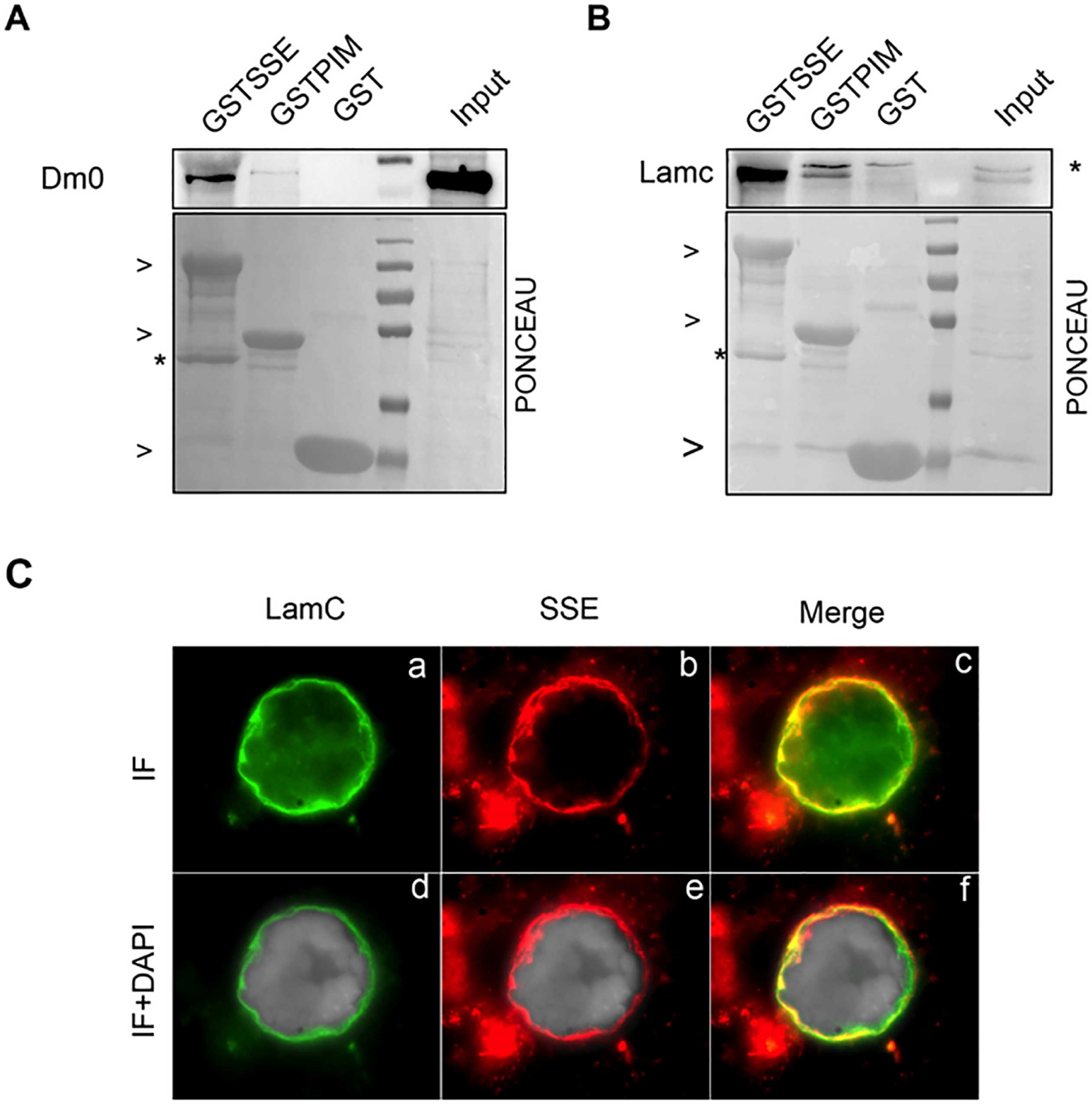
Separase interacts with lamins at the nuclear envelope and influences lamin localization. (A) GST pull-down assay using total protein extracts from wild-type third instar larval brains with GST, GSTPIM, and GSTSSE as baits. Interaction with Dm0 was assessed by western blot using an anti-Dm0 antibody. Input (IN) represents 1/10 of total protein extracts. (B) Western blot analysis with anti-LamC antibody following GST pull-down from brain extracts using GST, GSTPIM, and GSTSSE as baits. Input (IN) represents 1/10 of total protein extracts. The asterisk indicates a non-specific band (C) Immunofluorescence staining of intact nuclei from Or-R salivary glands using anti-LamC (green) and anti-SSE (red) antibodies. Co-localization is observed as yellow staining in the merged images (c and f). DNA is counterstained with DAPI (gray in d, e, and f).

Finally, immunolocalization analysis of salivary gland nuclei revealed that SSE co-localizes with LamC at the nuclear envelope, further supporting a physical interaction between these two proteins (Figure 3C).

However, despite our extensive efforts, we were unable to determine how SSE binding to LamC explains the accumulation of LamC upon SSE loss. We conducted several experiments to test whether LamC is a direct target of SSE peptidase activity, but these efforts did not yield conclusive results (See Discussion, and Supplementary Information).

### Separase-Mediated LamC Regulation is Essential for Muscle Physiology

Since LamC homeostasis plays a crucial role in muscle physiology in Drosophila ^29, 35, 36^, we investigated whether SSE depletion affects muscle function and organization in both larvae and adults.

Locomotion analysis revealed that *Sse^dft^* mutant larvae exhibited a ∼30% reduction in the number of peristaltic contractions per minute compared to controls, indicating that loss of SSE impairs larval motility (Figure 4A). Similarly, RNAi-mediated knockdown of *Sse* in adult muscles using the Mef2-GAL4 driver resulted in a significant reduction in locomotion, as measured by the negative geotaxis assay in 3-day-old flies (Figure 4B). This finding confirms that *Sse* depletion affects muscle function in both larval and adult stages.

**Figure 4.**
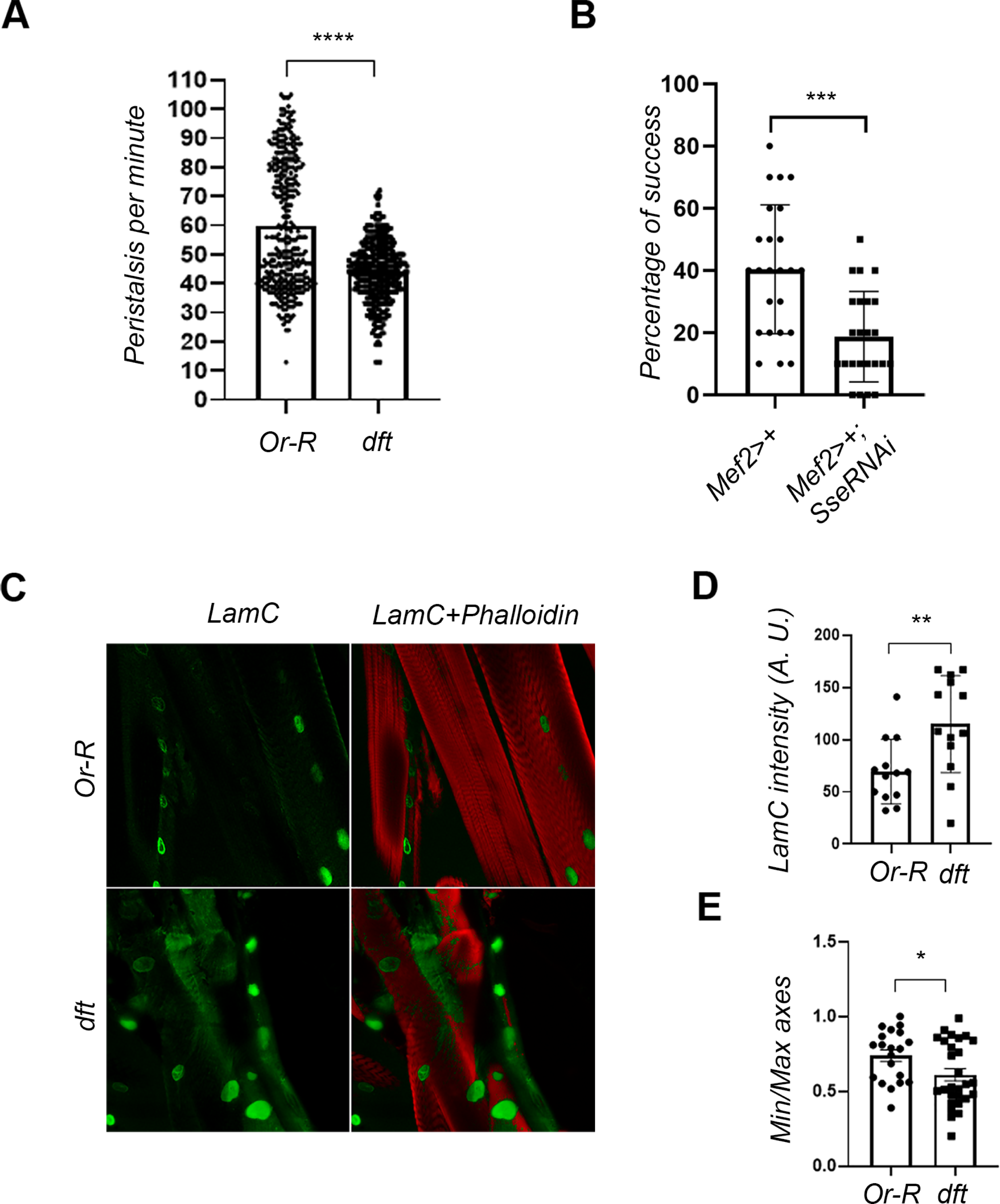
Loss of separase affects locomotion and disrupts Lamin C localization in Drosophila muscle cells. (A) Peristalsis count in *Sse^dft^* mutant larvae, showing a significant reduction in peristalsis per minute in SSE-depleted third instar larvae compared to Or-R controls (***p<0.001; Student’s t-test). (B) Climbing assay assessing locomotion in adult flies with muscle-specific SSE depletion achieved via Mef2-GAL4-driven UAS-Sse dsRNA expression (***p<0.001; Student’s t-test). (C) Immunostaining of larval muscle fillets with anti-LamC (green) in control (Or-R) and *Sse^dft^* mutant larvae. Muscular fibers are counterstained with fluorescently labeled phalloidin (red). (D, E) Quantification of LamC signal intensity (D) and nuclear circularity (E) from muscle fillet preparations (*p<0.05, **p<0.01; Student’s t-test).

To further explore the molecular basis of these defects, we performed anti-LamC immunofluorescence on third instar larval body wall muscles, followed by phalloidin staining. This analysis revealed a significant increase in LamC levels at the muscle nuclear rim in *Sse^dft^* mutants compared to Or-R controls (Figure 4C). This suggests that locomotion defects in *Sse^dft^* mutants may result from LamC accumulation at the nuclear periphery.

Additionally, we observed a reduction in nuclear circularity in *Sse^dft^* mutant muscle cells, compared to wild-type tissue (Figure 4D). This suggests that separase plays a crucial role in maintaining proper nuclear morphology in muscle cells.

### Separase-Mediated Nuclear Lamina Regulation Is Conserved in Human Cells

To determine whether the functional relationship between separase and lamin is conserved in human cells, we depleted human separase, Extra Spindle Pole Bodies-Like 1 (ESPL1), in SV40-fibroblasts using siRNAs and analyzed nuclear morphology and lamin regulation via immunofluorescence. In addition to endoreplication, depletion of ESPL1 resulted in the formation of misshapen nuclei (Supplementary Figure 7), a phenotype previously observed in mammalian cells lacking separase ^5, 37^. We also observed a statistically significant increase in lamin A levels in human SV40-fibroblasts (Figure 5A). These results were further validated by WB blot analysis (Figure 5B) and are consistent with the increased levels of LamC observed in Drosophila upon SSE depletion, suggesting that separase-dependent regulation of A-type lamins is evolutionarily conserved.

**Figure 5.**
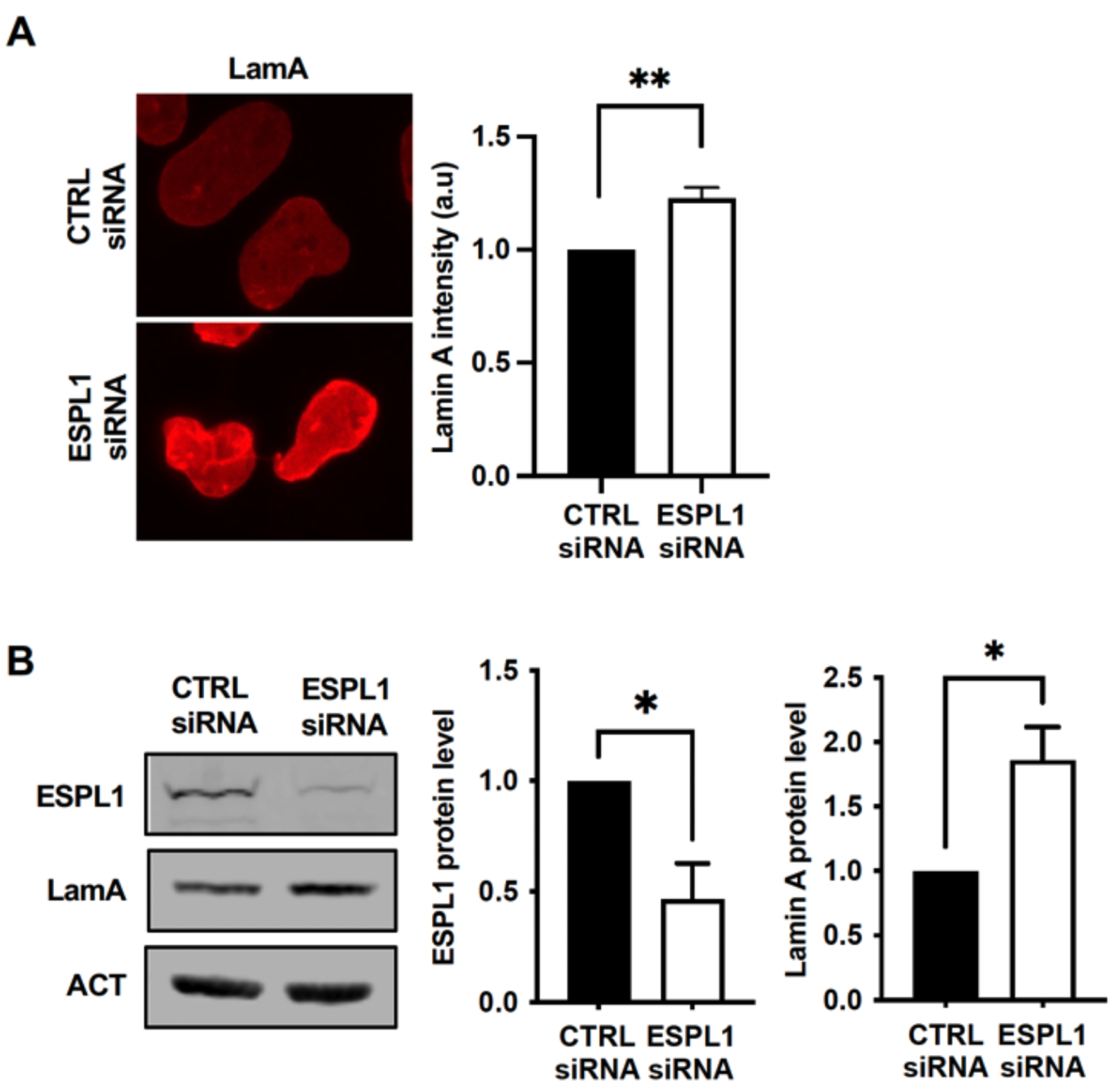
RNAi-mediated depletion of human separase (ESPL1) increases lamin A levels. (A) Immunofluorescent staining of lamin A following ESPL1 knockdown in human cells. SV40-fibroblasts transfected with scrambled siRNAs (CTRL) or ESPL1-specific siRNAs were stained with a lamin A-specific antibody. Representative images and quantification of lamin A intensity per nucleus (mean ± SEM from five independent experiments) are shown (**p<0.001; Student’s t-test). (B) Western blot analysis of lamin A and ESPL1 protein levels following ESPL1 knockdown in human SV40-fibroblasts. Cell lysates from fibroblasts transfected with either ESPL1-specific siRNA (ESPL1 siRNA) or control scrambled siRNA (CTRL siRNA) for 48 hours were probed with antibodies against lamin A, ESPL1, and ß-actin (loading control). Representative blots and quantification (mean ± SEM from three independent experiments) of lamin A and ESPL1 protein levels normalized to ß-actin are shown (*p<0.05; Student’s t-test).

Next, we investigated whether the interaction between A-type lamins and separase, identified in Drosophila, is also conserved in human cells. To this end, we performed in situ Proximity Ligation Assay (PLA) experiments in human SV40-fibroblasts and human primary fibroblasts. This analysis confirmed that human separase interacts with lamin A in situ, as they were detected in close proximity (<40 nm) in both primary and transformed fibroblasts (Figure 6A,B). Additionally, co-immunoprecipitation experiments in human SV40-fibroblasts using an anti-ESPL1 antibody reinforced that endogenous ESPL1 interacts with lamin A (Figure 6D). To rule out the possibility that the lamin A-ESPL1 complex—both of which are also DNA-binding proteins ^38, 39^—is bridged by DNA or RNA, we pretreated cellular lysates with benzonase nuclease, which degrades both nucleic acids, prior to immunoprecipitation. These experiments demonstrated that lamin A remains in complex with ESPL1 independently of DNA and RNA. Finally, reciprocal immunoprecipitation with an anti-lamin A antibody (Figure 6E) confirmed that nuclear A-type lamins form a stable complex with human separase.

**Figure 6.**
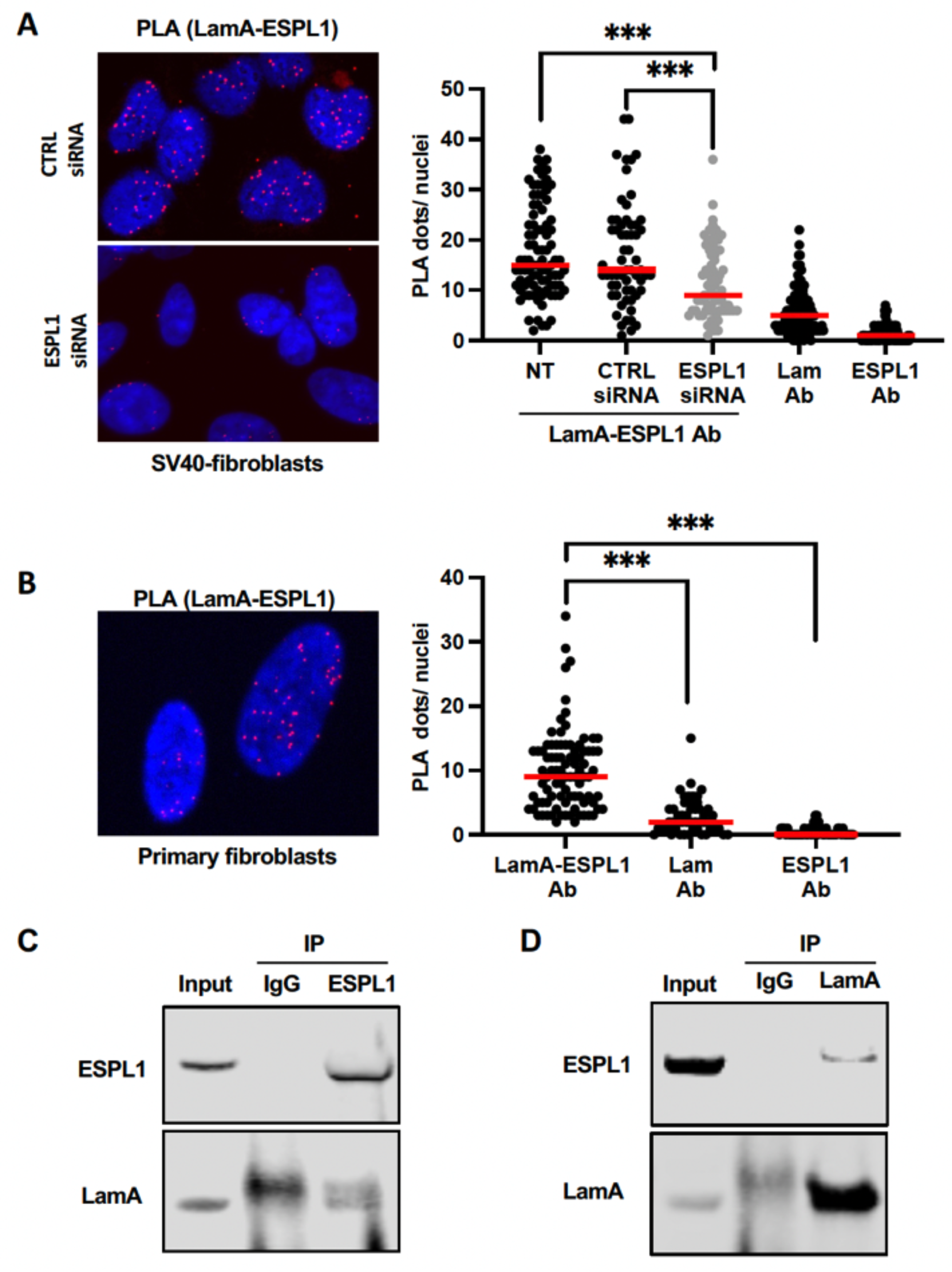
Separase interacts with lamin A in human cells. (A) Proximity ligation assay (PLA) using anti-lamin A (LamA) and anti-ESPL1 antibodies in human SV40-fibroblasts transfected with control siRNA (siCTRL) or ESPL1-specific siRNA (siESPL1). Non-transfected (NT) primary fibroblasts were also analyzed. Red dots indicate interactions between endogenous lamin A and ESPL1, with nuclei counterstained with DAPI (blue). Histograms depict the quantification of PLA dots per nucleus (***p<0.0001; Student’s t-test). Control PLA experiments were performed using a single primary antibody against either ESPL1 or lamin A alone. (B) Lamin A-ESPL1 PLA in human primary fibroblasts performed as described in (A). (C, D) Co-immunoprecipitation assays for endogenous human ESPL1 and lamin A. SV40-fibroblast lysates (1 mg) pre-treated with benzonase were immunoprecipitated with 5 µg of either mouse anti-ESPL1 (C) or mouse anti-lamin A (D) antibodies. Western blot analysis was performed on precipitates (IP) and 30 µg of input lysates using specific antibodies against ESPL1 and lamin A. A mouse IgG control (5 µg) was included (IgG).

Collectively, our findings indicate that the interaction between lamin and separase is conserved from Drosophila to humans.

## DISCUSSION

We demonstrate that separase interacts with lamins and colocalizes at the nuclear envelope in *Drosophila*, playing a crucial role in regulating lamin levels in both flies and human cells. Depletion of separase results in increased lamin A levels and nuclear shape alterations across both species. Our findings suggest that separase, beyond its established non-canonical functions, is a genuine lamin-interacting protein involved in nuclear lamina maintenance. This conclusion is also supported by a recent evidence of a physical interaction between Separase and Lamin B1 in human cells ^40^.

The observed effects of separase depletion on lamin regulation and nuclear morphology highlight its critical role in nuclear envelope integrity. Lamins are essential structural components of the nuclear lamina, and their misregulation has been linked to several pathological conditions, including laminopathies, premature aging disorders, and certain cancers. Our findings indicate that separase helps maintain nuclear homeostasis by modulating lamin levels, which in turn influences nuclear stiffness, chromatin organization, and gene expression. The fact that separase depletion leads to increased lamin A accumulation suggests a potential role in lamin turnover or degradation, possibly through direct cleavage or indirect regulation via proteasomal degradation pathways.

We also show that loss of separase affects nuclear morphology, inducing invaginations in *Drosophila* muscle cells, which subsequently leads to locomotion defects. Given that fruit flies serve as a model for human muscular dystrophies and laminopathies ^28, 41^, our findings highlight a conserved regulatory mechanism that may provide new insights into the aetiology of lamin-associated muscle diseases. Muscle cells are particularly susceptible to lamin dysfunction, as the nuclear envelope must endure significant mechanical stress. Thus, separase may therefore contribute to nuclear resilience by regulating lamin composition and ensuring proper nuclear mechanics.

Interestingly, we found that separase depletion has differential effects on Dm0 and LamC levels: it decreases Dm0 while increasing LamC. This suggests distinct regulatory mechanisms at play. The reduction in Dm0 appears to be transcriptionally mediated, likely due to the role of separase in facilitating transcription factor accessibility through regulation of the cohesin complex ^42^. This raises the possibility that separase positively regulates *LamDm0* gene expression by modulating chromatin structure.

Conversely, the increase in LamC levels does not result from higher mRNA expression, suggesting a post-translational mode of regulation. One intriguing possibility is that separase directly cleaves LamC, as the *Drosophila* LamC sequence contains multiple putative separase cleavage sites (EXXR). If this is the case, separase might function in LamC turnover and processing to ensure a balanced nuclear lamina composition. In support of this idea, we detected several LamC bands in Oregon R extracts that disappeared in *Sse* mutant extracts, which instead showed a significant accumulation of full-length LamC (∼75 kDa). However, our attempts to isolate LamC cleavage fragments by IP or detect them by IF were unsuccessful, leaving the precise nature of the role of separase in LamC regulation open for further investigation.

Our mass spectrometry (MS) analysis of *Sse^dft^* mutants confirmed the established roles of separase in cell cycle regulation and chromatin organization while uncovering an unprecedented function in *Drosophila* muscle homeostasis. While this could be an indirect consequence of lamin misregulation - given that lamins are critical for muscle development and nuclear integrity - it is also possible that separase has a more direct role in maintaining muscle function.

Notably, endopeptidase activity has been shown to be essential for generating micropeptides that regulate muscle contraction and heart function in *Drosophila* ^43^. Given that separase itself is an endopeptidase, it may participate in a similar process, potentially by processing key muscle proteins or micropeptides involved in contraction dynamics. This raises exciting possibilities regarding the role of separase in muscle physiology and suggests that its activity extends beyond mitotic regulation into post-mitotic tissues. Further biochemical and genetic studies will be necessary to explore this hypothesis.

Our findings reveal a novel regulatory axis linking separase, lamins, and nuclear organization, which could have broad implications for understanding nuclear architecture-related diseases. Given that separase is evolutionarily conserved, its role in lamin regulation may extend to vertebrates, including humans. Future research should explore whether separase mutations contribute to laminopathies or other nuclear envelope-related disorders.

## MATERIALS AND METHODS

### *Drosophila* strains, transgenic lines and crosses

The *Sse^dft^* mutant, the catalytic inactive separase UAS-6HA-Sse C497S expressing line (Dead Head, DH), the *dind*^2870^ and *dind*^1332^ mutant lines were described previously ^18, 31^. The *UAS Sse* RNAi (v29318), the *69B GAL4* and *Mef2 GAL4* drivers (which express GAL4 in brain and muscle, respectively) were obtained from the Vienna *Drosophila* stock center. To obtain flies expressing lamC with the N-terminal V5 and C-terminal Flag tags, synthetic V5lamCF coding sequence was cloned into pJZ4 vector (a pCASPER4 derivative) under the control of *Drosophila* tubulin promoter. Germline injection and transformation were carried out by the BestGene Company (Indiana, USA). To introduce recombinant V5-LamC-Flag encoding transgene in a *Sse^dft^* mutant genetic background, *w; pJZ4 w^+^V5lamCFlag/pJZ4 w^+^V5lamCFlag* (Ch.III) virgin females were crossed with *w; Sse^dft^/TM6B* males. *w; pJZ4 w^+^V5lamCFlag/ Sse^dft^* trans-heterozygous F1 females were crossed with *w; MKRS/TM6B* males. Single F2 recombinant males were then crossed with *w; MKRS/TM6B* virgin females to establish *w; w+pJZ4V5lamCFlag, Sse^dft^/TM6B* recombinant lines. *Sse^dft^* mutation was scored through homozygous late-lethality and chromosome endoreduplication/telomere fusions while *w^+^pJZ4V5lamCFlag* transgene was screened for *w^+^* phenotype. All strains were maintained on standard *Drosophila* cornmeal medium at 25 °C temperature.

### Real-time PCR

Third instar larvae RNA was extracted by using GENEzol Reagent (Geneaid, GZR100) following manual instructions as previously described ^44^. 1 μg of RNA for each sample was digested with 1 unit of DNase (DNase RNase-Free, Promega, M610C). RNA was reverse transcribed into cDNA with the 5x iScript™ Reverse Transcription Supermix kit (BIO-RAD, # 1708841) following manual protocol. Expression of lamins was evaluated through Real-Time PCR with the SsoAdvanced Universal SYBR Green Supermix kit (BIO-RAD) and using the following forward and reverse primers:

RP49 forward 5’-ATCGGTTACGGATCGAACAA-3’

RP49 reverse 5’-GACAAT CTCCTTGCGCTTCT-3’

Dm0 forward 5’-TTCGAGGAAACGCGGAAGAA-3’

Dm0 reverse 5’-TGTCCTCGTAGAGGGACTGG-3’

LamC RA forward 5’-GCTTGAACGCCAAGCTACAG-3’

LamC RA reverse 5’-GCAACATCCTCTTTGTCGGC-3’

LamC RB forward 5’ -TGGATCTCGAAATTGCCGCT-3’

LamC RB reverse 5’-CAAGTGCCGCATGTGTTGTAT-3’

### GST-pulldown and co-immunoprecipitation

To obtain GST-Sse and GST-Pim, bacterially expressed GST fusion proteins were purified with glutathione sepharose beads (Qiagen) from crude lysates by following manufacturer instructions. Protein extracts for GST-pulldown were obtained from third instar larval brains as previously described ^45^. Tissues were lysed in an ice-cold buffer containing 20mM Hepes KOH pH 7.9, 1.5mM MgCl2, 10mM KCl, 420mM NaCl, 20mM NaF, 10mM Na3VO4, 10mM BGP, 10mM phenylmethyl sulfonyl fluoride, 0.1% NP40 and 1 protease inhibitor cocktail (Roche). Protein extracts were incubated with each GST fusion protein bound to sepharose beads for 1h at 4 C in incubation buffer containing 20mM Hepes KOH, 20mM NaF and 0.8% NP40. After incubation Sepharose-bound GST proteins were collected and washed with a solution containing 20mM Hepes KOH, 20mM NaF and 0.8% NP40. For co-immunoprecipitation, whole cell extracts were obtained from human SV40-fibroblasts after lysing cells for one hour in a lysis buffer (50 mM Tris–HCl (pH 7.5), 150 mM NaCl, 5 mM EDTA, 0.5% NP40) supplemented with 1X complete protease inhibitor cocktail (Roche) and 1X phosphatase inhibitor cocktail (Roche), following by their centrifugation and the collection of supernatants containing protein extracts. Extracts were then treated with benzonase nuclease before co-immunoprecipitation using Dynabeads protein G kit (Life technologies). 5 µg of antibody raised against ESPL1 (Novus) or lamin A (Abcam) were incubated with Dynabeads for one hour at room temperature (RT). Normal mouse IgG (5 µg, Santa Cruz) was used as control. After washing with 0.05% Tween-20 in PBS, coupled beads were incubated with protein samples (0.5-1 mg) for 1.5 hour at RT. After washing, protein precipitates were eluted in Laemmli buffer (2X) with 4% ß-mercaptoethanol. Protein precipitates were further resolved by SDS-PAGE as described above using mouse anti-separase antibody (6H6, Novus Biological) or mouse anti-lamin A (Ab8980, Abcam). 60 µg of whole cell extracts were used as Input.

### Locomotion assays

Analysis of larval locomotion was performed on third instar larvae placed on a 2% agar substrate. The number of peristaltic movements for each larva over 1 minute was measured. Adult locomotion was evaluated through the Bang Sensitivity Test. After a mechanical stress, groups of 10 flies were pushed at the bottom of a cylinder and their ability to reach a distance of 4 cm in 5 seconds from the bottom was evaluated as the percentage of flies that covers the distance over the total flies’ number.

### Cell culture and transfection

Human SV40-transformed fibroblasts (GM0639) and normal human diploid embryonic fibroblast (WI-38) (Coriell Cell Repositories Camden, NJ) were cultured in Dulbecco’s modified Eagle’s medium (DMEM) supplemented with 10% fetal bovine serum (Gibco), 2mM-Glutamine, 200 U/ml penicillin and 200 mg/ml streptomycin (Sigma). For cell transfection, SV40-fibroblasts seeded on 6-plates wells (1.5 × 10^5^ cells/well) -containing coverslips for immunofluorescent experiments-for 24h were transfected with 20 nM siRNAs (smart-pool siRNAs designed against ESPL1 (Dharmacon) or a scrambled siRNA as control (Eurogenetec) using INTERFERin reagent (Polyplus). 48h after transfection, cells were collected for western blotting analysis or fixed for immunofluorescence staining or PLA experiments.

### *In situ* proximity ligation assay (PLA)

Cells plated on coverslips and fixed with ice-cold Methanol (10 min), were blocked one hour with 2% BSA/0,05% Tween 20 in PBS for 1 h, and incubated with a mouse primary antibody against ESPL1 (ab16170, Abcam) and a rabbit primary antibody against lamin A (1293, Sigma). Then, PLA was performed using the Duolink *in situ* PLA probes (anti-Mouse Minus and anti-Rabbit Plus, Sigma) and the Duolink *in situ* detection reagent red (Sigma) following the manufacturer’s instructions. Images were acquired with an SPE Leica DMRxA2 confocal microscope using a 63× objective lens and further analyzed with ImageJ software. Statistical analyses were performed with GraphPad PRISM (GraphPad Inc.) using unpaired 2-tailed Student t-tests.

### Cytology and immunofluorescence

For intact polytene nuclei preparation, third instar larvae salivary glands were dissected in a 0.7% NaCl and fixed in a drop of 1.8% formaldehyde and 40% acetic acid placed onto a cover slip. Samples were squashed gently and immediately frozen in liquid nitrogen. After flipping the coverslip, slides were raised in ice-cold Tris-buffered saline solution (TBS) for 10 minutes and subsequently permeabilized in TBS-Triton X-100 1% for 20 minutes. Slides were incubated with mouse anti-lamin Dm0 1:10 (ADL67.10-S Developmental Studies Hybridoma Bank), mouse anti-lamin C 1:10 (LC28.26-S Developmental Studies Hybridoma Bank), chicken anti-SSE 1:50 (16) and rabbit anti-V5 1:50 (abcam, ab15828) antibodies overnight at 4°C in a wet chamber. After washing with TBS-Triton 0.05%, salivary gland preparations were incubated with anti-mouse TRITC 1:200 (Jackson Immuno Research), anti-Chicken IgG (HþL) 1:200 (Alexa Fluor 488 ab150169) or anti-Rabbit IgG Alexa FluorTM Plus 594 1:200 (Invitrogen, # A 11012) for 1h at room temperature in a dark chamber. All slides were washed in TBS-Triton 0.05% and mounted with VECTASHIELD Antifade Mounting Medium with DAPI (Vector). Preparations were analyzed with a Zeiss AxioPlan epifluorescence microscope equipped with a cooled CCD camera (Photometrics). Images were pseudocoloured and merged with AdobePhotoshop CS4. For apoptosis analysis, third instar larvae were exposed to 10 γ rays from 137Cs sources using the Gammacell Exactor 40 (Nordion). At 8h post irradiation, larval brains were dissected and processed for anti-cleaved *Drosophila* Dcp1 staining as previously described ^46^. For larval muscle fillets preparation, third instar larvae were dissected in PBS1x and fixed in a solution containing 4% paraformaldehyde and 4% sucrose for 1 hour at room temperature. Tissues were washed 3 times with PBS and permeabilized with PBS-Triton 1% for 10 minutes. After 2 two washes, the preparations were incubated with mouse anti-lamin C 1:10 (LC28.26-S Developmental Studies Hybridoma Bank) O.N. at 4 °C in a humid chamber. After two washes in PBS, the fillets were incubated with Alexa Fluor phalloidin 594 1:300 (Thermo Fischer Scientific) and with anti-mouse FITC 1:200 (Jackson Immuno Research) in PBS-Triton 0.1% BSA 2% for 30 minutes in the dark at room temperature. The fillets were washed 3 times and mounted with VECTASHIELD Antifade Mounting Medium with DAPI (Vector).

For human fibroblasts staining, cells plated on coverslips and fixed in ice-cold Methanol (10 min) 48h after transfection, were permeabilized with 0,5% Triton X-100 in PBS for 10 min, then blocked with 2% BSA/0,05% Tween 20 in PBS for 1 h. After the blocking step, cells were incubated with a mouse primary antibody against lamin A (ab8980, Abcam) for 1 h at room temperature. Cells were then incubated with an anti-mouse IgG AlexaFluor 488 (Life Technologies). DNA was counterstained with DAPI and coverslips were mounted onto slide with fluoromount mounting medium (Southern Biotech). Images were acquired using a leica DM5500B fluorescence microscope with a 63×-oil objective. Following acquisition, images were analyzed with ImageJ software. Statistical analyses were performed with PRISM using unpaired t-tests.

### Western-blot

*Drosophila* third instar larval brains were homogenized in SDS buffer (10 mM Tris (pH 7.5), 1% SDS and 1X complete protease inhibitor cocktail (Roche). Protein samples were resolved by SDS-polyacrylamide gel electrophoresis and blotted onto nitrocellulose membrane (Amersham). Membranes were blocked in 5% skim milk in PBS/0.1% Tween-20 for 45 min then probed with the indicated primary antibodies: mouse anti-lamin Dm0 1:500 (ADL67.10-S Developmental Studies Hybridoma Bank), mouse anti-lamin C 1:500 (LC28.26-S Developmental Studies Hybridoma Bank), anti-actin HRP conjugated (1:5,000; Santa Cruz Biotechnology SC-1615), anti hts (AB528289 Developmental Studies Hybridoma Bank), anti-prc (EC11 Developmental Studies Hybridoma Bank), anti-FLAG HRP conjugated (1:3,000; Sigma A8592), anti-Tubulin (1:100,000; Sigma T6199), anti-V5 HRP conjugated (1:4000; abcam 1325), anti-Giotto (1:5000; ^47^). Primary antibodies were detected using sheep anti-mouse IgG HRP conjugated (1:5,000 NA931V Amersham Biosciences) and donkey anti-rabbit IgG HRP conjugated (1:5,000 NA934 Amersham Biosciences).

Human fibroblasts cells were homogenized in SDS buffer (10 mM Tris (pH 7.5), 1% SDS, 1X complete protease inhibitor cocktail (Roche) and phosphatase inhibitors cocktail 2 and 3 (Sigma). Protein samples were resolved by SDS-polyacrylamide gel electrophoresis on 4–12% NuPAGE Bis-Tris gradient gel or 3–8% Tris-acetate gel (Invitrogen) with MOPS or Tris-acetate running buffer (Invitrogen) and then, blotted onto on nitrocellulose membrane (Amersham). Membranes were blocked in 5% skim milk in PBS/0.1% Tween-20 for 45 min then probed with the indicated primary antibodies: ESPL1 (6H6, Novus Biologicals), lamin A (ab8980, Abam) and ß-actin (A2066, Sigma) as loading control. Primary antibodies were further detected using secondary fluorescent antibody (IR800 and IR700, Diagomics). Fluorescent signals were acquired with Odyssey imager (LI-COR Biosciences) and quantification were performed with Image Studio software (LI-COR Biosciences) or Image J software.

All WB images were assembled using the sections highlighted in the uncropped blots included in the Supplementary Figures.

### Mass spectrometry

Proteins extract from third instar larvae brains were obtained through lysis in a solution containing 8 M Urea, 20 mM Hepes pH 8, 1mM PMSF and protease inhibitors cocktail (Roche). Samples were digested with trypsin followed by isobaric Tag for Relative and Absolute Quantitation (iTRAQ) labeling. After high pH reversed-phase liquid chromatography (HP-RPLC) fractioning peptides were subjected to tandem Mass Spectrometry (Proteomics and Mass Spectrometry Facility, BRC, Cornell University). Proteins with at least 2 PSMs were considered for differential expression analysis. Differentially expressed proteins were detected with the *t-test* and multiple tests adjusted *p-values* were calculated with the Benjamini-Hochberg method. Gene Ontology (GO) enrichment analyses were performed using the WEB-based Gene SeT AnaLysis Toolkit (WebGestalt) ^48^ over Biological Process, Molecular function and Cellular Component GO categories, with the Over-Representation Analysis (ORA) method based on the Hypergeometric test (Benjamini-Hochberg multiple test correction, FDR threshold: < 0.05). All WebGestalt-based analyses used the protein coding genome for *Drosophila melanogaster* as a background reference set. Enrichment ratio parameter was calculated as the number of observed entries divided by the number of expected entries from each GO category. Metabolic pathways and domains enrichment analyses were performed using the FlyEnrichr tool ^49^ on KEGG (2019) and InterPro (2019) databases, calculating *p-values* with the hypergeometric tests based-methods. A rank score (or *z-score*) is calculated to weigh the deviation from an expected rank. After functional enrichment analysis, a combined score is given, which represents a combination of *p-value* and *z-score* in the form of c = ln(p)*z. The STRING tool ^50^ was used to investigate proteins networks among enriched GO categories. k-means algorithms were applied on the network to consider the distance matrix computed by STRING. Statistical analyses and plots and consensus were computed using R software.

### Statistical analysis

Statistical analyses were performed using GraphPad PRISM (GraphPad Inc.). Statistical significance of data was assessed by unpaired 2-tailed Student t tests. P > 0.05 was considered not significant.

## Acknowledgements

This work was supported by a grant from AFM-Téléthon to G.C. and P.B (N.21566), by Italy University and Research Ministry (MUR, N. 202227SYBW) and Istituto Pasteur Italia, Fondazione Cenci Bolognetti (Anna Tramontano Grant) to GC, and by a grant from Italy Ministry of University and Research (PRIN, N 2022KKMTL9) to F.C. P.B. ‘s laboratory is also supported by the Ligue Nationale Contre le Cancer (Haut-de-Seine committee), Association for Research against Cancer (Fondation ARC), AT Europe Association, INCA grant (PLBIO23-204-2023-179), Emergence Paris Cité, CEA Radiobiology Program and INSERM, Université Paris Cité, Université Paris-Saclay house funding (SGCSR unit)

## Authors contributions

F. C. designed and performed all the experiments on Drosophila, assembled the corresponding figures and wrote the manuscript. G.P. designed and performed all the experiments on human cells, designed the corresponding figures and wrote parts of the manuscript. G.B., L.T., S. N., and C. B. performed experiments on Drosophila. S. R. and A. P. analyzed the MS data and assembled the corresponding figures and data. MLG, designed and supervised the MS experiments. P.B. designed the experiments, supervised, and re-viewed the manuscript. G.C. designed and supervised the experiments, and wrote the manuscript.

