## Supplemental figures for "An evolutionarily conserved role for separase in the regulation of nuclear lamins"

### Supplementary Figures

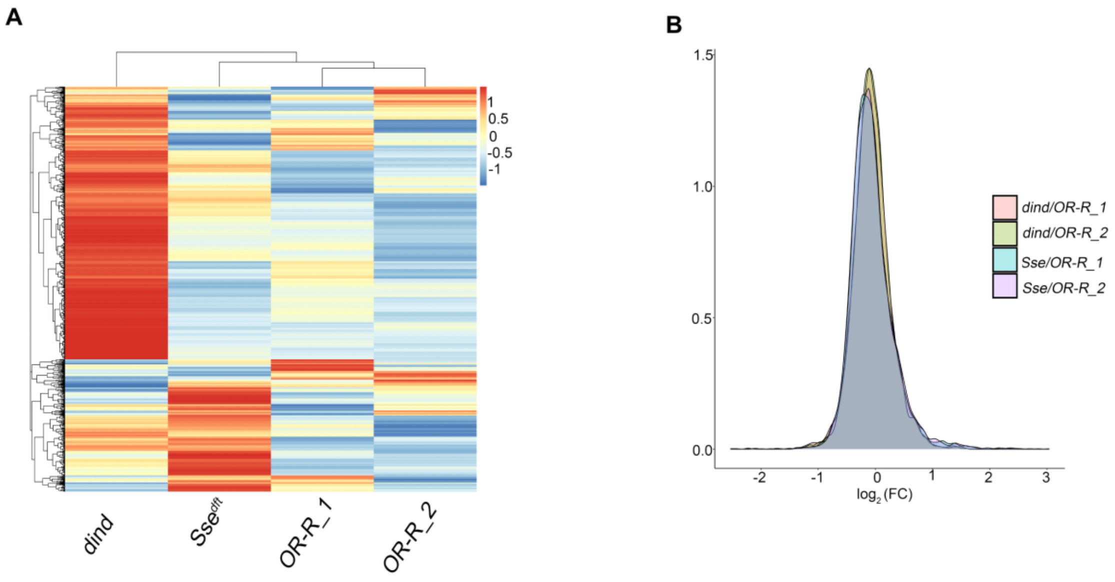


**Supplementary Figure 1**: A) Heatmap showing the $\log_2$ transformed abundance ratio values for the differentially expressed proteins in control (*OR-R_1* and *OR-R_2*) and in *diamond (dind)* and *Separase (Sse)* mutant extracts. B) $\log_2$(*FC*) values distribution. Dashed lines represent the thresholds of 0.4 and -0.4.

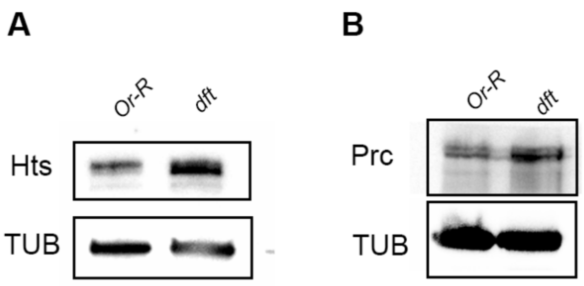


**Supplementary Figure 2**. WB from *Sse*$^{dft}$ mutant extracts *(dft)* showing that, consistently with our MS data (Table 1), Hts (A) and Prc (B) are upregulated upon the loss of Separase. Anti-Tubulin has been used as a loading control.

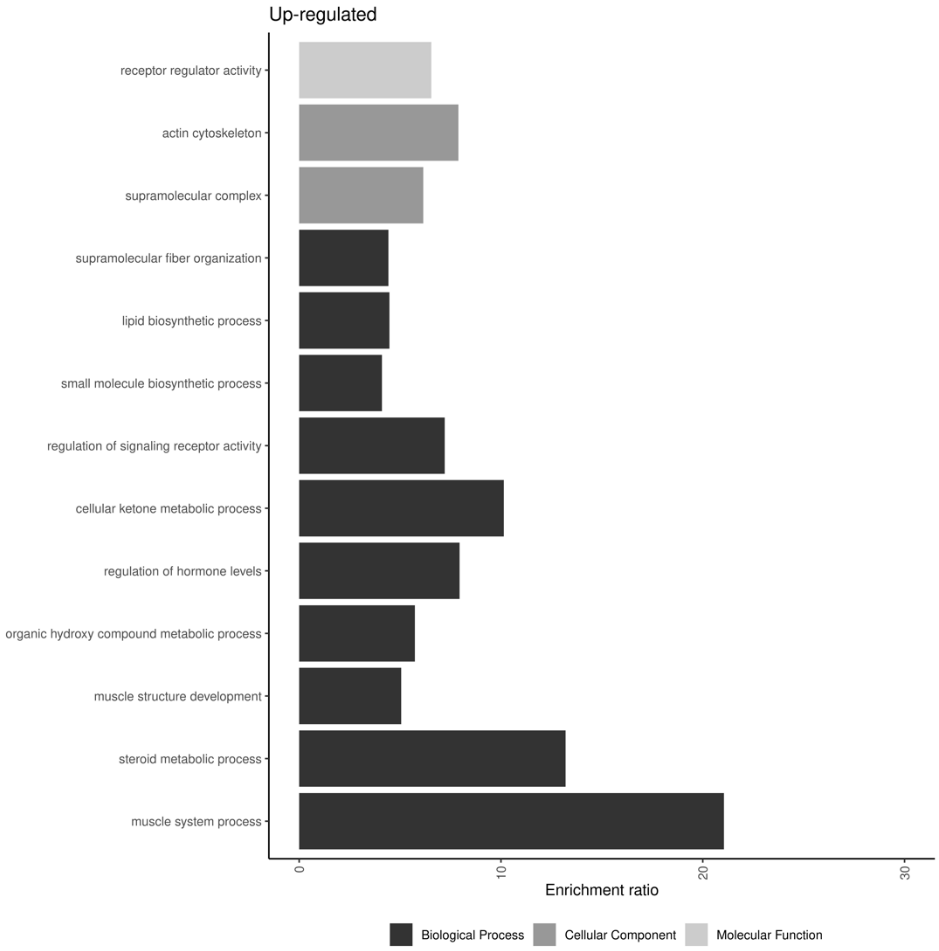


**Supplementary Figure 3**: Bar plot showing GO enriched categories of proteins up-regulated in the *Sse*$^{dft}$ mutant extracts after the elimination in the mitotic blockage mutant. All the enriched category showed an FDR < 0.05.

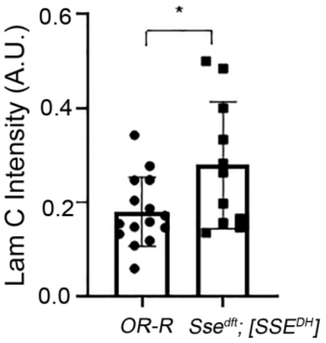


**Supplementary Figure 4**

Quantification of LamC fluorescence intensity from salivary gland immunostaining from control third instar larvae (Or-R) and *Sse*$^{dft}$ mutant larvae expressing a catalytically inactive Separase (SSE$^{DH}$). At least three slides were used for the quantification (*p<0.05; Student's t-test).

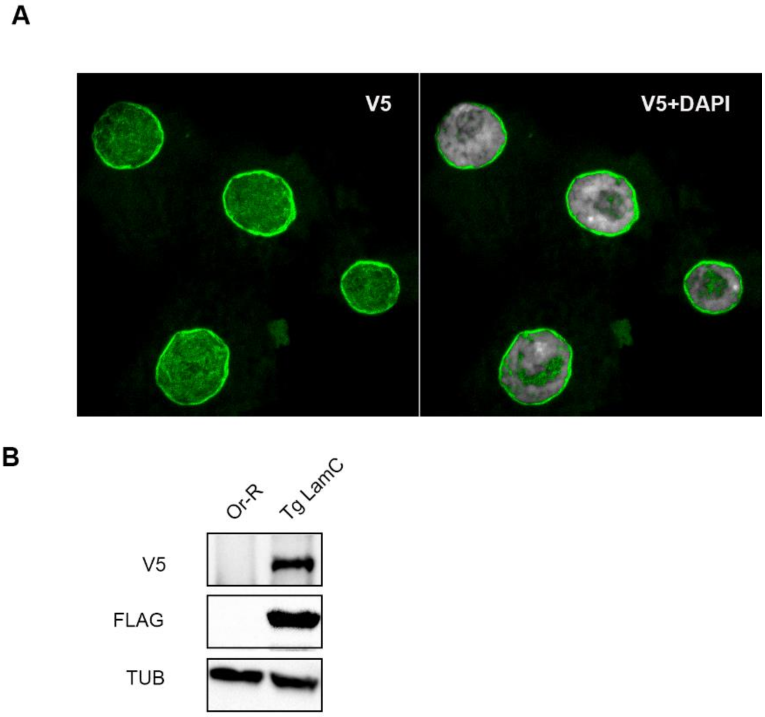


**Supplementary Figure 5.** Immunostaining (A) and WB (B) on larvae expressing the V5-LamC-FLAG encoding transgenes (Tg LamC) with either a commercial anti-V5 (A and B) or anti-FLAG (B) antibody. Note the perinuclear pattern of the recombinant protein in A indicating that both TAGs did not affect Lamin C expected localization.

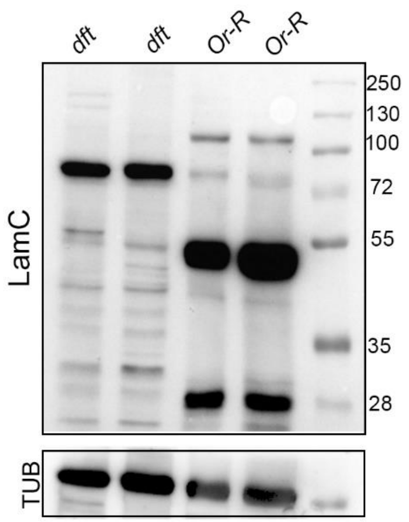


**Supplementary figure 6.** WB on *Sse*$^{dft}$ mutant and *OR-R* (control) larval brain extracts providing evidence of a potential SSE dependent cleavage of Lamin C that in normal conditions generates two prevalent Lam C fragments of ~50 kDa and ~25 kDa as well as a less abundant full length Lam C (~75 kDa). The cleavage is missing when Separase is lost with only the ~75 kDa full length Lamin C observable in *Sse*$^{dft}$ mutant extracts. It is worth mentioning that the presence of the two bands in control extract is consistent with the localization of one EXXR potential Separase cleavage site at position 401. Interestingly, the EXXR cleavage site at position 398-401 is also conserved in human LMNA (position 382-386) and its mutation is found in some AD-EDMD patients (R386K) (Scharner et al., 2011). We would like to mention that, for some still unclear reasons, the LamC pattern shown in this figure was not always as evident as it in all our experiments. Thus, although we are still convinced that lamin C is cleaved in physiological conditions, because of this unknown variability, we decided not to include and consider these results in our manuscript. However, we wish to show this pattern as a supplementary information hoping that this finding could provide a useful hint for all interested readers who are willing to successfully address this potential cleavage.

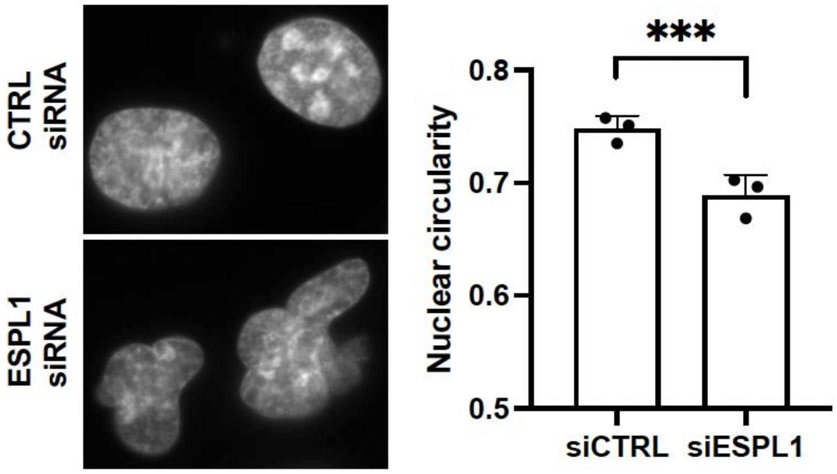


**Supplementary Figure 7**. DAPI staining of human fibroblast nuclei transfected with either scrambled siRNA (CTRL) or ESPL1 siRNA. Note the presence of misshapen nuclei upon ESPL1 depletion. On the right, quantification of nuclear circularity from control and ESPL1 depleted cells of 3 independent experiments (*** p<0.001).

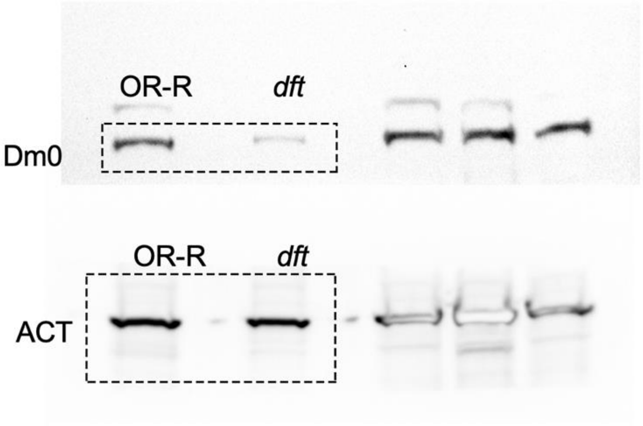


**Uncropped Figure 2 A**

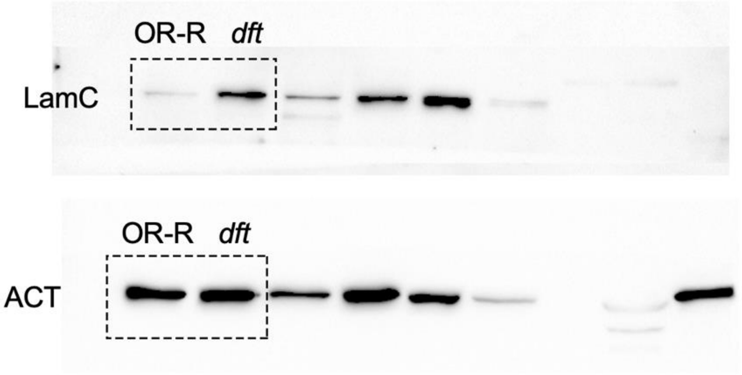


**Uncropped Figure 2 B**

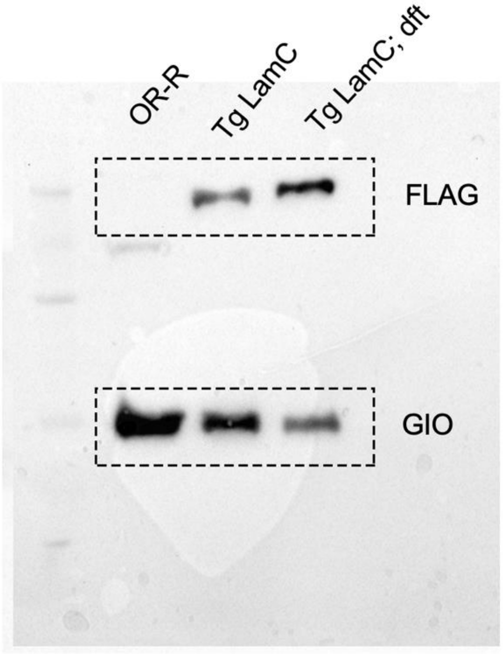


**Uncropped Figure 2 G**

**Supplementary Figure 8.** Uncropped blots used for Figures 2A, 2B and 2 G. Rectangles refer to the sections used for panels of Figure 3.

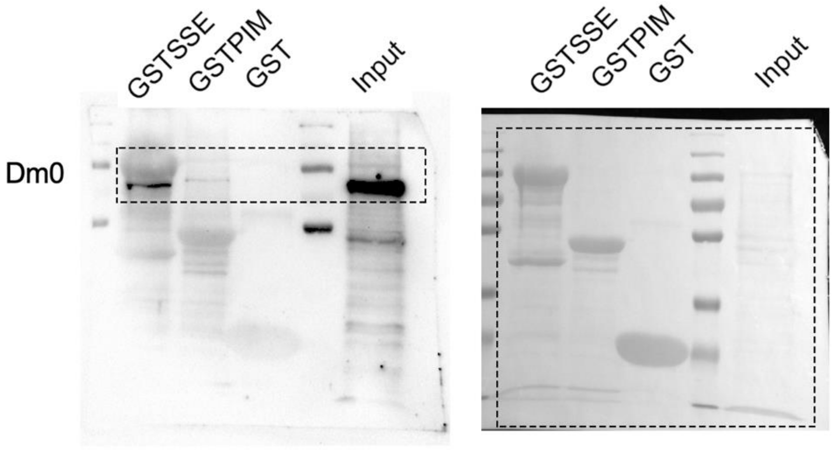


**Uncropped Figure 3 A**

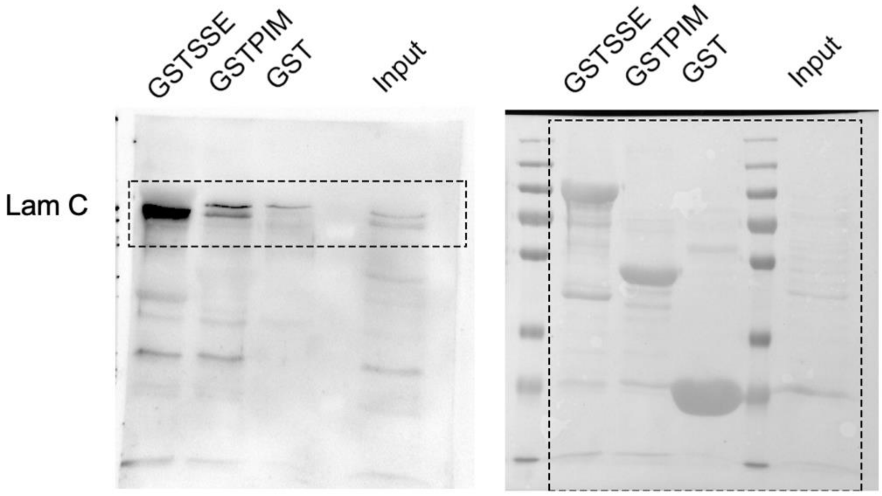


**Uncropped Figure 3 B**

**Supplementary Figure 9.** Uncropped blots used for Figures 3A and 3B. WBs and Ponceau are shown in the left and in the right, respectively. Rectangles refer to the sections used for panels of Figure 3.

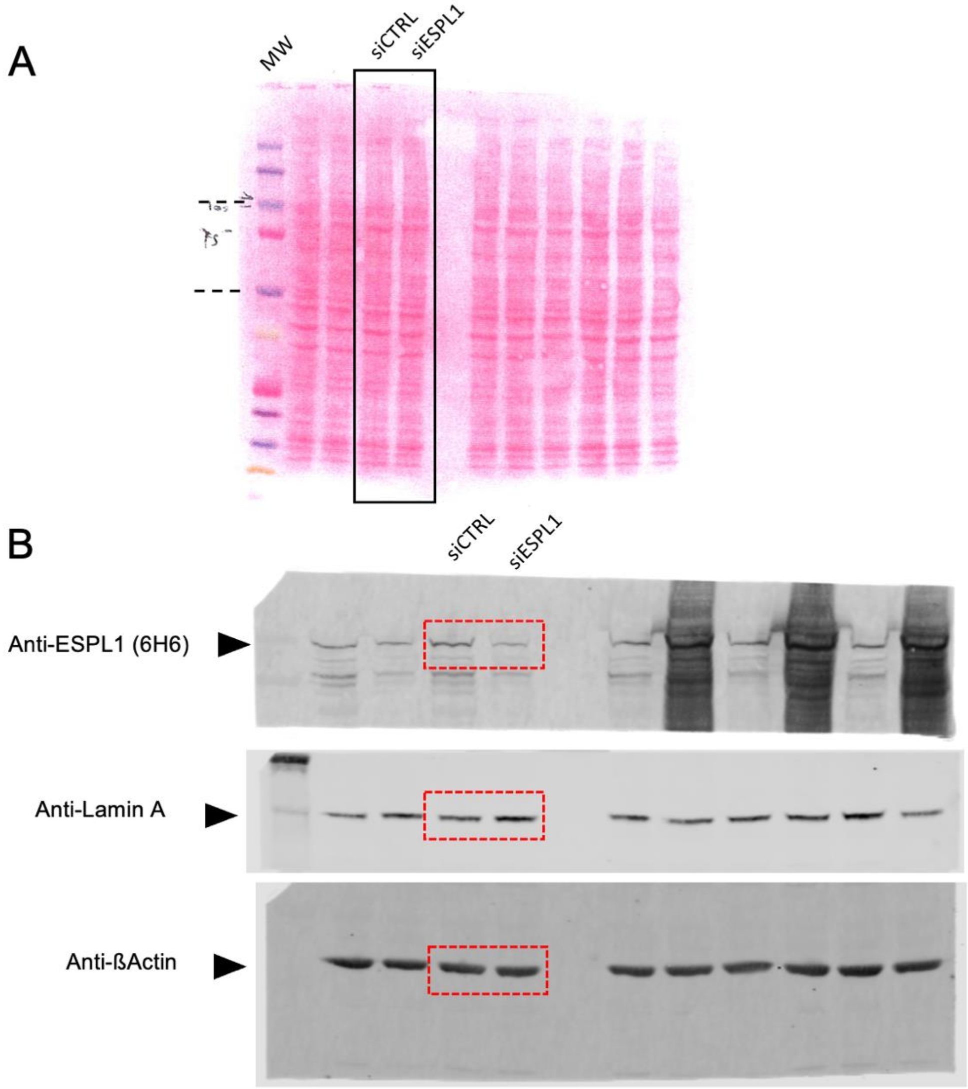


**Supplementary Figure 10.** Uncropped scan of WB shown in Figure 5B. A. Ponceau S- stained membrane immunoblotted in Figure 5B is shown. MW: Kaleidocope (Biorad). B. Scans of immunoblots shown in Figure 5B. Membrane was cutted in 3 parts according to molecular weight (> 100 kda; 50-100 kda and < 50 Kda) (dashed line indicates cut locations) and hybridized with the following antibodies, respectively: anti-ESPL1, anti-lamin A and anti-ßactin. Red rectangles referring to the sections used for panels of Figure 5B.

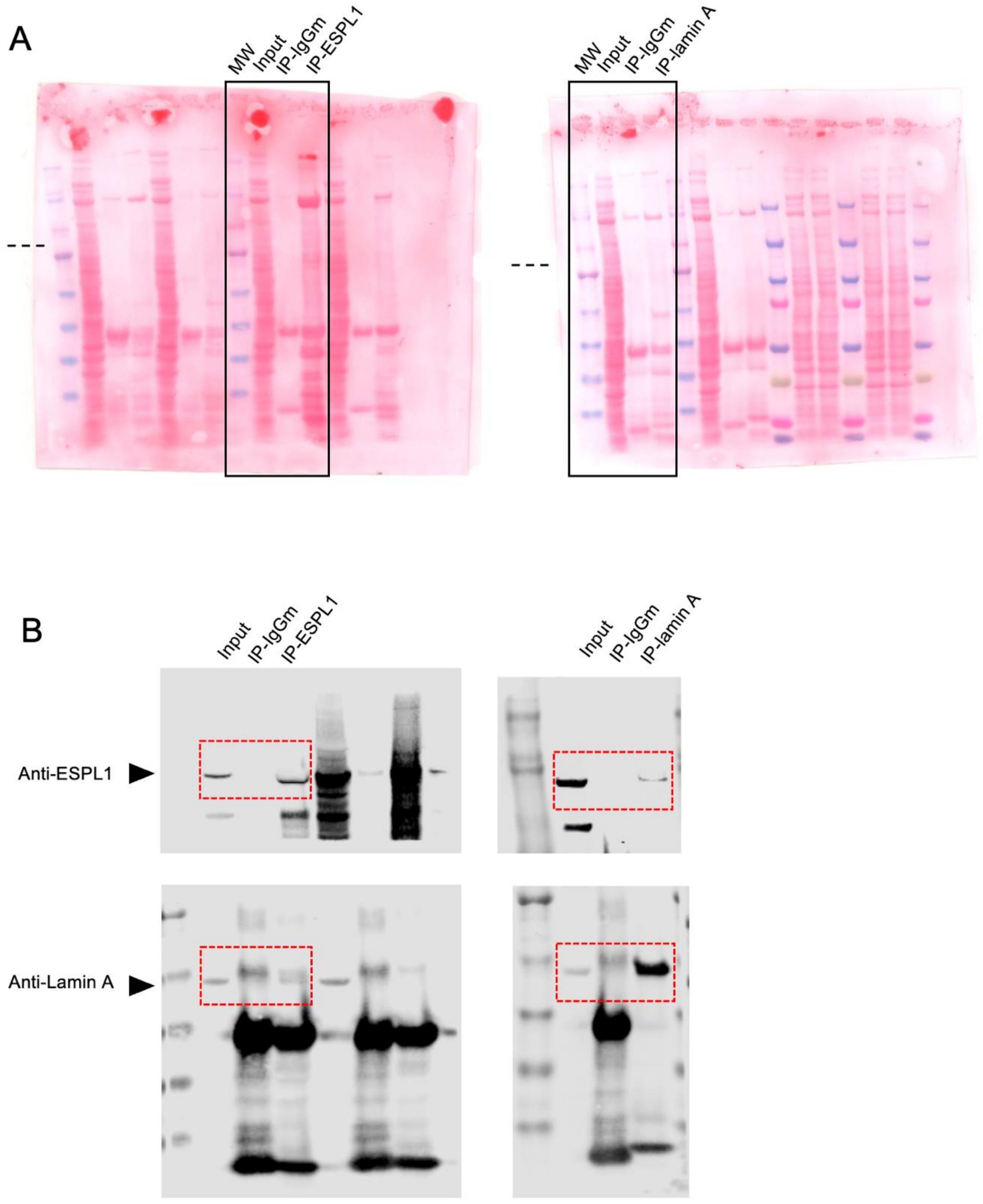


**Supplementary Figure 11**. Uncropped scan of WB shown in Figure 6C and D. A. Ponceau S-stained membranes immunoblotted in Figure 6C (left) and D (right) are shown;  MW= High Mark (Invitrogen). B. Membranes were cutted in 2 parts according to molecular weight (≥ 117 kda and ≤ 117 Kda) and hybridized with the following antibodies anti-ESPL1 and anti-lamin A ; dashed line indicates cut locations. Red rectangles refer to the sections used for panels of Figure 6C and D.
