## Supplemental Table 1 for "An evolutionarily conserved role for separase in the regulation of nuclear lamins"

| Accession | Description | Abundance.Ratio…Sse…Cntr | Abundance.Ratio…Sse…Cnt |
|---|---|---|---|
| Q9VWD3 | CG12531, isoform A OS=Drosophila melanogaster GN=CG12531 PE=1 SV=2 | 0.401630466584741 | 0.410341104614087 |
| Q9VBP6 | GH21316p OS=Drosophila melanogaster GN=Ssadh PE=1 SV=1 | 0.40381306161659 | 0.441483479582645 |
| A0A0B4KFE4 | Acyl-CoA synthetase long-chain, isoform J OS=Drosophila melanogaster GN=Acsl PE=1 SV=1 | 0.407080775450501 | 0.423309237242387 |
| Q9VF53 | CG18522 OS=Drosophila melanogaster GN=AOX1 PE=1 SV=1 | 0.407080775450501 | 0.415758666522498 |
| D3DML7 | MIP14691p OS=Drosophila melanogaster GN=sky PE=1 SV=1 | 0.408168370708106 | 0.404903122145131 |
| Q7K4C7 | CTP:phosphocholine cytidylyltransferase 1, isoform A OS=Drosophila melanogaster GN=Cct1 PE=1 SV=1 | 0.408168370708106 | 0.424384671538097 |
| O97061 | Ccp84Ad OS=Drosophila melanogaster GN=Ccp84Ad PE=4 SV=1 | 0.409255146684838 | 0.492622328574446 |
| Q9W4N6 | CG6428 OS=Drosophila melanogaster GN=CG6428 PE=1 SV=1 | 0.409255146684838 | 0.426533138116673 |
| Q9W247 | CG4752 OS=Drosophila melanogaster GN=CG4752-RA PE=1 SV=2 | 0.411426245726465 | 0.526068811667588 |
| M9PF67 | CG34356, isoform F OS=Drosophila melanogaster GN=CG34356 PE=4 SV=1 | 0.411426245726465 | 0.425459304765355 |
| Q9VWV5 | CG32549, isoform G OS=Drosophila melanogaster GN=CG32549 PE=1 SV=3 | 0.412510571249805 | 0.421155960662223 |
| Q95029 | Cathepsin L OS=Drosophila melanogaster GN=Cp1 PE=2 SV=2 | 0.412510571249805 | 0.459431618637297 |
| Q9VGA0 | Glutathione S transferase D9, isoform A OS=Drosophila melanogaster GN=GstD9 PE=1 SV=1 | 0.412510571249805 | 0.444667066841908 |
| Q9VDC3 | Adaptor protein complex 2, sigma subunit OS=Drosophila melanogaster GN=AP-2sigma PE=1 SV=1 | 0.412510571249805 | 0.440420720756767 |
| Q9VG97 | Inactive glutathione S-transferase D3 OS=Drosophila melanogaster GN=GstD3 PE=2 SV=1 | 0.413594082409175 | 0.570462931026041 |
| Q9VAG9 | CG7789 OS=Drosophila melanogaster GN=CG7789 PE=1 SV=1 | 0.417920007811965 | 0.421155960662223 |
| M9PHG3 | Diacylglycerol kinase OS=Drosophila melanogaster GN=rdgA PE=3 SV=1 | 0.418999465431266 | 0.535057594894842 |
| Q9VI25 | Neurochondrin homolog OS=Drosophila melanogaster GN=Neurochondrin PE=2 SV=1 | 0.421155960662223 | 0.402722176845605 |
| Q9VGU7 | CG14696 OS=Drosophila melanogaster GN=CG14696 PE=1 SV=1 | 0.422233000683048 | 0.483364360713349 |
| Q9I7D0 | GH22314p OS=Drosophila melanogaster GN=PhKgamma PE=2 SV=3 | 0.424384671538097 | 0.402722176845605 |
| Q960D4 | CG4199, isoform B OS=Drosophila melanogaster GN=EG:22E5.5 PE=1 SV=1 | 0.426533138116673 | 0.400537929583729 |
| P22979 | Heat shock protein 67B3 OS=Drosophila melanogaster GN=Hsp67Bc PE=2 SV=2 | 0.426533138116673 | 0.485426827170242 |
| P25455 | 1-phosphatidylinositol 4,5-bisphosphate phosphodiesterase classes I and II OS=Drosophila melanogaster GN=Plc21C PE=2 SV=3 | 0.42867840994823 | 0.435095151620097 |
| Q7K511 | CG3835, isoform A OS=Drosophila melanogaster GN=D2hgdh PE=1 SV=1 | 0.429749850800216 | 0.469885976274464 |
| Q9VIF2 | CG9248, isoform A OS=Drosophila melanogaster GN=CG9248 PE=1 SV=1 | 0.429749850800216 | 0.482332020747376 |
| B7Z107 | CG42376, isoform A OS=Drosophila melanogaster GN=cin-RB PE=2 SV=1 | 0.431890348286181 | 0.473007567916174 |
| Q9W543 | Rabconnectin-3B, isoform A OS=Drosophila melanogaster GN=Rbcn-3B PE=1 SV=1 | 0.432959407276106 | 0.454175893185802 |
| P54611 | V-type proton ATPase subunit E OS=Drosophila melanogaster GN=Vha26 PE=2 SV=1 | 0.434027674663597 | 0.404903122145131 |
| Q9VHF9 | CG8861, isoform B OS=Drosophila melanogaster GN=CG8861 PE=4 SV=3 | 0.43616183931445 | 0.543000877402426 |
| Q8SY19 | CG1742-PA, isoform A OS=Drosophila melanogaster GN=Mgstl PE=1 SV=1 | 0.43616183931445 | 0.451013242943973 |
| Q7KN85 | ATP-citrate synthase OS=Drosophila melanogaster GN=ATPCL PE=1 SV=1 | 0.438292851579147 | 0.408168370708106 |
| Q9VIH9 | CG9338, isoform B OS=Drosophila melanogaster GN=CG9338 PE=1 SV=1 | 0.438292851579147 | 0.43722773891291 |
| Q9VCF2 | Diacylglycerol kinase OS=Drosophila melanogaster GN=CG31140 PE=1 SV=2 | 0.439357178474257 | 0.420078115979374 |
| Q9W2V2 | CG32683, isoform A OS=Drosophila melanogaster GN=CG32683-RA PE=1 SV=2 | 0.439357178474257 | 0.504874589398464 |
| P48596 | GTP cyclohydrolase 1 OS=Drosophila melanogaster GN=Pu PE=2 SV=3 | 0.439357178474257 | 0.422233000683048 |
| A1ZA86 | CG8401 OS=Drosophila melanogaster GN=CG8401 PE=4 SV=2 | 0.439357178474257 | 0.531069492725954 |

| | | | |
|---|---|---|---|
| Q9V8R9 | Protein 4.1 homolog OS=Drosophila melanogaster GN=cora PE=1 SV=1 | 0.440420720756767 | 0.457331625485014 |
| Q8MRM6 | CG12065, isoform C OS=Drosophila melanogaster GN=CG12065 PE=1 SV=1 | 0.441483479582645 | 0.409255146684838 |
| Q9Y166 | BcDNA.GH02431 OS=Drosophila melanogaster GN=Dic1 PE=1 SV=1 | 0.441483479582645 | 0.509949146304311 |
| P07668 | Choline O-acetyltransferase OS=Drosophila melanogaster GN=Cha PE=1 SV=3 | 0.442545456105304 | 0.592636428606577 |
| Q9VBS0 | CHKov2 OS=Drosophila melanogaster GN=CHKov2 PE=1 SV=2 | 0.442545456105304 | 0.554834395894193 |
| Q9W425 | Rabconnectin-3A OS=Drosophila melanogaster GN=Rbcn-3A PE=1 SV=3 | 0.444667066841908 | 0.404903122145131 |
| A0A0B4K7U4 | Mbl, isoform H OS=Drosophila melanogaster GN=mbl PE=4 SV=1 | 0.444667066841908 | 0.58207422139633 |
| Q9VM58 | CG10399, isoform A OS=Drosophila melanogaster GN=CG10399 PE=1 SV=2 | 0.445726703349984 | 0.55581615506164 |
| A0A0B4JDC9 | RIM-binding protein, isoform F OS=Drosophila melanogaster GN=Rbp PE=1 SV=1 | 0.446785562143122 | 0.51500591643373 |
| M9PE01 | Ecdysone-induced protein 63E, isoform N OS=Drosophila melanogaster GN=Eip63E PE=4 SV=1 | 0.446785562143122 | 0.479230561206336 |
| M9PDX2 | CG4577, isoform B OS=Drosophila melanogaster GN=CG4577 PE=4 SV=1 | 0.447843644362086 | 0.414676780426886 |
| A0A0B4JCY6 | Cheerio, isoform I OS=Drosophila melanogaster GN=cher PE=1 SV=1 | 0.448900951145128 | 0.488515008957812 |
| Q7KTW5 | CG9391, isoform C OS=Drosophila melanogaster GN=CG9391 PE=1 SV=1 | 0.448900951145128 | 0.468843942974637 |
| Q8INU6 | CG31674, isoform B OS=Drosophila melanogaster GN=CG31674-RA PE=1 SV=1 | 0.448900951145128 | 0.620117165028684 |
| Q9VUY9 | Phosphoglucomutase OS=Drosophila melanogaster GN=Pgm PE=1 SV=1 | 0.451013242943973 | 0.431890348286181 |
| Q9VSF2 | Mediator of RNA polymerase II transcription subunit 24 OS=Drosophila melanogaster GN=MED24 PE=1 SV=2 | 0.451013242943973 | 0.413594082409175 |
| Q9VHC3 | Blistery, isoform A OS=Drosophila melanogaster GN=by PE=2 SV=1 | 0.453122446595813 | 0.558757430373762 |
| B7Z001 | CG3523, isoform C OS=Drosophila melanogaster GN=FASN1 PE=1 SV=2 | 0.455228571117139 | 0.445726703349984 |
| Q9W293 | CG6613 OS=Drosophila melanogaster GN=CG6613 PE=1 SV=1 | 0.455228571117139 | 0.457331625485014 |
| Q95NU8 | GH16255p OS=Drosophila melanogaster GN=jeb PE=2 SV=1 | 0.459431618637297 | 0.506906554580693 |
| E1JIH3 | Unc-115a, isoform B OS=Drosophila melanogaster GN=Unc-115a PE=1 SV=1 | 0.460480470040011 | 0.435095151620097 |
| Q9VPG2 | CG13248 OS=Drosophila melanogaster GN=CG13248-RA PE=2 SV=1 | 0.460480470040011 | 0.430820496519772 |
| O97064 | Ccp84Ag OS=Drosophila melanogaster GN=Ccp84Ag PE=1 SV=1 | 0.461528559472877 | 0.562669826102702 |
| Q5U191 | CG1882, isoform A OS=Drosophila melanogaster GN=CG1882 PE=1 SV=1 | 0.461528559472877 | 0.551885103471725 |
| H9XVM3 | ADP ribosylation factor-like 4, isoform B OS=Drosophila melanogaster GN=Arl4 PE=4 SV=1 | 0.462575888042204 | 0.488515008957812 |
| Q86PA0 | CG17816, isoform D OS=Drosophila melanogaster GN=CG17816 PE=1 SV=1 | 0.464668267003444 | 0.418999465431266 |
| A8JUZ7 | CG15894, isoform B OS=Drosophila melanogaster GN=CG15894 PE=1 SV=1 | 0.464668267003444 | 0.586884812852185 |
| Q86BM0 | CG9515, isoform B OS=Drosophila melanogaster GN=CG9515 PE=1 SV=1 | 0.468843942974637 | 0.453122446595813 |
| Q7K1W4 | Juvenile hormone epoxide hydrolase 3 OS=Drosophila melanogaster GN=Jheh3 PE=1 SV=1 | 0.469885976274464 | 0.486456955768731 |
| Q9VB69 | Malic enzyme OS=Drosophila melanogaster GN=Men-b PE=1 SV=1 | 0.470927257475127 | 0.486456955768731 |
| M9PDQ9 | CG31974, isoform D OS=Drosophila melanogaster GN=CG31974 PE=1 SV=1 | 0.470927257475127 | 0.431890348286181 |
| Q9VZI3 | Unc-112-related protein OS=Drosophila melanogaster GN=Fit1 PE=1 SV=1 | 0.474046599319306 | 0.486456955768731 |
| Q9V4M2 | Protein wech OS=Drosophila melanogaster GN=wech PE=1 SV=2 | 0.475084882948783 | 0.466757615726171 |
| O46100 | CG12773, isoform A OS=Drosophila melanogaster GN=EG:8D8.3 PE=1 SV=1 | 0.477159211186641 | 0.535057594894842 |
| Q9VT29 | CG16717 OS=Drosophila melanogaster GN=CG16717 PE=1 SV=1 | 0.478195257939166 | 0.519038609600059 |
| O62619 | Pyruvate kinase OS=Drosophila melanogaster GN=PyK PE=2 SV=2 | 0.479230561206336 | 0.475084882948783 |
| P15007 | Enolase OS=Drosophila melanogaster GN=Eno PE=1 SV=2 | 0.480265122054463 | 0.528071164578735 |

| | | | |
|---|---|---|---|
| Q9W058 | Succinyl-CoA:3-ketoacid-coenzyme A transferase OS=Drosophila melanogaster GN=SCOT PE=1 SV=1 | 0.480265122054463 | 0.495695162624069 |
| Q9VED8 | Deoxyribonuclease II OS=Drosophila melanogaster GN=DNaseII PE=1 SV=1 | 0.481298941547565 | 0.43722773891291 |
| Q9VLJ6 | Angiotensin-converting enzyme-related protein OS=Drosophila melanogaster GN=Acer PE=1 SV=1 | 0.481298941547565 | 0.43722773891291 |
| Q9W3M8 | CG1515-PA OS=Drosophila melanogaster GN=Ykt6 PE=1 SV=1 | 0.485426827170242 | 0.454175893185802 |
| A8DYL2 | CG34370, isoform G OS=Drosophila melanogaster GN=CG34370 PE=4 SV=1 | 0.486456955768731 | 0.478195257939166 |
| Q9Y112 | BcDNA.GH10614 OS=Drosophila melanogaster GN=BcDNA.GH10614 PE=1 SV=1 | 0.487486349348536 | 0.541019153133559 |
| Q9VSW2 | RE14081p OS=Drosophila melanogaster GN=UGP PE=1 SV=2 | 0.488515008957812 | 0.453122446595813 |
| Q9W330 | Hexokinase OS=Drosophila melanogaster GN=Hex-A PE=1 SV=1 | 0.490570130446201 | 0.463622456851893 |
| P29613 | Triosephosphate isomerase OS=Drosophila melanogaster GN=Tpi PE=1 SV=3 | 0.492622328574446 | 0.551885103471725 |
| Q8IM93 | CG32017, isoform B OS=Drosophila melanogaster GN=CG32017-RB PE=1 SV=2 | 0.492622328574446 | 0.466757615726171 |
| Q9VXB0 | NECAP-like protein CG9132 OS=Drosophila melanogaster GN=CG9132 PE=2 SV=1 | 0.492622328574446 | 0.492622328574446 |
| Q9VQR0 | CG3246 OS=Drosophila melanogaster GN=CG3246-RA PE=1 SV=1 | 0.492622328574446 | 0.487486349348536 |
| Q8IQX3 | CG32544, isoform B OS=Drosophila melanogaster GN=CG32544 PE=1 SV=1 | 0.496717987935177 | 0.640620928035698 |
| Q9VLX6 | CG7191, isoform A OS=Drosophila melanogaster GN=CG7191 PE=2 SV=3 | 0.496717987935177 | 0.561692721398309 |
| P13217 | 1-phosphatidylinositol 4,5-bisphosphate phosphodiesterase OS=Drosophila melanogaster GN=norpA PE=1 SV=4 | 0.497740088609093 | 0.40381306161659 |
| Q9VFP6 | Inositol polyphosphate 1-phosphatase OS=Drosophila melanogaster GN=Ipp PE=1 SV=1 | 0.498761465671852 | 0.425459304765355 |
| Q9W5G7 | CG13375, isoform B OS=Drosophila melanogaster GN=EG:BACR37P7.8 PE=4 SV=2 | 0.502839758255932 | 0.636914580355878 |
| Q9VM14 | AT21758p OS=Drosophila melanogaster GN=CG5261 PE=1 SV=1 | 0.504874589398464 | 0.405992359675837 |
| Q9W3K9 | CG2254-PA OS=Drosophila melanogaster GN=CG2254 PE=1 SV=1 | 0.504874589398464 | 0.602171790753379 |
| B7Z0B0 | Sosondowah, isoform G OS=Drosophila melanogaster GN=sowah PE=1 SV=2 | 0.505890929729957 | 0.756169328139299 |
| A1Z9J3 | Short stop, isoform H OS=Drosophila melanogaster GN=shot PE=1 SV=1 | 0.506906554580693 | 0.520045024039818 |
| P08120 | Collagen alpha-1(IV) chain OS=Drosophila melanogaster GN=Cg25C PE=2 SV=3 | 0.506906554580693 | 0.545968369105292 |
| Q9VEF9 | CG43102, isoform E OS=Drosophila melanogaster GN=CG43102 PE=4 SV=1 | 0.506906554580693 | 0.487486349348536 |
| X2J979 | Synaptotagmin 1, isoform H OS=Drosophila melanogaster GN=Syt1 PE=4 SV=1 | 0.509949146304311 | 0.511973981781801 |
| Q9V3V2 | FK506-binding protein 14 ortholog, isoform A OS=Drosophila melanogaster GN=Fkbp14 PE=1 SV=1 | 0.509949146304311 | 0.416839741912829 |
| A0A0B4K7L3 | CAP, isoform X OS=Drosophila melanogaster GN=CAP PE=1 SV=1 | 0.509949146304311 | 0.50182126542091 |
| A0A0B4LHA6 | CG7694, isoform C OS=Drosophila melanogaster GN=CG7694 PE=4 SV=1 | 0.511973981781801 | 0.573374526445944 |
| P14199 | Protein ref(2)P OS=Drosophila melanogaster GN=ref(2)P PE=1 SV=2 | 0.511973981781801 | 0.621993231666123 |
| P06604 | Tubulin alpha-2 chain OS=Drosophila melanogaster GN=alphaTub85E PE=1 SV=1 | 0.512985334813676 | 0.670840335881562 |
| Q9VGA3 | CG4115 OS=Drosophila melanogaster GN=CG4115 PE=1 SV=2 | 0.512985334813676 | 0.625738061908648 |
| M9PEL1 | Ras opposite, isoform B OS=Drosophila melanogaster GN=Rop PE=4 SV=1 | 0.51500591643373 | 0.566571640626761 |
| Q9VWD0 | GH23568p OS=Drosophila melanogaster GN=parvin PE=1 SV=2 | 0.51500591643373 | 0.478195257939166 |
| A0A0B4KF38 | Bruchpilot, isoform M OS=Drosophila melanogaster GN=brp PE=1 SV=1 | 0.516015147003665 | 0.699551632523089 |
| A0A0B4K679 | Rab3 interacting molecule, isoform V OS=Drosophila melanogaster GN=Rim PE=4 SV=1 | 0.520045024039818 | 0.499782120147312 |
| Q9V773 | Probable cytochrome P450 6a20 OS=Drosophila melanogaster GN=Cyp6a20 PE=2 SV=2 | 0.521050736900963 | 0.818032474658094 |
| Q8IR95 | RH08789p OS=Drosophila melanogaster GN=ssp7 PE=1 SV=1 | 0.521050736900963 | 0.618238655595455 |
| Q7KVX1 | Pyruvate dehydrogenase E1 component subunit alpha OS=Drosophila melanogaster GN=l(1)G0334 PE=1 SV=1 | 0.522055749160964 | 0.562669826102702 |

| | | | |
|---|---|---|---|
| X2JGF5 | Stoned B, isoform G OS=Drosophila melanogaster GN=stnB PE=4 SV=1 | 0.528071164578735 | 0.486456955768731 |
| Q9VLS7 | CG8552, isoform A OS=Drosophila melanogaster GN=PAPLA1 PE=1 SV=1 | 0.528071164578735 | 0.462575888042204 |
| Q9VA09 | Guanylyl cyclase beta-subunit at 100B OS=Drosophila melanogaster GN=Gycbeta100B PE=1 SV=1 | 0.528071164578735 | 0.558757430373762 |
| Q9VBJ3 | CG42261 OS=Drosophila melanogaster GN=CG42261 PE=4 SV=3 | 0.528071164578735 | 0.506906554580693 |
| A0A0B4KFY8 | CG9813, isoform G OS=Drosophila melanogaster GN=CG9813 PE=1 SV=1 | 0.529071299829111 | 0.57918014812715 |
| Q9VYA1 | CG12177, isoform A OS=Drosophila melanogaster GN=CG12177 PE=2 SV=1 | 0.530070742225084 | 0.452068230223811 |
| Q9VXC1 | CG34325, isoform A OS=Drosophila melanogaster GN=CG34325-RA PE=2 SV=3 | 0.531069492725954 | 0.513995979367013 |
| A1Z7R9 | CG42382 OS=Drosophila melanogaster GN=rad201 PE=4 SV=1 | 0.533064921869638 | 0.542010355536609 |
| M9NF14 | IGF-II mRNA-binding protein, isoform L OS=Drosophila melanogaster GN=Imp PE=1 SV=1 | 0.534061602421118 | 0.465713319595964 |
| Q9V3N7 | BcDNA.HL02693 OS=Drosophila melanogaster GN=CRMP PE=1 SV=1 | 0.534061602421118 | 0.499782120147312 |
| Q9W2M4 | CG10527 OS=Drosophila melanogaster GN=CG10527 PE=1 SV=1 | 0.534061602421118 | 0.519038609600059 |
| Q9W1D9 | Oxysterol-binding protein OS=Drosophila melanogaster GN=CG3860 PE=1 SV=1 | 0.534061602421118 | 0.464668267003444 |
| M9ND55 | Multiplexin, isoform R OS=Drosophila melanogaster GN=Mp PE=1 SV=1 | 0.534061602421118 | 0.482332020747376 |
| A0A0B4KFZ3 | Phosphodiesterase 6, isoform C OS=Drosophila melanogaster GN=Pde6 PE=4 SV=1 | 0.535057594894842 | 0.641546029087524 |
| P13706 | Glycerol-3-phosphate dehydrogenase [NAD(+)], cytoplasmic OS=Drosophila melanogaster GN=Gpdh PE=1 SV=3 | 0.537047519404657 | 0.574343753920013 |
| Q9W3W4 | COQ7 OS=Drosophila melanogaster GN=COQ7 PE=1 SV=2 | 0.537047519404657 | 0.43722773891291 |
| A0A0B4KGU5 | Eclosion hormone, isoform B OS=Drosophila melanogaster GN=Eh PE=4 SV=1 | 0.537047519404657 | 0.529071299829111 |
| Q9VHK6 | CG9836 OS=Drosophila melanogaster GN=IscU PE=1 SV=1 | 0.539034702970754 | 0.492622328574446 |
| B7YZI0 | Vacuolar H[+] ATPase 44kD subunit, isoform F OS=Drosophila melanogaster GN=Vha44 PE=1 SV=1 | 0.540027269257507 | 0.526068811667588 |
| Q7KK90 | GH14654p OS=Drosophila melanogaster GN=GstE1 PE=1 SV=1 | 0.540027269257507 | 0.405992359675837 |
| Q8MLP9 | CG30172, isoform A OS=Drosophila melanogaster GN=CSP1 PE=2 SV=1 | 0.541019153133559 | 0.526068811667588 |
| Q9VGF3 | CG18547 OS=Drosophila melanogaster GN=CG18547 PE=1 SV=1 | 0.542010355536609 | 0.489542935642474 |
| Q9VCM6 | CG4393, isoform B OS=Drosophila melanogaster GN=CG4393 PE=2 SV=4 | 0.542010355536609 | 0.53605290024021 |
| Q7JR83 | Hormone-sensitive lipase ortholog, isoform A OS=Drosophila melanogaster GN=Hsl PE=1 SV=1 | 0.542010355536609 | 0.524063675777211 |
| Q9VCU1 | CG4721 OS=Drosophila melanogaster GN=CG4721-RA PE=2 SV=2 | 0.543000877402426 | 0.584962500721156 |
| Q9V3T9 | NADPH:adrenodoxin oxidoreductase, mitochondrial OS=Drosophila melanogaster GN=dare PE=2 SV=1 | 0.544979883255804 | 0.490570130446201 |
| X2JES6 | CG1552, isoform C OS=Drosophila melanogaster GN=CG1552 PE=1 SV=1 | 0.544979883255804 | 0.82130203988953 |
| Q9VL70 | CG4600-PA OS=Drosophila melanogaster GN=yip2 PE=1 SV=1 | 0.545968369105292 | 0.553851968181126 |
| Q9VDH3 | CG10830 OS=Drosophila melanogaster GN=Ktl PE=1 SV=1 | 0.54794331129035 | 0.58207422139633 |
| Q9W3J1 | Gbeta5 OS=Drosophila melanogaster GN=Gbeta5 PE=1 SV=1 | 0.5489297694764 | 0.506906554580693 |
| P52034 | ATP-dependent 6-phosphofructokinase OS=Drosophila melanogaster GN=Pfk PE=2 SV=2 | 0.552868871011303 | 0.497740088609093 |
| Q9VIX7 | CG15825-PB, isoform B OS=Drosophila melanogaster GN=fon PE=1 SV=1 | 0.552868871011303 | 0.540027269257507 |
| Q7K0S5 | GDI interacting protein 3, isoform B OS=Drosophila melanogaster GN=Gint3 PE=1 SV=1 | 0.552868871011303 | 0.54399071966485 |
| M9MRD1 | Muscle-specific protein 300 kDa, isoform D OS=Drosophila melanogaster GN=Msp300 PE=1 SV=1 | 0.554834395894193 | 0.58014548442338 |
| Q9VII9 | CG31673, isoform A OS=Drosophila melanogaster GN=CG31673 PE=1 SV=2 | 0.554834395894193 | 0.471967787661516 |
| Q95SI7 | CG6028 OS=Drosophila melanogaster GN=CG6028 PE=1 SV=1 | 0.557777671394926 | 0.525066592078211 |
| X2JDA5 | Glutamine synthetase OS=Drosophila melanogaster GN=Gs2 PE=1 SV=1 | 0.558757430373762 | 0.54794331129035 |

| | | | |
|---|---|---|---|
| Q9U6R9 | GH13039p OS=Drosophila melanogaster GN=gammaSnap1 PE=1 SV=1 | 0.560714954474479 | 0.578214165472454 |
| P46461 | Vesicle-fusing ATPase 1 OS=Drosophila melanogaster GN=comt PE=2 SV=1 | 0.561692721398309 | 0.552868871011303 |
| A0A0B4LHE7 | Vacuolar H[+] ATPase 13kD subunit, isoform B OS=Drosophila melanogaster GN=Vha13 PE=4 SV=1 | 0.562669826102702 | 0.633198686374004 |
| Q9U9P7 | Cytoplasmic phosphatidylinositol transfer protein 1 OS=Drosophila melanogaster GN=rdgBbeta PE=2 SV=1 | 0.562669826102702 | 0.504874589398464 |
| Q9VG31 | Malic enzyme OS=Drosophila melanogaster GN=Men PE=1 SV=1 | 0.565597175854225 | 0.622930350920177 |
| Q8SXQ5 | CG14407 OS=Drosophila melanogaster GN=CG14407 PE=1 SV=1 | 0.566571640626761 | 0.519038609600059 |
| B7YZV2 | Pde1c, isoform E OS=Drosophila melanogaster GN=Pde1c PE=1 SV=1 | 0.567545447643747 | 0.525066592078211 |
| Q9VPV8 | IA-2 ortholog, isoform C OS=Drosophila melanogaster GN=IA-2 PE=1 SV=4 | 0.569491091958716 | 0.636914580355878 |
| Q9VG55 | Protein hugin OS=Drosophila melanogaster GN=Hug PE=1 SV=1 | 0.571434115876509 | 0.576280257621355 |
| Q9VTV9 | Delta-aminolevulinic acid dehydratase OS=Drosophila melanogaster GN=Pbgs PE=1 SV=1 | 0.573374526445944 | 0.657182660128423 |
| Q8IPM8 | Complexin OS=Drosophila melanogaster GN=cpx PE=2 SV=1 | 0.574343753920013 | 0.562669826102702 |
| P00334 | Alcohol dehydrogenase OS=Drosophila melanogaster GN=Adh PE=1 SV=2 | 0.575312330687437 | 0.618238655595455 |
| Q8IQ31 | CG1695 OS=Drosophila melanogaster GN=CG1695 PE=4 SV=2 | 0.578214165472454 | 0.623866861852698 |
| A0A0B4KGY6 | Pasilla, isoform R OS=Drosophila melanogaster GN=ps PE=1 SV=1 | 0.583037623796664 | 0.593592805864596 |
| Q01583 | Diacylglycerol kinase 1 OS=Drosophila melanogaster GN=Dgk PE=2 SV=5 | 0.583037623796664 | 0.848798181244189 |
| E2QCF1 | ATP-citrate synthase OS=Drosophila melanogaster GN=ATPCL PE=1 SV=1 | 0.584962500721156 | 0.491596594410448 |
| Q8INQ9 | AT21926p OS=Drosophila melanogaster GN=stck PE=1 SV=1 | 0.585923976958601 | 0.573374526445944 |
| A1Z9M6 | Vesicular GABA transporter OS=Drosophila melanogaster GN=VGAT PE=1 SV=1 | 0.585923976958601 | 0.634128557525041 |
| Q9VFC8 | Glycogen [starch] synthase OS=Drosophila melanogaster GN=GlyS PE=1 SV=2 | 0.587845009254277 | 0.540027269257507 |
| Q9VAN7 | GH13304p OS=Drosophila melanogaster GN=Pglym78 PE=1 SV=2 | 0.58880456701555 | 0.666302128173095 |
| Q9VZG1 | Cuticular protein 64Ab OS=Drosophila melanogaster GN=Cpr64Ab PE=4 SV=2 | 0.58880456701555 | 0.687956494044482 |
| Q7K4Q9 | HMG coenzyme A synthase, isoform A OS=Drosophila melanogaster GN=Hmgs PE=1 SV=1 | 0.590721770009841 | 0.559736524432983 |
| Q7K5K3 | CG11876, isoform A OS=Drosophila melanogaster GN=CG11876 PE=1 SV=1 | 0.59741198755465 | 0.587845009254277 |
| P20477 | Glutamine synthetase 1, mitochondrial OS=Drosophila melanogaster GN=Gs1 PE=2 SV=3 | 0.598365205323645 | 0.668119124686432 |
| Q7JVH0 | CG8435 OS=Drosophila melanogaster GN=CG8435 PE=1 SV=1 | 0.598365205323645 | 0.523060061795249 |
| O97477 | Inositol-3-phosphate synthase OS=Drosophila melanogaster GN=Inos PE=1 SV=1 | 0.601221085584946 | 0.492622328574446 |
| Q9VGE7 | Beta-galactosidase OS=Drosophila melanogaster GN=Ect3 PE=1 SV=1 | 0.601221085584946 | 0.591679416935737 |
| E1JHT6 | Reticulon-like protein OS=Drosophila melanogaster GN=Rtnl1 PE=1 SV=1 | 0.603121869839996 | 0.611644543373662 |
| Q23983 | Alpha-soluble NSF attachment protein OS=Drosophila melanogaster GN=alphaSnap PE=1 SV=1 | 0.605968358841458 | 0.645240512645265 |
| M9PBJ1 | Zormin, isoform J OS=Drosophila melanogaster GN=zormin PE=1 SV=1 | 0.606915941825205 | 0.654435540845399 |
| P15215 | Laminin subunit gamma-1 OS=Drosophila melanogaster GN=LanB2 PE=2 SV=2 | 0.607862902831235 | 0.660837367696284 |
| A0A0B4K620 | Mustard, isoform V OS=Drosophila melanogaster GN=mtd PE=1 SV=1 | 0.610700062134764 | 0.564622052436981 |
| M9NE45 | Ectoderm-expressed 4, isoform I OS=Drosophila melanogaster GN=Ect4 PE=1 SV=1 | 0.611644543373662 | 0.519038609600059 |
| Q9W401 | Probable citrate synthase, mitochondrial OS=Drosophila melanogaster GN=kdn PE=2 SV=1 | 0.613531652917927 | 0.577247535593151 |
| X2J6D4 | Cytochrome c proximal, isoform B OS=Drosophila melanogaster GN=Cyt-c-p PE=3 SV=1 | 0.614474282837701 | 0.593592805864596 |
| B5RIU6 | AT06279p OS=Drosophila melanogaster GN=EndoA PE=1 SV=1 | 0.617298482840846 | 0.682573297347578 |
| Q8MMD2 | Epidermal growth factor receptor pathway substrate clone 15, isoform B OS=Drosophila melanogaster GN=Eps-15 PE=1 SV=1 | 0.619178216059069 | 0.58880456701555 |

| | | | |
|---|---|---|---|
| Q9W0Y1 | Troponin C-akin-1 protein OS=Drosophila melanogaster GN=Tina-1 PE=2 SV=1 | 0.619178216059069 | 0.558757430373762 |
| Q9VVV3 | CG14075 OS=Drosophila melanogaster GN=CG14075 PE=4 SV=1 | 0.620117165028684 | 0.742437445376266 |
| Q9VIQ0 | Short neuropeptide F OS=Drosophila melanogaster GN=sNPF PE=1 SV=4 | 0.621993231666123 | 0.593592805864596 |
| Q03427 | Lamin-C OS=Drosophila melanogaster GN=LamC PE=1 SV=2 | 0.623866861852698 | 0.68885174386588 |
| E1JHJ3 | Myosin heavy chain, isoform O OS=Drosophila melanogaster GN=Mhc PE=1 SV=1 | 0.624802765252947 | 0.520045024039818 |
| Q9VIM0 | CG2493, isoform A OS=Drosophila melanogaster GN=CG2493 PE=1 SV=1 | 0.626672752605993 | 0.5489297694764 |
| Q9VDL4 | CG10877 OS=Drosophila melanogaster GN=CG10877 PE=1 SV=1 | 0.62760683812965 | 0.632268215499513 |
| Q9VJ31 | CG10623 protein OS=Drosophila melanogaster GN=CG10623 PE=1 SV=1 | 0.62760683812965 | 0.617298482840846 |
| Q8IQU7 | 825-Oak OS=Drosophila melanogaster GN=825-Oak PE=4 SV=2 | 0.631337144127481 | 0.743299527888257 |
| Q9VVZ7 | CG18294 OS=Drosophila melanogaster GN=CG18294 PE=1 SV=2 | 0.632268215499513 | 0.637842060324105 |
| Q9Y136 | CG14526 OS=Drosophila melanogaster GN=CG14526 PE=1 SV=2 | 0.634128557525041 | 0.753604536279995 |
| Q9VF15 | Globin 1, isoform A OS=Drosophila melanogaster GN=glob1 PE=1 SV=1 | 0.639695233399582 | 0.57918014812715 |
| Q9V8Y2 | General odorant-binding protein 56a OS=Drosophila melanogaster GN=Obp56a PE=1 SV=1 | 0.639695233399582 | 0.65901117119815 |
| X2J9Z1 | Silver, isoform N OS=Drosophila melanogaster GN=svr PE=1 SV=1 | 0.644317778337577 | 0.646162657157894 |
| M9PD18 | Vacuolar H[+] ATPase 68kD subunit 1, isoform B OS=Drosophila melanogaster GN=Vha68-1 PE=1 SV=1 | 0.645240512645265 | 0.608809242675524 |
| A1Z877 | LP19846p OS=Drosophila melanogaster GN=Ndg PE=1 SV=1 | 0.645240512645265 | 0.63876894441695 |
| Q9VGK3 | Peptidyl-prolyl cis-trans isomerase OS=Drosophila melanogaster GN=CG14715 PE=1 SV=1 | 0.645240512645265 | 0.695102985640479 |
| Q8SWS3 | CG13049 OS=Drosophila melanogaster GN=CG13049 PE=1 SV=1 | 0.648925559453121 | 0.565597175854225 |
| Q9TVP3 | J domain-containing protein OS=Drosophila melanogaster GN=jdp PE=2 SV=2 | 0.64984535230601 | 0.692427198089708 |
| A8JNS4 | Starvin, isoform E OS=Drosophila melanogaster GN=stv PE=1 SV=1 | 0.652601217596519 | 0.657182660128423 |
| Q9VS89 | FI18763p1 OS=Drosophila melanogaster GN=frac PE=2 SV=4 | 0.654435540845399 | 0.686164326061359 |
| Q7K3E2 | CG5080, isoform A OS=Drosophila melanogaster GN=CG5080 PE=1 SV=1 | 0.654435540845399 | 0.651683180632109 |
| C8VV14 | Fructose-bisphosphate aldolase OS=Drosophila melanogaster GN=Ald PE=1 SV=1 | 0.655351828612554 | 0.680774425492461 |
| Q9VWV6 | Transferrin 1, isoform A OS=Drosophila melanogaster GN=Tsf1 PE=1 SV=1 | 0.656267534794289 | 0.764685852522134 |
| O97062 | Ccp84Ae OS=Drosophila melanogaster GN=Ccp84Ae PE=1 SV=1 | 0.656267534794289 | 0.641546029087524 |
| Q26377 | Pro-corazonin OS=Drosophila melanogaster GN=Crz PE=1 SV=2 | 0.658097205351372 | 0.693319678811575 |
| Q9VU19 | Wbp2 ortholog, isoform E OS=Drosophila melanogaster GN=Wbp2 PE=1 SV=3 | 0.65901117119815 | 0.765534746362977 |
| Q7JRL9 | CG31221, isoform A OS=Drosophila melanogaster GN=CG31221 PE=1 SV=1 | 0.660837367696284 | 0.644317778337577 |
| Q9VTP0 | CG42255 OS=Drosophila melanogaster GN=CG42255 PE=1 SV=4 | 0.663572335417523 | 0.647084212628954 |
| P11046 | Laminin subunit beta-1 OS=Drosophila melanogaster GN=LanB1 PE=1 SV=4 | 0.669026765509631 | 0.732052073300289 |
| Q9VQC0 | Vesicular glutamate transporter, isoform A OS=Drosophila melanogaster GN=VGlut PE=1 SV=2 | 0.669026765509631 | 0.631337144127481 |
| Q8IQS5 | CG32195, isoform A OS=Drosophila melanogaster GN=CG32195 PE=4 SV=1 | 0.672651629328385 | 0.633198686374004 |
| Q00174 | Laminin subunit alpha OS=Drosophila melanogaster GN=LanA PE=1 SV=2 | 0.674460651560259 | 0.662661255475094 |
| Q9V496 | Apolipophorins OS=Drosophila melanogaster GN=Rfabg PE=1 SV=2 | 0.674460651560259 | 0.683471892855228 |
| Q9VA42 | Niemann-Pick type C-2g, isoform A OS=Drosophila melanogaster GN=Npc2g PE=1 SV=1 | 0.67626740826589 | 0.727702672837238 |
| Q9VBA0 | CG6330, isoform A OS=Drosophila melanogaster GN=CG6330 PE=1 SV=2 | 0.67897330785417 | 0.658097205351372 |
| Q9W4W5 | CG2680 OS=Drosophila melanogaster GN=EG:100G10.4 PE=1 SV=2 | 0.679874147746623 | 0.741574847418796 |

| | | | |
|---|---|---|---|
| M9NGG5 | Futsch, isoform F OS=Drosophila melanogaster GN=futsch PE=1 SV=1 | 0.683471892855228 | 0.684369929013007 |
| Q9VFI3 | CG8066, isoform A OS=Drosophila melanogaster GN=CG8066 PE=1 SV=1 | 0.683471892855228 | 0.860764202628828 |
| Q9VVE2 | Protein rogdi OS=Drosophila melanogaster GN=rogdi PE=1 SV=2 | 0.685267406516842 | 0.698662999885884 |
| Q7KTA1 | CG31839-PA OS=Drosophila melanogaster GN=NimB2 PE=2 SV=1 | 0.687060688339892 | 0.756169328139299 |
| E2QCY9 | Synapsin, isoform D OS=Drosophila melanogaster GN=Syn PE=1 SV=1 | 0.687956494044482 | 0.675364312749146 |
| P05031 | Aromatic-L-amino-acid decarboxylase OS=Drosophila melanogaster GN=Ddc PE=1 SV=4 | 0.68885174386588 | 0.708407983483596 |
| P10552 | FMRFamide-related peptides OS=Drosophila melanogaster GN=FMRFa PE=1 SV=2 | 0.693319678811575 | 0.789937868980195 |
| Q95028 | L-lactate dehydrogenase OS=Drosophila melanogaster GN=ImpL3 PE=2 SV=1 | 0.6959938131099 | 0.6535186707536 |
| Q7KTI5 | CG8086, isoform L OS=Drosophila melanogaster GN=CG8086 PE=1 SV=3 | 0.696884090855454 | 0.801572569463598 |
| L0MLR4 | Calcium/calmodulin-dependent protein kinase II, isoform L OS=Drosophila melanogaster GN=CaMKII PE=1 SV=1 | 0.696884090855454 | 0.603121869839996 |
| A4V4W0 | RH38069p1 OS=Drosophila melanogaster GN=stnA PE=1 SV=1 | 0.697773819555186 | 0.708407983483596 |
| P82890 | Low molecular weight phosphotyrosine protein phosphatase 1 OS=Drosophila melanogaster GN=primo-1 PE=2 SV=1 | 0.697773819555186 | 0.722466024471091 |
| Q9VLU6 | Succinate dehydrogenase assembly factor 4, mitochondrial OS=Drosophila melanogaster GN=Sirup PE=3 SV=2 | 0.703986603863431 | 0.73725410432433 |
| Q9VNH7 | CG2082, isoform A OS=Drosophila melanogaster GN=CG2082 PE=1 SV=1 | 0.704871964456353 | 0.727702672837238 |
| Q95RR6 | LD15002p OS=Drosophila melanogaster GN=Pur-alpha PE=1 SV=1 | 0.704871964456353 | 0.715454127115718 |
| A0A0B4KFM8 | Metallothionein A, isoform B OS=Drosophila melanogaster GN=MtnA PE=4 SV=1 | 0.704871964456353 | 0.739848102699327 |
| A1ZBK7 | Crammer OS=Drosophila melanogaster GN=cer PE=1 SV=1 | 0.711935356978922 | 0.781569544815974 |
| Q9NIP6 | Cardio acceleratory peptide 2b OS=Drosophila melanogaster GN=Capa PE=1 SV=1 | 0.712815854437372 | 0.743299527888257 |
| Q9VSU8 | Nervous wreck, isoform D OS=Drosophila melanogaster GN=nwk PE=1 SV=3 | 0.715454127115718 | 0.655351828612554 |
| Q59E09 | Acetyl-coenzyme A synthetase OS=Drosophila melanogaster GN=AcCoAS PE=1 SV=2 | 0.715454127115718 | 0.590721770009841 |
| Q9VLV9 | Proctolin OS=Drosophila melanogaster GN=Proc PE=2 SV=2 | 0.715454127115718 | 0.738983954700512 |
| Q8SYD9 | Endophilin B, isoform A OS=Drosophila melanogaster GN=EndoB PE=1 SV=1 | 0.718087583960517 | 0.835924074254375 |
| X2JCV2 | Larval serum protein 2, isoform B OS=Drosophila melanogaster GN=Lsp2 PE=4 SV=1 | 0.72159139877538 | 0.661749599810705 |
| P36951 | Putative hydroxypyruvate isomerase OS=Drosophila melanogaster GN=Gip PE=2 SV=1 | 0.723340120251171 | 0.663572335417523 |
| Q7JWW6 | Ady43A OS=Drosophila melanogaster GN=Ady43A PE=2 SV=1 | 0.725959234509188 | 0.675364312749146 |
| Q23997 | Chitinase-like protein CG5210 OS=Drosophila melanogaster GN=CG5210 PE=1 SV=2 | 0.726831217032493 | 0.741574847418796 |
| Q9VPY9 | CG5001, isoform B OS=Drosophila melanogaster GN=CG5001-RA PE=2 SV=3 | 0.728573602559365 | 0.417920007811965 |
| Q9VSY0 | Cuticular protein 67B OS=Drosophila melanogaster GN=Cpr67B PE=1 SV=1 | 0.735522177296537 | 0.942608336111663 |
| A1Z7Z4 | CG1648, isoform B OS=Drosophila melanogaster GN=CG1648 PE=1 SV=1 | 0.738983954700512 | 0.747602230274943 |
| E1JJ78 | Like-AP180, isoform D OS=Drosophila melanogaster GN=lap PE=1 SV=1 | 0.742437445376266 | 0.680774425492461 |
| D1YSG8 | Calcium/calmodulin-dependent protein kinase II, isoform J OS=Drosophila melanogaster GN=CaMKII PE=1 SV=1 | 0.743299527888257 | 0.774839759827326 |
| Q9NGX9 | Cytochrome P450 302a1, mitochondrial OS=Drosophila melanogaster GN=dib PE=2 SV=2 | 0.745882688902259 | 0.825378603892931 |
| Q9W4Y3 | Terribly reduced optic lobes, isoform AT OS=Drosophila melanogaster GN=trol PE=1 SV=4 | 0.750177705593057 | 0.834306702962908 |
| M9MRJ4 | Muscle-specific protein 300 kDa, isoform G OS=Drosophila melanogaster GN=Msp300 PE=1 SV=1 | 0.751035176667862 | 0.759581973233555 |
| Q9W4Y1 | CG13759, isoform A OS=Drosophila melanogaster GN=EG:BACR25B3.5 PE=2 SV=2 | 0.752748591407134 | 0.848798181244189 |
| Q9VTJ4 | Putative alpha-L-fucosidase OS=Drosophila melanogaster GN=Fuca PE=2 SV=2 | 0.754459973625479 | 0.795766947782392 |
| Q9XTL9 | Glycogen phosphorylase OS=Drosophila melanogaster GN=GlyP PE=2 SV=2 | 0.756169328139299 | 0.793271654498083 |

| | | | |
|---|---|---|---|
| Q9VZF9 | Cuticular protein 64Ad OS=Drosophila melanogaster GN=Cpr64Ad PE=1 SV=2 | 0.759581973233555 | 0.833497336859835 |
| Q9VU77 | CG10133 OS=Drosophila melanogaster GN=CG10133 PE=1 SV=1 | 0.760433874670112 | 0.747602230274943 |
| P82147 | Protein lethal(2)essential for life OS=Drosophila melanogaster GN=l(2)efl PE=1 SV=1 | 0.762136169901112 | 0.762136169901112 |
| P81900 | cAMP-dependent protein kinase type II regulatory subunit OS=Drosophila melanogaster GN=Pka-R2 PE=1 SV=2 | 0.762986564880576 | 0.759581973233555 |
| A1ZB68 | FI01423p OS=Drosophila melanogaster GN=GstE3 PE=1 SV=1 | 0.768078435016455 | 0.722466024471091 |
| Q8I941 | CG32485 OS=Drosophila melanogaster GN=CG32485 PE=2 SV=1 | 0.768078435016455 | 0.814755482809874 |
| A8DYP0 | Unc-89, isoform C OS=Drosophila melanogaster GN=Unc-89 PE=1 SV=1 | 0.768925335563751 | 0.814755482809874 |
| Q9VCH7 | CG10208 OS=Drosophila melanogaster GN=CG10208 PE=1 SV=1 | 0.772307974958403 | 0.865522959139964 |
| X2JGQ6 | CG45057, isoform E OS=Drosophila melanogaster GN=CG45057 PE=4 SV=1 | 0.773152397014052 | 0.729444006833663 |
| A1Z6N4 | MIP05841p OS=Drosophila melanogaster GN=Tdc2 PE=2 SV=1 | 0.773152397014052 | 0.577247535593151 |
| P20228 | Glutamate decarboxylase OS=Drosophila melanogaster GN=Gad1 PE=2 SV=2 | 0.780730036476377 | 0.747602230274943 |
| A1Z9F4 | CG33156, isoform E OS=Drosophila melanogaster GN=CG33156 PE=3 SV=1 | 0.794935662803536 | 0.776525151421912 |
| M9PDR0 | Thioester-containing protein 4, isoform C OS=Drosophila melanogaster GN=Tep4 PE=1 SV=1 | 0.7982579326445 | 0.860764202628828 |
| Q9VA76 | CG15537, isoform B OS=Drosophila melanogaster GN=CG15537-RB PE=2 SV=2 | 0.7982579326445 | 0.704871964456353 |
| A0A0B4K7K9 | Bruchpilot, isoform J OS=Drosophila melanogaster GN=brp PE=1 SV=1 | 0.800744623936778 | 0.838346736521231 |
| O96299 | LD47736p OS=Drosophila melanogaster GN=Sodh-2 PE=1 SV=1 | 0.803227036434928 | 0.679874147746623 |
| X2J8F5 | Larval serum protein 1 beta, isoform B OS=Drosophila melanogaster GN=Lsp1beta PE=4 SV=1 | 0.805705184838511 | 0.752748591407134 |
| Q9VNX4 | Delta-1-Pyrroline-5-carboxylate dehydrogenase 1, isoform A OS=Drosophila melanogaster GN=P5CDh1 PE=1 SV=1 | 0.807354922057604 | 0.784922701562861 |
| Q9VLC5 | Aldehyde dehydrogenase OS=Drosophila melanogaster GN=Aldh PE=1 SV=1 | 0.809002774939086 | 0.723340120251171 |
| Q9VIF7 | CG1099-PA, isoform A OS=Drosophila melanogaster GN=Dap160 PE=1 SV=1 | 0.811471030529836 | 0.813114191346289 |
| Q9VXF1 | Serine/threonine-protein phosphatase 2B catalytic subunit 3 OS=Drosophila melanogaster GN=CanA-14F PE=1 SV=4 | 0.811471030529836 | 0.828631581688019 |
| Q9W2M2 | Trehalase OS=Drosophila melanogaster GN=Treh PE=1 SV=1 | 0.819668183496456 | 0.851998837112446 |
| A1ZB72 | Glutathione S transferase E7 OS=Drosophila melanogaster GN=GstE7 PE=1 SV=1 | 0.82130203988953 | 0.894526770257395 |
| Q9W4J9 | AT21585p OS=Drosophila melanogaster GN=CG3568 PE=1 SV=2 | 0.822118274729345 | 0.657182660128423 |
| Q8IPU1 | AT19489p OS=Drosophila melanogaster GN=CG33054-RB PE=1 SV=1 | 0.82781902461732 | 0.644317778337577 |
| Q7JYX0 | Glutathione S-transferase E14 OS=Drosophila melanogaster GN=GstE14 PE=1 SV=1 | 0.829443681366591 | 0.776525151421912 |
| Q9VHK7 | CG8369, isoform A OS=Drosophila melanogaster GN=CG8369 PE=1 SV=1 | 0.830255324167683 | 0.84639302085236 |
| Q03042 | cGMP-dependent protein kinase, isozyme 1 OS=Drosophila melanogaster GN=Pkg21D PE=1 SV=2 | 0.831066510605073 | 1,041242982231880 |
| Q9V3D4 | Chitinase-like protein Idgf2 OS=Drosophila melanogaster GN=Idgf2 PE=1 SV=1 | 0.832687516439532 | 0.767231037023876 |
| Q9W332 | Cubilin ortholog OS=Drosophila melanogaster GN=Cubn PE=1 SV=3 | 0.833497336859835 | 0.851199338593294 |
| E1JJA4 | Shibire, isoform L OS=Drosophila melanogaster GN=shi PE=1 SV=1 | 0.836732080459136 | 0.583037623796664 |
| A1Z847 | CG2269, isoform B OS=Drosophila melanogaster GN=CG2269 PE=1 SV=1 | 0.843179890257174 | 0.860764202628828 |
| Q7KUD5 | Probable insulin-like peptide 5 OS=Drosophila melanogaster GN=Ilp5 PE=1 SV=2 | 0.848798181244189 | 0.715454127115718 |
| Q9VWR5 | Cytochrome P450 306a1 OS=Drosophila melanogaster GN=phm PE=1 SV=1 | 0.852797892818771 | 0.831066510605073 |
| Q961D1 | CG15218-PA, isoform A OS=Drosophila melanogaster GN=CycK PE=2 SV=1 | 0.852797892818771 | 1,08065766334523 |
| B7Z0B8 | CG10625, isoform I OS=Drosophila melanogaster GN=CG10625 PE=1 SV=1 | 0.854394677753101 | 0.870266070395983 |
| Q8SZA8 | CG1319 OS=Drosophila melanogaster GN=Fdx2 PE=1 SV=1 | 0.859174455866435 | 0.972325041557152 |

| | | | |
|---|---|---|---|
| Q24400 | Muscle LIM protein Mlp84B OS=Drosophila melanogaster GN=Mlp84B PE=1 SV=1 | 0.859969548221026 | 0.806530289259566 |
| A0A0B4KGT7 | Myosuppressin, isoform B OS=Drosophila melanogaster GN=Ms PE=4 SV=1 | 0.861558419572029 | 0.859174455866435 |
| Q9VEB1 | IP09655p OS=Drosophila melanogaster GN=Mdh2 PE=1 SV=1 | 0.865522959139964 | 0.828631581688019 |
| X2JDD7 | Tiggrin, isoform B OS=Drosophila melanogaster GN=Tig PE=1 SV=1 | 0.867105729502655 | 0.929033478645856 |
| Q9VG92 | Glutathione S transferase D8 OS=Drosophila melanogaster GN=GstD8 PE=2 SV=1 | 0.867896463992655 | 1 |
| E2QD99 | CG34417, isoform I OS=Drosophila melanogaster GN=CG34417 PE=1 SV=1 | 0.874206785877799 | 0.907659827853289 |
| M9PEA0 | Sallimus, isoform P OS=Drosophila melanogaster GN=sls PE=4 SV=1 | 0.874993638932967 | 0.84639302085236 |
| A0A0B4K843 | Bruchpilot, isoform I OS=Drosophila melanogaster GN=brp PE=1 SV=1 | 0.876566058751721 | 0.973794929652606 |
| Q9VGH1 | Cytochrome P450 315a1, mitochondrial OS=Drosophila melanogaster GN=sad PE=2 SV=1 | 0.882838655767251 | 0.869476633965402 |
| D1FYT3 | Odorant-binding protein 99b OS=Drosophila melanogaster GN=Obp99b PE=1 SV=1 | 0.885183866320351 | 0.824564212089827 |
| Q9W1F8 | CG13565, isoform A OS=Drosophila melanogaster GN=Orcokinin PE=2 SV=2 | 0.885183866320351 | 0.908428650168787 |
| Q09024 | Neural/ectodermal development factor IMP-L2 OS=Drosophila melanogaster GN=ImpL2 PE=1 SV=4 | 0.892196710465485 | 0.978195629681652 |
| Q8MM24 | Chitinase-like protein Idgf1 OS=Drosophila melanogaster GN=Idgf1 PE=1 SV=2 | 0.896853072896707 | 0.967906370290147 |
| Q9VFC7 | Mf5 protein OS=Drosophila melanogaster GN=Mf PE=1 SV=2 | 0.898401859992193 | 0.981121989794311 |
| Q86B94 | FI19380p1 OS=Drosophila melanogaster GN=mfas PE=1 SV=1 | 0.906890595608518 | 0.969380764941608 |
| P42281 | Acyl-CoA-binding protein homolog OS=Drosophila melanogaster GN=Dbi PE=2 SV=1 | 0.908428650168787 | 0.900721927564115 |
| A1ZB71 | Glutathione S transferase E6 OS=Drosophila melanogaster GN=GstE6 PE=1 SV=1 | 0.911499848861108 | 0.865522959139964 |
| Q9VIB5 | Carboxylic ester hydrolase OS=Drosophila melanogaster GN=alpha-Est7 PE=1 SV=1 | 0.916858764699754 | 0.899948986189672 |
| Q9VE01 | Probable cytochrome P450 12a5, mitochondrial OS=Drosophila melanogaster GN=Cyp12a5 PE=2 SV=1 | 0.923719678749389 | 0.903809558578697 |
| Q7K0P0 | Juvenile hormone-inducible protein 26 OS=Drosophila melanogaster GN=JhI-26 PE=1 SV=1 | 0.94335876267781 | 1,2147465226824 |
| Q9VY05 | CG9512, isoform A OS=Drosophila melanogaster GN=CG9512 PE=1 SV=1 | 0.971589535530062 | 1,1427401721161 |
| Q9VH98 | Diuretic hormone 44, isoform A OS=Drosophila melanogaster GN=Dh44 PE=2 SV=4 | 0.973060172804084 | 1,0335111023613 |
| B5X0J4 | FI04429p OS=Drosophila melanogaster GN=ple PE=1 SV=1 | 0.974529312483882 | 0.978927776381399 |
| Q9V3Y7 | CG15293, isoform A OS=Drosophila melanogaster GN=CG15293 PE=1 SV=1 | 0.975263321678493 | 1,03139519627553 |
| A1ZA47 | PDZ and LIM domain protein Zasp OS=Drosophila melanogaster GN=Zasp52 PE=1 SV=2 | 0.996388746447621 | 0.940354711533293 |
| P06754 | Tropomyosin-1, isoforms 9A/A/B OS=Drosophila melanogaster GN=Tm1 PE=2 SV=2 | 1,00216242115062 | 0.999278472082541 |
| Q9VBS7 | CG10550, isoform B OS=Drosophila melanogaster GN=CG10550 PE=1 SV=1 | 1,00719550140420 | 0.855192407959154 |
| Q07171 | Gelsolin OS=Drosophila melanogaster GN=Gel PE=1 SV=2 | 1,01077983875324 | 1,03139519627553 |
| Q7JZW0 | Cuticular protein 51A OS=Drosophila melanogaster GN=Cpr51A PE=1 SV=1 | 1,02076886509457 | 0,937344392 |
| Q9XZ56 | 4E-binding protein THOR OS=Drosophila melanogaster GN=Thor PE=1 SV=1 | 1,03491998434916 | 1,46727948045998 |
| Q9W145 | Putative cholesterol transporter OS=Drosophila melanogaster GN=Start1 PE=2 SV=2 | 1,04404433270602 | 1,14077865578280 |
| Q9VFZ4 | CG7966 OS=Drosophila melanogaster GN=CG7966 PE=1 SV=2 | 1,04893364519792 | 1,07176266930009 |
| Q9I7J0 | CG5023 OS=Drosophila melanogaster GN=CG5023 PE=1 SV=1 | 1,04893364519792 | 1,02644598030380 |
| Q9W303 | Chitinase-like protein Idgf4 OS=Drosophila melanogaster GN=Idgf4 PE=2 SV=1 | 1,05241589415114 | 1,07860983469637 |
| P11995 | Larval serum protein 1 alpha chain OS=Drosophila melanogaster GN=Lsp1alpha PE=2 SV=3 | 1,07313470463022 | 1,01792190799726 |
| Q9W2J5 | CG44245, isoform A OS=Drosophila melanogaster GN=CG44245 PE=1 SV=3 | 1,09423606984577 | 1,28214322878150 |
| Q9VSU2 | Tequila, isoform G OS=Drosophila melanogaster GN=Tequila PE=1 SV=4 | 1,09963185001446 | 1,13619138628714 |

| | | | |
|---|---|---|---|
| A0A0B4LIT4 | Microtubule-associated protein OS=Drosophila melanogaster GN=tau PE=1 SV=1 | 1,10030490579569 | 1,06557231159362 |
| P81829 | Leucokinin OS=Drosophila melanogaster GN=Lk PE=1 SV=3 | 1,10433665981474 | 1,07244885006969 |
| Q8SZK9 | C-type lectin 27kD, isoform B OS=Drosophila melanogaster GN=Clect27 PE=1 SV=1 | 1,10835717809042 | 1,08678376614207 |
| Q9W4C1 | CG15784, isoform A OS=Drosophila melanogaster GN=CG15784 PE=1 SV=1 | 1,11370049916473 | 1,37795663381011 |
| Q9I7C6 | Cuticular protein 65Ax1 OS=Drosophila melanogaster GN=Cpr65Ax1 PE=1 SV=1 | 1,13158948432814 | 1,02998286621571 |
| P14318 | Muscle-specific protein 20 OS=Drosophila melanogaster GN=Mp20 PE=2 SV=2 | 1,13356352574111 | 1,03068920407114 |
| D1YSG0 | Bent, isoform F OS=Drosophila melanogaster GN=bt PE=1 SV=1 | 1,16156525639703 | 1,12035194036522 |
| A1Z9M5 | CG30069 OS=Drosophila melanogaster GN=CG30069 PE=1 SV=1 | 1,16156525639703 | 1,15250794836490 |
| P83967 | Actin, indirect flight muscle OS=Drosophila melanogaster GN=Act88F PE=1 SV=1 | 1,17120682741814 | 1,03562390973072 |
| Q9V4C1 | CG1674, isoform E OS=Drosophila melanogaster GN=CG1674 PE=1 SV=2 | 1,17184731357334 | 1,15380533607904 |
| Q9VAI9 | Odorant-binding protein 99c, isoform A OS=Drosophila melanogaster GN=Obp99c PE=1 SV=1 | 1,17184731357334 | 1,10030490579569 |
| Q7K1U0 | Activity-regulated cytoskeleton associated protein 1 OS=Drosophila melanogaster GN=Arc1 PE=1 SV=1 | 1,17632277264046 | 1,19597745864259 |
| Q9VYD5 | Branched-chain-amino-acid aminotransferase OS=Drosophila melanogaster GN=CG1673 PE=1 SV=2 | 1,18142064028014 | 1,27262045466299 |
| Q9W306 | CG9691, isoform B OS=Drosophila melanogaster GN=CG9691 PE=1 SV=1 | 1,18396283484259 | 1,03351110236132 |
| Q0E8W5 | Ion transport peptide, isoform C OS=Drosophila melanogaster GN=ITP PE=4 SV=2 | 1,18459768415576 | 1,18903382439002 |
| Q9W5X1 | CG9572, isoform A OS=Drosophila melanogaster GN=CG9572-RA PE=2 SV=1 | 1,18713429147454 | 1,06143060049775 |
| Q9VD23 | Pyruvate kinase OS=Drosophila melanogaster GN=CG7069 PE=3 SV=2 | 1,20037879798403 | 0.867896463992655 |
| A0A0B4LGZ7 | CG45076, isoform H OS=Drosophila melanogaster GN=CG45076 PE=1 SV=1 | 1,20288783347014 | 1,12167855658825 |
| Q7JV70 | Arc2 OS=Drosophila melanogaster GN=Arc2 PE=1 SV=1 | 1,21847121074326 | 1,22712493444993 |
| Q9VSN2 | CG6416-PA, isoform A OS=Drosophila melanogaster GN=Zasp66 PE=1 SV=1 | 1,22589186169034 | 1,39725534559443 |
| Q9W030 | CG32302 OS=Drosophila melanogaster GN=CG32302 PE=1 SV=2 | 1,22774107583913 | 1,23695175858808 |
| X2JB24 | Neuropeptide-like 2, isoform B OS=Drosophila melanogaster GN=Nplp2 PE=4 SV=1 | 1,23388806017428 | 1,19597745864259 |
| Q0E8R1 | FI07211p OS=Drosophila melanogaster GN=vir-1 PE=1 SV=1 | 1,25882039687641 | 1,05727696507325 |
| A0A0B4KEP5 | Mitochondrial ribosomal protein S16, isoform B OS=Drosophila melanogaster GN=mRpS16 PE=4 SV=1 | 1,26963165094044 | 1,52456522107720 |
| Q95TZ7 | GH19182p OS=Drosophila melanogaster GN=Zasp66 PE=1 SV=1 | 1,28806320032532 | 1,19219416528334 |
| A8JQX3 | Curled, isoform D OS=Drosophila melanogaster GN=cu PE=4 SV=1 | 1,29278174922785 | 1,28392177230762 |
| Q8SYQ4 | CG31997, isoform A OS=Drosophila melanogaster GN=CG31997 PE=1 SV=1 | 1,29807173493400 | 1,13553486976119 |
| X2JCI6 | Troponin C at 73F, isoform C OS=Drosophila melanogaster GN=TpnC73F PE=4 SV=1 | 1,31498648546852 | 1,13093086982645 |
| P36188 | Troponin I OS=Drosophila melanogaster GN=wupA PE=2 SV=3 | 1,33799646351502 | 1,41088377719558 |
| P35415 | Paramyosin, long form OS=Drosophila melanogaster GN=Prm PE=1 SV=1 | 1,34936527759421 | 1,36176835941915 |
| P61855 | Adipokinetic hormone OS=Drosophila melanogaster GN=Akh PE=1 SV=1 | 1,34993137336011 | 1,35501626421955 |
| Q0E9J4 | Down syndrome cell adhesion molecule 1, isoform AB OS=Drosophila melanogaster GN=Dscam1 PE=1 SV=1 | 1,35332329116290 | 1,46832264394787 |
| A0A0B4K6I5 | Spartin, isoform B OS=Drosophila melanogaster GN=spartin PE=4 SV=1 | 1,35614381022528 | 0.733788168625182 |
| Q9VEK7 | Cellular repressor of E1A-stimulated genes, isoform A OS=Drosophila melanogaster GN=CREG PE=1 SV=1 | 1,35783347862949 | 1,43935717847426 |
| P18432 | Myosin regulatory light chain 2 OS=Drosophila melanogaster GN=Mlc2 PE=1 SV=2 | 1,36793014143188 | 1,44731470025318 |
| P02515 | Heat shock protein 22 OS=Drosophila melanogaster GN=Hsp22 PE=1 SV=4 | 1,39396527566024 | 1,66266125547509 |
| Q6IL18 | CG43085, isoform A OS=Drosophila melanogaster GN=CG43085 PE=4 SV=1 | 1,40490312214513 | 1,37072227514817 |

| | | | |
|---|---|---|---|
| Q9VTR6 | Pericardin OS=Drosophila melanogaster GN=prc PE=1 SV=2 | 1,40762467556661 | 1,38570712465793 |
| Q9VQU4 | Lectin-24A OS=Drosophila melanogaster GN=lectin-24A PE=2 SV=1 | 1,41467678042689 | 1,42061713897869 |
| A0A0B4LGF8 | Muscle LIM protein at 60A, isoform F OS=Drosophila melanogaster GN=Mlp60A PE=1 SV=1 | 1,42223300068305 | 1,41737997597302 |
| Q8ML70 | Immune-induced peptides OS=Drosophila melanogaster GN=IM10 PE=1 SV=2 | 1,42921422983958 | 1,38349694415537 |
| Q9VQT8 | CG16712, isoform B OS=Drosophila melanogaster GN=IM33 PE=1 SV=1 | 1,44466706684191 | 1,41034110461409 |
| P06742 | Myosin light chain alkali OS=Drosophila melanogaster GN=Mlc1 PE=1 SV=4 | 1,45364926604329 | 1,36569253719753 |
| A1Z9I0 | CG6357 OS=Drosophila melanogaster GN=CG6357 PE=1 SV=1 | 1,49671798793518 | 1,60502015306241 |
| P54398 | Fat body protein 2 OS=Drosophila melanogaster GN=Fbp2 PE=2 SV=2 | 1,51450103627429 | 1,51954190457844 |
| Q9VM18 | Trehalose 6-phosphate phosphatase OS=Drosophila melanogaster GN=CG5177 PE=1 SV=1 | 1,53455968460832 | 1,59263642860658 |
| Q86PF4 | CG10126, isoform B OS=Drosophila melanogaster GN=CG10126-RB PE=1 SV=2 | 1,53505759489484 | 1,58496250072116 |
| P23779 | Cystatin-like protein OS=Drosophila melanogaster GN=Cys PE=2 SV=2 | 1,58784500925428 | 1,50080205305716 |
| A1Z8Z3 | Cuticular protein 49Ag OS=Drosophila melanogaster GN=Cpr49Ag PE=1 SV=1 | 1,59311469648462 | 1,54201035553661 |
| M9PJQ5 | Wings up A, isoform K OS=Drosophila melanogaster GN=wupA PE=1 SV=1 | 1,59645813955899 | 1,55777767139493 |
| P92181 | CG6956-PA OS=Drosophila melanogaster GN=Lcp65Ac PE=1 SV=1 | 1,59836520532364 | 1,66993383566896 |
| M9NE66 | Nervous wreck, isoform B OS=Drosophila melanogaster GN=nwk PE=1 SV=1 | 1,59979385212224 | 1,33571191032046 |
| Q8IN44 | Protein Turandot A OS=Drosophila melanogaster GN=TotA PE=1 SV=1 | 1,64800517981049 | 1,74459168654238 |
| M9NEP1 | Myosin heavy chain, isoform T OS=Drosophila melanogaster GN=Mhc PE=1 SV=1 | 1,67581593117227 | 1,67987414774662 |
| M9PF57 | CG43897, isoform M OS=Drosophila melanogaster GN=CG43897 PE=1 SV=1 | 1,69955163252309 | 1,52155333054509 |
| M9PHR2 | Upheld, isoform O OS=Drosophila melanogaster GN=up PE=1 SV=1 | 1,72071624252113 | 1,68167414179283 |
| A0A0B4KHJ9 | Tropomyosin 2, isoform E OS=Drosophila melanogaster GN=Tm2 PE=1 SV=1 | 1,73855168652023 | 1,79991620298901 |
| M9ND95 | Myosin heavy chain, isoform U OS=Drosophila melanogaster GN=Mhc PE=1 SV=1 | 1,84358192315399 | 1,77610398807316 |
| A0A0B4KEF3 | Larval cuticle protein 3, isoform B OS=Drosophila melanogaster GN=Lcp3 PE=4 SV=1 | 1,92067441104162 | 1,92561971200294 |
| Q9VET0 | Neuropeptide F OS=Drosophila melanogaster GN=NPF PE=1 SV=1 | 1,98258294692259 | 1,77230797495840 |
| P47948 | Troponin C, isoform 2 OS=Drosophila melanogaster GN=TpnC47D PE=2 SV=2 | 2,00072116724365 | 1,96532254836725 |
| P07189 | Larval cuticle protein 4 OS=Drosophila melanogaster GN=Lcp4 PE=1 SV=2 | 2,13520649943067 | 2,19093085956261 |
| Q9W3L4 | CG2233 OS=Drosophila melanogaster GN=CG2233 PE=1 SV=1 | 2,16059754588055 | 2,21038886444540 |
| Q8IP30 | CG4793 OS=Drosophila melanogaster GN=CG4793 PE=3 SV=2 | 2,17568427073100 | 1,97342759800431 |
| Q9VJD7 | CG6639 OS=Drosophila melanogaster GN=SPH93 PE=1 SV=1 | 2,32250505751888 | 2,72900887033786 |
| Q7JZV0 | Cuticular protein 47Eg OS=Drosophila melanogaster GN=Cpr47Eg PE=1 SV=1 | 2,32308178950373 | 2,40735275114004 |
| A0A0B4KF10 | Larval cuticle protein 1, isoform B OS=Drosophila melanogaster GN=Lcp1 PE=4 SV=1 | 2,40381306161659 | 2,45285896471381 |
| P82384 | Larval cuticle protein 9 OS=Drosophila melanogaster GN=Lcp9 PE=1 SV=2 | 2,42304025336151 | 2,28095631383106 |
| P05661 | Myosin heavy chain, muscle OS=Drosophila melanogaster GN=Mhc PE=1 SV=4 | 2,68054940871888 | 2,54695617814141 |
| P92192 | Larval cuticle protein 5 OS=Drosophila melanogaster GN=Lcp65Ab1 PE=1 SV=1 | 2,85319725477036 | 3,03579983742326 |

:rl2
