## Supplemental Table 2 for "An evolutionarily conserved role for separase in the regulation of nuclear lamins"

| Accession | Description | Abundance.Ratio…Sse …Cntrl1 | Abundance.Ratio…Sse…Cn trl2 |
|---|---|---|---|
| Q59E34 | CG8103-PB, isoform B OS=Drosophila melanogaster GN=Mi-2 PE=1 SV=1 | -0.422752464406849 | -0.473931188332412 |
| P08928 | Lamin Dm0 OS=Drosophila melanogaster GN=Lam PE=1 SV=4 | -0.415037499278844 | -0.426625473554056 |
| P02572 | Actin-42A OS=Drosophila melanogaster GN=Act42A PE=1 SV=3 | -0.46394709975979 | -0.461958546666336 |
| D6W4T8 | Coronin OS=Drosophila melanogaster GN=pod1 PE=1 SV=1 | -0.508403405589552 | -0.539519529959989 |
| Q24478 | Centrosome-associated zinc finger protein CP190 OS=Drosophila melanogaster GN=Cp190 PE=1 SV=2 | -0.469929257774916 | -0.481968507397831 |
| M9MS15 | Neurotactin, isoform C OS=Drosophila melanogaster GN=Nrt PE=4 SV=1 | -117.462.139.610.707 | -12.446.850.959.549 |
| P19109 | ATP-dependent RNA helicase p62 OS=Drosophila melanogaster GN=Rm62 PE=1 SV=3 | -0.65744525452268 | -0.63486740654747 |
| Q9VP57 | CG7752-PA OS=Drosophila melanogaster GN=pzg PE=1 SV=1 | -0.403541860441014 | -0.413115187147815 |
| P09180 | 60S ribosomal protein L4 OS=Drosophila melanogaster GN=RpL4 PE=1 SV=2 | -0.537424111957796 | -0.539519529959989 |
| A0A0B4KHI4 | Syncrip, isoform Q OS=Drosophila melanogaster GN=Syp PE=1 SV=1 | -152.699.243.208.383 | -155.639.334.852.439 |
| Q86BM5 | A kinase anchor protein 200, isoform G OS=Drosophila melanogaster GN=Akap200 PE=1 SV=1 | -0.440263475567017 | -0.500217879852688 |
| A0A0B4KFA6 | CD98 heavy chain, isoform D OS=Drosophila melanogaster GN=CD98hc PE=1 SV=1 | -0.422752464406849 | -0.488026018218199 |
| Q9V9U3 | CG1910, isoform A OS=Drosophila melanogaster GN=CG1910 PE=1 SV=1 | -0.465938397578882 | -0.516635639286651 |
| A0A0B4LEV2 | Disc proliferation abnormal, isoform B OS=Drosophila melanogaster GN=dpa PE=1 SV=1 | -0.623709616662948 | -0.655171503002559 |
| Q9VLL3 | A kinase anchor protein 200, isoform A OS=Drosophila melanogaster GN=Akap200 PE=1 SV=3 | -0.673462651860048 | -0.734563103950902 |
| Q9I7K6 | Protein NASP homolog OS=Drosophila melanogaster GN=CG8223 PE=1 SV=1 | -0.401634794676355 | -0.483984852996335 |
| Q24368 | Chromatin-remodeling complex ATPase chain Iswi OS=Drosophila melanogaster GN=Iswi PE=1 SV=1 | -0.673462651860048 | -0.579921884020626 |
| P25171 | Regulator of chromosome condensation OS=Drosophila melanogaster GN=Rcc1 PE=1 SV=2 | -0.454031630894707 | -0.446148031818874 |
| Q86BS3 | Chromator, isoform A OS=Drosophila melanogaster GN=Chro PE=1 SV=1 | -0.50635266602479 | -0.588573754273535 |
| Q9W2U4 | Serine/threonine-protein phosphatase 4 regulatory subunit 2 OS=Drosophila melanogaster GN=PPP4R2r PE=1 SV=2 | -0.457989644463391 | -0.446148031818874 |
| M9PDH1 | Enhancer of bithorax, isoform H OS=Drosophila melanogaster GN=E(bx) PE=1 SV=1 | -0.588573754273535 | -0.552156355637914 |
| M9PBD9 | Ran GTPase activating protein, isoform B OS=Drosophila melanogaster GN=RanGAP PE=1 SV=1 | -0.457989644463391 | -0.446148031818874 |
| P02518 | Heat shock protein 27 OS=Drosophila melanogaster GN=Hsp27 PE=1 SV=2 | -0.891642821909581 | -0.86775220170156 |
| Q9VL18 | Probable elongation factor 1-delta OS=Drosophila melanogaster GN=eEF1delta PE=1 SV=1 | -0.426625473554056 | -0.473931188332412 |
| Q9VNF6 | MTA1-like, isoform A OS=Drosophila melanogaster GN=MTA1-like PE=1 SV=2 | -0.440263475567017 | -0.45600928033515 |
| Q9XYU1 | DNA replication licensing factor Mcm3 OS=Drosophila melanogaster GN=Mcm3 PE=1 SV=1 | -0.543719518489275 | -0.522840788813359 |
| Q0E9M2 | Down syndrome cell adhesion molecule 1, isoform BA OS=Drosophila melanogaster GN=Dscam1 PE=1 SV=1 | -0.471928835421265 | -0.446148031818874 |
| Q9TVM2 | Exportin-1 OS=Drosophila melanogaster GN=emb PE=1 SV=1 | -0.432454552356253 | -0.508403405589552 |
| Q8SWU7 | Obg-like ATPase 1 OS=Drosophila melanogaster GN=CG1354 PE=1 SV=1 | -0.510457064357526 | -0.564904848379903 |
| P00967 | Trifunctional purine biosynthetic protein adenosine-3 OS=Drosophila melanogaster GN=ade3 PE=1 SV=2 | -0.531156057025363 | -0.543719518489275 |
| Q9V431 | Apoptosis inhibitor 5 homolog OS=Drosophila melanogaster GN=Aac11 PE=2 SV=1 | -0.442222328605074 | -0.473931188332412 |
| Q9XZ61 | Ubiquitin carboxyl-terminal hydrolase OS=Drosophila melanogaster GN=Uch-L5 PE=1 SV=1 | -0.415037499278844 | -0.430508908041284 |
| Q9VAA9 | CG7946, isoform A OS=Drosophila melanogaster GN=CG7946 PE=1 SV=1 | -0.448114896528275 | -0.473931188332412 |
| M9PJN2 | Flotillin 2, isoform L OS=Drosophila melanogaster GN=Flo2 PE=1 SV=1 | -0.696657605512669 | -0.708396441969435 |
| Q94920 | Voltage-dependent anion-selective channel OS=Drosophila melanogaster GN=porin PE=1 SV=3 | -0.504304837375931 | -0.512513650651464 |
| O61491 | Flotillin-1 OS=Drosophila melanogaster GN=Flo1 PE=2 SV=1 | -0.512513650651464 | -0.535331732996556 |

| | | | |
|---|---|---|---|
| Q9VMI5 | CG9135 protein OS=Drosophila melanogaster GN=CG9135 PE=1 SV=1 | -0.556393348524385 | -0.678071905112638 |
| P49735 | DNA replication licensing factor Mcm2 OS=Drosophila melanogaster GN=Mcm2 PE=1 SV=1 | -0.77349147019132 | -0.736965594166206 |
| Q9VGS2 | Translationally-controlled tumor protein homolog OS=Drosophila melanogaster GN=Tctp PE=1 SV=1 | -0.554273296650016 | -0.556393348524385 |
| A0A0B4LHX6 | Nucleoside diphosphate kinase OS=Drosophila melanogaster GN=awd PE=1 SV=1 | -0.529072742524873 | -0.442222328605074 |
| Q26365 | ADP,ATP carrier protein OS=Drosophila melanogaster GN=sesB PE=2 SV=4 | -0.610433188237274 | -0.689659879387849 |
| Q8MT06 | Guanine nucleotide-binding protein-like 3 homolog OS=Drosophila melanogaster GN=Ns1 PE=1 SV=2 | -0.446148031818874 | -0.518701058452435 |
| A0A0B4KER0 | Relative of woc, isoform C OS=Drosophila melanogaster GN=row PE=1 SV=1 | -0.424687669312563 | -0.450084446378045 |
| Q76NR6 | Regucalcin, isoform D OS=Drosophila melanogaster GN=regucalcin PE=1 SV=1 | -0.467932447710969 | -0.434402824145775 |
| Q9XZU1 | Exportin-2 OS=Drosophila melanogaster GN=Cas PE=2 SV=2 | -0.573466861883327 | -0.582079992188035 |
| Q9VGP4 | LP08082p OS=Drosophila melanogaster GN=Ranbp9 PE=1 SV=1 | -0.430508908041284 | -0.428565884123491 |
| Q7KK96 | Structural maintenance of chromosomes protein OS=Drosophila melanogaster GN=SMC2 PE=1 SV=1 | -0.575615328461903 | -0.531156057025363 |
| Q9VHC7 | FI21236p1 OS=Drosophila melanogaster GN=rump PE=1 SV=1 | -0.522840788813359 | -0.508403405589552 |
| Q9VRP2 | CG10576, isoform A OS=Drosophila melanogaster GN=CG10576 PE=1 SV=1 | -0.426625473554056 | -0.430508908041284 |
| O18388 | Importin subunit beta OS=Drosophila melanogaster GN=Fs(2)Ket PE=2 SV=2 | -0.407363571393423 | -0.496142467422571 |
| Q8SX89 | Kugelkern, isoform A OS=Drosophila melanogaster GN=kuk PE=1 SV=1 | -0.766111939825723 | -0.732164607902385 |
| A0A0B4KGG9 | C-terminal binding protein, isoform F OS=Drosophila melanogaster GN=CtBP PE=1 SV=1 | -0.416962376203336 | -0.454031630894707 |
| P48159 | 60S ribosomal protein L23 OS=Drosophila melanogaster GN=RpL23 PE=1 SV=2 | -0.488026018218199 | -0.438307278601691 |
| Q9VWG3 | 40S ribosomal protein S10b OS=Drosophila melanogaster GN=RpS10b PE=1 SV=2 | -0.465938397578882 | -0.454031630894707 |
| P08266 | DNA-directed RNA polymerase II subunit RPB2 OS=Drosophila melanogaster GN=RpII140 PE=2 SV=2 | -0.46394709975979 | -0.430508908041284 |
| Q9VGW7 | Arginine methyltransferase 1 OS=Drosophila melanogaster GN=Art1 PE=1 SV=1 | -0.481968507397831 | -0.502259911390907 |
| Q9VQF7 | Bacchus, isoform B OS=Drosophila melanogaster GN=Bacc PE=1 SV=1 | -0.483984852996335 | -0.586405917590825 |
| X2JAF4 | Fasciclin 2, isoform G OS=Drosophila melanogaster GN=Fas2 PE=1 SV=1 | -0.586405917590825 | -0.590744853315162 |
| Q9V491 | Plexin A, isoform A OS=Drosophila melanogaster GN=PlexA PE=1 SV=1 | -0.483984852996335 | -0.524915117051217 |
| Q9VYV4 | Amun, isoform A OS=Drosophila melanogaster GN=Amun PE=1 SV=1 | -0.45205668870965 | -0.492078535042672 |
| Q9VMR6 | CG12512 OS=Drosophila melanogaster GN=CG12512-RA PE=1 SV=1 | -0.520769438793664 | -0.508403405589552 |
| P11997 | Larval serum protein 1 gamma chain OS=Drosophila melanogaster GN=Lsp1gamma PE=2 SV=2 | -0.625934281777462 | -0.715485866755755 |
| Q9XYU0 | DNA replication licensing factor Mcm7 OS=Drosophila melanogaster GN=Mcm7 PE=1 SV=1 | -0.751465163861321 | -0.729770092762002 |
| P08111 | Lethal(2) giant larvae protein OS=Drosophila melanogaster GN=l(2)gl PE=1 SV=2 | -0.477944250839036 | -0.44418384493836 |
| A0A0C4FEI8 | Granny smith, isoform F OS=Drosophila melanogaster GN=grsm PE=1 SV=1 | -0.401634794676355 | -0.424687669312563 |
| Q9V461 | DNA replication licensing factor Mcm6 OS=Drosophila melanogaster GN=Mcm6 PE=1 SV=1 | -0.666576266274808 | -0.706041020971306 |
| Q9VGW6 | DNA replication licensing factor Mcm5 OS=Drosophila melanogaster GN=Mcm5 PE=1 SV=1 | -0.592919224549499 | -0.541617995843987 |
| A1Z6S7 | FI02061p OS=Drosophila melanogaster GN=vimar PE=1 SV=1 | -0.434402824145775 | -0.477944250839036 |
| Q9V406 | Activator protein 4 OS=Drosophila melanogaster GN=crp PE=1 SV=1 | -0.529072742524873 | -0.573466861883327 |
| Q9VA37 | DJ-1 beta OS=Drosophila melanogaster GN=dj-1beta PE=1 SV=2 | -0.448114896528275 | -0.522840788813359 |
| Q9V3I1 | BcDNA.LD08534 OS=Drosophila melanogaster GN=dUTPase PE=1 SV=1 | -0.42081985187285 | -0.432454552356253 |
| Q7K180 | LD02709p OS=Drosophila melanogaster GN=Map60 PE=1 SV=1 | -0.614845103115656 | -0.625934281777462 |
| Q59E36 | REST corepressor OS=Drosophila melanogaster GN=CoRest PE=1 SV=2 | -0.720231578406405 | -0.533242384273829 |

| | | | |
|---|---|---|---|
| P54397 | 39 kDa FK506-binding nuclear protein OS=Drosophila melanogaster GN=FK506-bp1 PE=1 SV=2 | -0.535331732996556 | -0.502259911390907 |
| E1JIL4 | Chromatin assembly factor 1 subunit, isoform B OS=Drosophila melanogaster GN=Caf1-55 PE=1 SV=1 | -0.680382065799839 | -0.678071905112638 |
| Q9W3W8 | 60S ribosomal protein L17 OS=Drosophila melanogaster GN=RpL17 PE=1 SV=1 | -0.403541860441014 | -0.41119543298445 |
| A4V441 | Singed, isoform B OS=Drosophila melanogaster GN=sn PE=1 SV=1 | -0.475936324222789 | -0.554273296650016 |
| Q8IMX4 | FI04408p OS=Drosophila melanogaster GN=Rox8 PE=1 SV=1 | -0.592919224549499 | -0.623709616662948 |
| Q7KSM5 | Aos1 OS=Drosophila melanogaster GN=Aos1 PE=1 SV=1 | -0.488026018218199 | -0.469929257774916 |
| Q7KNM2 | SUMO-conjugating enzyme OS=Drosophila melanogaster GN=lwr PE=1 SV=1 | -0.623709616662948 | -0.694321256757713 |
| A0A0B4KI24 | Nucleoplasmin, isoform B OS=Drosophila melanogaster GN=Nlp PE=4 SV=1 | -0.940644722383579 | -10.086.822.430.998 |
| Q9V3A7 | Structural maintenance of chromosomes protein OS=Drosophila melanogaster GN=glu PE=1 SV=1 | -0.496142467422571 | -0.465938397578882 |
| M9PI41 | Sticky, isoform B OS=Drosophila melanogaster GN=sti PE=1 SV=1 | -0.508403405589552 | -0.500217879852688 |
| A1Z6M0 | Brahma associated protein 170kD OS=Drosophila melanogaster GN=Bap170 PE=1 SV=1 | -0.558516520417355 | -0.567040592723894 |
| P84051 | Histone H2A OS=Drosophila melanogaster GN=His2A PE=1 SV=2 | -116.812.275.880.833 | -114.241.704.461.585 |
| Q9VSL4 | Glutathione S transferase O2, isoform B OS=Drosophila melanogaster GN=GstO2 PE=1 SV=2 | -0.475936324222789 | -0.481968507397831 |
| M9PFF0 | 60S ribosomal protein L13 OS=Drosophila melanogaster GN=RpL13 PE=1 SV=1 | -0.46394709975979 | -0.486004020632987 |
| Q9VRP3 | AT08565p OS=Drosophila melanogaster GN=Txl PE=1 SV=1 | -0.575615328461903 | -0.586405917590825 |
| P17917 | Proliferating cell nuclear antigen OS=Drosophila melanogaster GN=PCNA PE=1 SV=2 | -0.500217879852688 | -0.545824106814198 |
| Q6AWJ9 | Tyrosine-protein kinase-like otk OS=Drosophila melanogaster GN=otk PE=1 SV=1 | -0.940644722383579 | -105.889.368.905.357 |
| Q9VXK5 | Putative rRNA methyltransferase OS=Drosophila melanogaster GN=CG8939 PE=1 SV=1 | -0.520769438793664 | -0.597277823154395 |
| A4UZI6 | Rho1, isoform C OS=Drosophila melanogaster GN=Rho1 PE=1 SV=1 | -0.510457064357526 | -0.490050853695689 |
| P05205 | Heterochromatin protein 1 OS=Drosophila melanogaster GN=Su(var)205 PE=1 SV=2 | -0.422752464406849 | -0.442222328605074 |
| Q24298 | DE-cadherin OS=Drosophila melanogaster GN=shg PE=1 SV=2 | -0.899695094204314 | -0.926865295369785 |
| O61345 | Protein penguin OS=Drosophila melanogaster GN=peng PE=2 SV=1 | -0.459972730742493 | -0.516635639286651 |
| Q9VH48 | Probable histone-arginine methyltransferase CARMER OS=Drosophila melanogaster GN=Art4 PE=1 SV=1 | -0.467932447710969 | -0.467932447710969 |
| O61305 | DEAD-box helicase Dbp80 OS=Drosophila melanogaster GN=Dbp80 PE=1 SV=1 | -0.577766999316952 | -0.63486740654747 |
| Q86BY9 | Protein rigor mortis OS=Drosophila melanogaster GN=rig PE=1 SV=1 | -0.461958546666336 | -0.496142467422571 |
| P55935 | 40S ribosomal protein S9 OS=Drosophila melanogaster GN=RpS9 PE=1 SV=2 | -0.531156057025363 | -0.610433188237274 |
| Q9VCD8 | Structural maintenance of chromosomes protein OS=Drosophila melanogaster GN=SMC1 PE=1 SV=1 | -0.586405917590825 | -0.477944250839036 |
| M9NGB9 | Mushroom-body expressed, isoform H OS=Drosophila melanogaster GN=mub PE=1 SV=1 | -0.579921884020626 | -0.535331732996556 |
| M9PGG4 | Armadillo, isoform F OS=Drosophila melanogaster GN=arm PE=4 SV=1 | -0.467932447710969 | -0.44418384493836 |
| Q9V9J3 | Tyrosine-protein kinase Src42A OS=Drosophila melanogaster GN=Src42A PE=2 SV=1 | -0.612637459164004 | -0.691988685447822 |
| P52304 | Serine/threonine-protein kinase polo OS=Drosophila melanogaster GN=polo PE=1 SV=2 | -0.720231578406405 | -0.768567591552035 |
| B7Z0E2 | FI18812p1 OS=Drosophila melanogaster GN=Unr PE=1 SV=1 | -0.428565884123491 | -0.430508908041284 |
| Q9VCI7 | LD02979p OS=Drosophila melanogaster GN=RanBP3 PE=1 SV=1 | -0.465938397578882 | -0.50635266602479 |
| Q9VZE4 | CG1316 OS=Drosophila melanogaster GN=CG1316 PE=1 SV=2 | -0.508403405589552 | -0.560642821525743 |
| P34739 | Transcription termination factor 2 OS=Drosophila melanogaster GN=lds PE=1 SV=2 | -0.573466861883327 | -0.687334826441606 |
| Q9VD51 | Probable ATP-dependent RNA helicase pitchoune OS=Drosophila melanogaster GN=pit PE=2 SV=2 | -0.471928835421265 | -0.504304837375931 |
| Q9VXG4 | Annexin B11 OS=Drosophila melanogaster GN=AnxB11 PE=2 SV=2 | -0.416962376203336 | -0.440263475567017 |

| | | | |
|---|---|---|---|
| Q9W2I2 | CG9752 OS=Drosophila melanogaster GN=CG9752 PE=1 SV=1 | -0.539519529959989 | -0.575615328461903 |
| P16620 | Tyrosine-protein phosphatase 69D OS=Drosophila melanogaster GN=Ptp69D PE=1 SV=2 | -0.706041020971306 | -0.67116353577046 |
| Q9I7K0 | Microtubule-associated protein Jupiter OS=Drosophila melanogaster GN=Jupiter PE=1 SV=2 | -0.577766999316952 | -0.552156355637914 |
| Q9V3E9 | FI17138p1 OS=Drosophila melanogaster GN=spag PE=1 SV=1 | -0.403541860441014 | -0.442222328605074 |
| Q7JXU4 | Nuclear GTP binding protein OS=Drosophila melanogaster GN=Ns2 PE=1 SV=1 | -0.564904848379903 | -0.575615328461903 |
| P02283 | Histone H2B OS=Drosophila melanogaster GN=His2B PE=1 SV=2 | -13.621.579.396.759 | -141.888.982.477.445 |
| Q9VCR2 | Aminoacylase-1 OS=Drosophila melanogaster GN=CG6726 PE=1 SV=2 | -0.481968507397831 | -0.492078535042672 |
| E1JGK5 | Mapmodulin, isoform D OS=Drosophila melanogaster GN=Mapmodulin PE=1 SV=1 | -0.873027143742234 | -0.883635243308215 |
| Q9VHJ7 | CG11964 OS=Drosophila melanogaster GN=CG11964 PE=1 SV=1 | -0.524915117051217 | -0.475936324222789 |
| Q9W1H4 | DNA ligase 1 OS=Drosophila melanogaster GN=DNA-ligI PE=1 SV=2 | -0.727379545337008 | -0.739372091873301 |
| A1Z6I7 | Bub1-related kinase OS=Drosophila melanogaster GN=BubR1 PE=1 SV=1 | -0.477944250839036 | -0.496142467422571 |
| Q6NP69 | GST-containing FLYWCH zinc-finger protein OS=Drosophila melanogaster GN=gfzf PE=1 SV=1 | -0.454031630894707 | -0.459972730742493 |
| Q8INH6 | CG32473, isoform C OS=Drosophila melanogaster GN=CG32473-RC PE=1 SV=1 | -0.432454552356253 | -0.440263475567017 |
| Q0KIB3 | CG2051, isoform A OS=Drosophila melanogaster GN=CG2051 PE=1 SV=1 | -0.524915117051217 | -0.547931769776189 |
| Q9VM49 | CG11266, isoform A OS=Drosophila melanogaster GN=Caper PE=1 SV=1 | -0.504304837375931 | -0.50635266602479 |
| P55841 | 60S ribosomal protein L14 OS=Drosophila melanogaster GN=RpL14 PE=1 SV=1 | -0.590744853315162 | -0.446148031818874 |
| Q94516 | ATP synthase subunit b, mitochondrial OS=Drosophila melanogaster GN=ATPsynB PE=2 SV=2 | -0.469929257774916 | -0.44418384493836 |
| Q9VZY0 | LD45195p OS=Drosophila melanogaster GN=Non2 PE=1 SV=1 | -0.617056130431009 | -0.63262893435147 |
| Q9VVI1 | CG7564, isoform B OS=Drosophila melanogaster GN=CG7564 PE=1 SV=1 | -0.44418384493836 | -0.45600928033515 |
| P53997 | Protein SET OS=Drosophila melanogaster GN=Set PE=1 SV=2 | -0.682695931638085 | -0.763660460831626 |
| Q8T8R1 | CCHC-type zinc finger protein CG3800 OS=Drosophila melanogaster GN=CG3800 PE=1 SV=1 | -0.430508908041284 | -0.434402824145775 |
| A0A0C4DHF6 | Dystroglycan, isoform D OS=Drosophila melanogaster GN=Dg PE=1 SV=1 | -0.650634722405787 | -0.648371670897218 |
| Q9VJ43 | LD10783p OS=Drosophila melanogaster GN=ScpX PE=1 SV=1 | -0.512513650651464 | -0.569179503480228 |
| Q7JQN4 | LD15481p OS=Drosophila melanogaster GN=Rs1 PE=1 SV=1 | -0.442222328605074 | -0.430508908041284 |
| Q9V9T4 | ATP-dependent chromatin assembly factor large subunit OS=Drosophila melanogaster GN=Acf PE=1 SV=1 | -0.689659879387849 | -0.703689439291908 |
| Q9VIP9 | Condensin complex subunit 2 OS=Drosophila melanogaster GN=barr PE=1 SV=1 | -0.701341684435485 | -0.685013514531485 |
| M9PCH8 | Barrier to autointegration factor, isoform B OS=Drosophila melanogaster GN=baf PE=4 SV=1 | -0.612637459164004 | -0.601649629654035 |
| Q9VA18 | Lethal (3) 03670 OS=Drosophila melanogaster GN=l(3)03670 PE=1 SV=1 | -0.496142467422571 | -0.438307278601691 |
| Q9VQI7 | CG3083-PA OS=Drosophila melanogaster GN=Prx6005 PE=1 SV=1 | -0.508403405589552 | -0.518701058452435 |
| P84040 | Histone H4 OS=Drosophila melanogaster GN=His4 PE=1 SV=2 | -0.691988685447822 | -0.696657605512669 |
| P39018 | 40S ribosomal protein S19a OS=Drosophila melanogaster GN=RpS19a PE=1 SV=3 | -0.512513650651464 | -0.496142467422571 |
| P35875 | Poly [ADP-ribose] polymerase OS=Drosophila melanogaster GN=Parp PE=2 SV=1 | -0.479954975960182 | -0.481968507397831 |
| M9MRC9 | Ribosomal protein L27A, isoform C OS=Drosophila melanogaster GN=RpL27A PE=3 SV=1 | -0.586405917590825 | -0.554273296650016 |
| A0A0B4LFD9 | Ribosomal protein S23, isoform B OS=Drosophila melanogaster GN=RpS23 PE=3 SV=1 | -0.547931769776189 | -0.461958546666336 |
| Q494K2 | Ctf4 OS=Drosophila melanogaster GN=Ctf4 PE=1 SV=1 | -0.488026018218199 | -0.46394709975979 |
| Q9V4S8 | COP9 signalosome complex subunit 7 OS=Drosophila melanogaster GN=CSN7 PE=1 SV=2 | -0.486004020632987 | -0.479954975960182 |
| Q9VSI1 | CG7182, isoform A OS=Drosophila melanogaster GN=CG7182 PE=1 SV=2 | -0.545824106814198 | -0.432454552356253 |

| | | | |
|---|---|---|---|
| Q9VIP0 | Nessun dorma, isoform A OS=Drosophila melanogaster GN=nesd PE=1 SV=1 | -0.440263475567017 | -0.479954975960182 |
| Q9VEX6 | AAA family protein Bor OS=Drosophila melanogaster GN=bor PE=1 SV=2 | -0.486004020632987 | -0.520769438793664 |
| Q9V5M6 | Longitudinals lacking protein, isoforms J/P/Q/S/Z OS=Drosophila melanogaster GN=lola PE=1 SV=4 | -0.416962376203336 | -0.529072742524873 |
| Q7KQZ4 | Longitudinals lacking protein, isoforms A/B/D/L OS=Drosophila melanogaster GN=lola PE=1 SV=1 | -0.531156057025363 | -0.573466861883327 |
| M9MSJ1 | Brain tumor, isoform E OS=Drosophila melanogaster GN=brat PE=1 SV=1 | -0.780908941753803 | -0.746615764199925 |
| Q9VK58 | Pih1D1, isoform D OS=Drosophila melanogaster GN=Pih1D1 PE=1 SV=5 | -0.42081985187285 | -0.422752464406849 |
| Q9VX15 | CG8142 OS=Drosophila melanogaster GN=CG8142 PE=1 SV=2 | -0.56277226108709 | -0.524915117051217 |
| M9PET3 | Simjang, isoform D OS=Drosophila melanogaster GN=simj PE=1 SV=1 | -0.45205668870965 | -0.446148031818874 |
| Q9U9Q1 | Replication factor C 38kD subunit, isoform A OS=Drosophila melanogaster GN=RfC38 PE=1 SV=1 | -0.481968507397831 | -0.504304837375931 |
| Q9VKW3 | CG5313-PA OS=Drosophila melanogaster GN=RfC3 PE=1 SV=2 | -0.45600928033515 | -0.550042516371997 |
| Q7KVQ0 | Probable H/ACA ribonucleoprotein complex subunit 1 OS=Drosophila melanogaster GN=CG4038 PE=2 SV=1 | -0.49817873457909 | -0.471928835421265 |
| P39769 | Polyhomeotic-proximal chromatin protein OS=Drosophila melanogaster GN=ph-p PE=1 SV=2 | -0.678071905112638 | -0.79836613883035 |
| Q9VJ44 | CG17597, isoform B OS=Drosophila melanogaster GN=CG17597 PE=1 SV=2 | -0.788364746672851 | -0.788364746672851 |
| A8DYK5 | CG4266, isoform B OS=Drosophila melanogaster GN=CG4266 PE=1 SV=1 | -0.481968507397831 | -0.436353730515936 |
| P08985 | Histone H2A.v OS=Drosophila melanogaster GN=His2Av PE=1 SV=2 | -0.778432211461591 | -0.741782610463982 |
| A1Z6H7 | Gp210 ortholog, isoform A OS=Drosophila melanogaster GN=Gp210 PE=1 SV=1 | -0.405451450449646 | -0.450084446378045 |
| Q9VXP4 | Platelet-activating factor acetylhydrolase IB subunit beta homolog OS=Drosophila melanogaster GN=Paf-AHalpha PE=1 SV=1 | -0.575615328461903 | -0.569179503480228 |
| Q9VI15 | CG10286 OS=Drosophila melanogaster GN=CG10286 PE=1 SV=1 | -0.430508908041284 | -0.416962376203336 |
| A1Z920 | FI24025p1 OS=Drosophila melanogaster GN=fra PE=1 SV=1 | -0.569179503480228 | -0.63262893435147 |
| E1JJE6 | CG42492, isoform C OS=Drosophila melanogaster GN=CG42492 PE=1 SV=1 | -0.595096877854869 | -0.795859283219775 |
| P20353 | G protein alpha i subunit OS=Drosophila melanogaster GN=Galphai PE=1 SV=2 | -0.761213140412883 | -0.783389931257558 |
| Q7KJ08 | Alk, isoform A OS=Drosophila melanogaster GN=Alk PE=1 SV=1 | -0.486004020632987 | -0.475936324222789 |
| M9NEY8 | DNA N6-methyl adenine demethylase OS=Drosophila melanogaster GN=Tet PE=1 SV=1 | -0.461958546666336 | -0.415037499278844 |
| P05812 | Heat shock protein 67B1 OS=Drosophila melanogaster GN=Hsp67Ba PE=3 SV=1 | -0.479954975960182 | -0.461958546666336 |
| Q76NQ0 | Dolichyl-diphosphooligosaccharide--protein glycosyltransferase subunit 1 OS=Drosophila melanogaster GN=CG33303 PE=1 S | -0.401634794676355 | -0.45205668870965 |
| Q86B87 | Modifier of mdg4 OS=Drosophila melanogaster GN=mod(mdg4) PE=1 SV=1 | -0.601649629654035 | -0.608232280044003 |
| Q9W542 | LD07342p OS=Drosophila melanogaster GN=mip130 PE=1 SV=3 | -0.440263475567017 | -0.465938397578882 |
| M9PCL7 | Sema-1a, isoform G OS=Drosophila melanogaster GN=Sema-1a PE=1 SV=1 | -0.763660460831626 | -0.706041020971306 |
| Q9VL96 | Pescadillo homolog OS=Drosophila melanogaster GN=CG4364 PE=1 SV=1 | -0.502259911390907 | -0.560642821525743 |
| Q27601 | Amidophosphoribosyltransferase OS=Drosophila melanogaster GN=Prat PE=1 SV=2 | -0.520769438793664 | -0.550042516371997 |
| Q9VD52 | CG6015 OS=Drosophila melanogaster GN=CG6015 PE=1 SV=1 | -0.405451450449646 | -0.45205668870965 |
| Q9VQJ8 | Protein-lysine N-methyltransferase CG9643 OS=Drosophila melanogaster GN=CG9643 PE=1 SV=1 | -0.694321256757713 | -0.63710935733414 |
| Q9VCF8 | CG5854 OS=Drosophila melanogaster GN=CG5854 PE=1 SV=1 | -0.650634722405787 | -0.701341684435485 |
| P20348 | Sex-regulated protein janus-A OS=Drosophila melanogaster GN=janA PE=2 SV=2 | -0.662003536484984 | -0.588573754273535 |
| Q9V4D4 | CG2009-PA OS=Drosophila melanogaster GN=bip2 PE=1 SV=1 | -0.535331732996556 | -0.490050853695689 |
| P29617 | Homeobox protein prospero OS=Drosophila melanogaster GN=pros PE=1 SV=3 | -0.710755714843358 | -0.678071905112638 |
| Q9W523 | Polyhomeotic distal, isoform A OS=Drosophila melanogaster GN=ph-d PE=1 SV=3 | -0.708396441969435 | -0.873027143742234 |

| | | | |
|---|---|---|---|
| P23572 | Cyclin-dependent kinase 1 OS=Drosophila melanogaster GN=Cdk1 PE=1 SV=1 | -0.614845103115656 | -0.734563103950902 |
| Q9W260 | GH03113p OS=Drosophila melanogaster GN=wrapper PE=1 SV=2 | -0.741782610463982 | -0.567040592723894 |
| E1JJB2 | Broad, isoform P OS=Drosophila melanogaster GN=br PE=1 SV=2 | -0.808437349299244 | -0.768567591552035 |
| Q9W5W6 | CG9578 OS=Drosophila melanogaster GN=CG9578 PE=1 SV=2 | -0.494109070270043 | -0.488026018218199 |
| Q9VCH9 | CG10375, isoform A OS=Drosophila melanogaster GN=CG10375 PE=1 SV=1 | -0.595096877854869 | -0.529072742524873 |
| B7YZV6 | Scm-related gene containing four mbt domains, isoform C OS=Drosophila melanogaster GN=Sfmbt PE=1 SV=1 | -0.475936324222789 | -0.459972730742493 |
| Q9VE69 | CG31122, isoform A OS=Drosophila melanogaster GN=CG31122 PE=1 SV=3 | -0.481968507397831 | -0.50635266602479 |
| Q9VAJ2 | Bub3, isoform A OS=Drosophila melanogaster GN=Bub3 PE=1 SV=1 | -0.49817873457909 | -0.539519529959989 |
| Q7K7A9 | Flap endonuclease 1 OS=Drosophila melanogaster GN=Fen1 PE=2 SV=1 | -0.552156355637914 | -0.610433188237274 |
| Q9VDR1 | Mediator of RNA polymerase II transcription subunit 25 OS=Drosophila melanogaster GN=MED25 PE=2 SV=1 | -0.579921884020626 | -0.533242384273829 |
| Q9VBU4 | CG11858 OS=Drosophila melanogaster GN=CG11858 PE=1 SV=1 | -0.643856189774725 | -0.575615328461903 |
| Q9VND7 | CG2931 OS=Drosophila melanogaster GN=CG2931 PE=1 SV=1 | -0.481968507397831 | -0.543719518489275 |
| Q9V597 | 60S ribosomal protein L31 OS=Drosophila melanogaster GN=RpL31 PE=1 SV=1 | -0.465938397578882 | -0.438307278601691 |
| Q9VUB8 | CG6513-PA, isoform A OS=Drosophila melanogaster GN=endos PE=1 SV=1 | -0.401634794676355 | -0.42081985187285 |
| Q7KSQ0 | LD46175p OS=Drosophila melanogaster GN=sea PE=1 SV=1 | -0.494109070270043 | -0.450084446378045 |
| Q9XZ14 | CG9634, isoform A OS=Drosophila melanogaster GN=goe PE=1 SV=1 | -0.49817873457909 | -0.577766999316952 |
| M9PBM1 | U2 small nuclear riboprotein auxiliary factor 38, isoform B OS=Drosophila melanogaster GN=U2af38 PE=4 SV=1 | -0.465938397578882 | -0.405451450449646 |
| Q9VUL8 | LD41491p OS=Drosophila melanogaster GN=Pex3 PE=1 SV=1 | -0.788364746672851 | -0.744197163397282 |
| Q9VFR0 | Protein BCCIP homolog OS=Drosophila melanogaster GN=CG9286 PE=2 SV=2 | -0.475936324222789 | -0.422752464406849 |
| Q7KMH5 | BcDNA.LD29892 OS=Drosophila melanogaster GN=Smyd4-4 PE=1 SV=1 | -0.543719518489275 | -0.62148837674627 |
| P48588 | 40S ribosomal protein S25 OS=Drosophila melanogaster GN=RpS25 PE=1 SV=3 | -0.426625473554056 | -0.520769438793664 |
| E1JJF3 | Innexin OS=Drosophila melanogaster GN=Inx2 PE=1 SV=1 | -0.727379545337008 | -0.849440323424619 |
| Q9W0B6 | CG17249 OS=Drosophila melanogaster GN=CG17249 PE=1 SV=1 | -0.518701058452435 | -0.529072742524873 |
| Q9VHI1 | Hyrax OS=Drosophila melanogaster GN=hyx PE=1 SV=1 | -0.486004020632987 | -0.401634794676355 |
| Q9VLV5 | Probable small nuclear ribonucleoprotein E OS=Drosophila melanogaster GN=SmE PE=1 SV=1 | -0.442222328605074 | -0.625934281777462 |
| Q9VAJ1 | Condensin complex subunit 1 OS=Drosophila melanogaster GN=Cap-D2 PE=1 SV=1 | -0.586405917590825 | -0.550042516371997 |
| Q9V3Y4 | CG6851-PA, isoform A OS=Drosophila melanogaster GN=Mtch PE=1 SV=1 | -0.595096877854869 | -0.42081985187285 |
| Q8IRE4 | tRNA (guanine(37)-N1)-methyltransferase OS=Drosophila melanogaster GN=CG32281 PE=2 SV=2 | -0.56277226108709 | -0.617056130431009 |
| Q9V535 | RNA-binding protein 8A OS=Drosophila melanogaster GN=tsu PE=1 SV=1 | -0.405451450449646 | -0.465938397578882 |
| D0IQG7 | Chitinase 2, isoform B OS=Drosophila melanogaster GN=Cht2 PE=1 SV=1 | -0.524915117051217 | -0.430508908041284 |
| Q7K3D8 | DMAP1 OS=Drosophila melanogaster GN=DMAP1 PE=1 SV=1 | -0.537424111957796 | -0.442222328605074 |
| X2JCI2 | Tyrosine-protein kinase OS=Drosophila melanogaster GN=Src64B PE=3 SV=1 | -0.610433188237274 | -0.722610301189136 |
| O18338 | CG8287-PA OS=Drosophila melanogaster GN=Rab8 PE=1 SV=1 | -0.603840510926846 | -0.50635266602479 |
| Q7K2X8 | Nucleoporin at 44A, isoform A OS=Drosophila melanogaster GN=Nup44A PE=1 SV=1 | -0.552156355637914 | -0.459972730742493 |
| Q9VRQ2 | Mad2 OS=Drosophila melanogaster GN=mad2 PE=1 SV=1 | -0.628162382669579 | -0.623709616662948 |
| B7Z060 | TBP-associated factor 4, isoform E OS=Drosophila melanogaster GN=Taf4 PE=1 SV=1 | -0.409278229990159 | -0.403541860441014 |
| Q9VJY6 | 60S ribosomal protein L24 OS=Drosophila melanogaster GN=RpL24 PE=1 SV=1 | -0.567040592723894 | -0.442222328605074 |

| | | | |
|---|---|---|---|
| P20193 | Protein suppressor of variegation 3-7 OS=Drosophila melanogaster GN=Su(var)3-7 PE=1 SV=4 | -0.450084446378045 | -0.428565884123491 |
| A0A0B4LF54 | Cytochrome P450-6a17, isoform B OS=Drosophila melanogaster GN=Cyp6a17 PE=1 SV=1 | -0.687334826441606 | -0.641603738043346 |
| Q7JVG2 | Aps, isoform A OS=Drosophila melanogaster GN=Aps PE=1 SV=1 | -0.457989644463391 | -0.41888982477445 |
| A1Z729 | CG2064 OS=Drosophila melanogaster GN=CG2064 PE=1 SV=1 | -0.481968507397831 | -0.564904848379903 |
| M9PEK6 | Mushroom body defect, isoform L OS=Drosophila melanogaster GN=mud PE=1 SV=1 | -0.703689439291908 | -0.694321256757713 |
| Q9W0G1 | Tyrosine-protein phosphatase non-receptor type 61F OS=Drosophila melanogaster GN=Ptp61F PE=1 SV=1 | -0.514573172829758 | -0.603840510926846 |
| Q9Y113 | Negative elongation factor B OS=Drosophila melanogaster GN=NELF-B PE=1 SV=1 | -0.481968507397831 | -0.469929257774916 |
| Q9VZL1 | LP07226p OS=Drosophila melanogaster GN=mge PE=1 SV=1 | -0.689659879387849 | -0.586405917590825 |
| Q9VMP9 | Glucosamine-6-phosphate isomerase OS=Drosophila melanogaster GN=Gnpda1 PE=2 SV=1 | -0.467932447710969 | -0.512513650651464 |
| Q7JVZ8 | Glutathione S transferase E11, isoform A OS=Drosophila melanogaster GN=GstE11 PE=1 SV=1 | -0.442222328605074 | -0.407363571393423 |
| Q9Y124 | BcDNA.GH08385 OS=Drosophila melanogaster GN=BcDNA.GH08385 PE=1 SV=1 | -0.428565884123491 | -0.46394709975979 |
| O62609 | Mothers against decapentaplegic homolog OS=Drosophila melanogaster GN=Med PE=1 SV=1 | -0.416962376203336 | -0.45600928033515 |
| Q9VJI7 | CG13277-PA OS=Drosophila melanogaster GN=LSm7 PE=1 SV=1 | -0.403541860441014 | -0.461958546666336 |
| A8Y560 | Ribosomal protein L15 OS=Drosophila melanogaster GN=RpL15 PE=1 SV=2 | -0.50635266602479 | -0.475936324222789 |
| Q9W1I6 | LD12035p OS=Drosophila melanogaster GN=ytr PE=1 SV=1 | -0.72499295250013 | -0.698997743967186 |
| O96692 | RE63021p OS=Drosophila melanogaster GN=Rap2l PE=1 SV=1 | -0.424687669312563 | -0.477944250839036 |
| P48611 | 6-pyruvoyl tetrahydrobiopterin synthase OS=Drosophila melanogaster GN=pr PE=1 SV=1 | -0.428565884123491 | -0.475936324222789 |
| Q94513 | Boundary element associated factor OS=Drosophila melanogaster GN=BEAF-32 PE=1 SV=1 | -0.573466861883327 | -0.438307278601691 |
| Q9VWQ7 | CG6891-PA, isoform A OS=Drosophila melanogaster GN=CG6891 PE=1 SV=1 | -0.573466861883327 | -0.590744853315162 |
| Q9V3F8 | Pyrroline-5-carboxylate reductase OS=Drosophila melanogaster GN=P5cr PE=1 SV=1 | -0.556393348524385 | -0.512513650651464 |
| Q9VXI6 | Cytochrome b-c1 complex subunit 7 OS=Drosophila melanogaster GN=UQCR-14 PE=1 SV=1 | -0.426625473554056 | -0.529072742524873 |
| Q7K0L8 | FLASH ortholog, isoform A OS=Drosophila melanogaster GN=FLASH PE=1 SV=1 | -0.524915117051217 | -0.422752464406849 |
| O01382 | Caspase OS=Drosophila melanogaster GN=Drice PE=1 SV=2 | -0.50635266602479 | -0.597277823154395 |
| Q9W3D1 | Caf1-180 OS=Drosophila melanogaster GN=Caf1-180 PE=1 SV=1 | -0.44418384493836 | -0.407363571393423 |
| A0A0B4KHJ7 | MRG15, isoform B OS=Drosophila melanogaster GN=MRG15 PE=4 SV=1 | -0.490050853695689 | -0.41888982477445 |
| Q9VK57 | LD15349p OS=Drosophila melanogaster GN=Pih1D1 PE=1 SV=1 | -0.504304837375931 | -0.529072742524873 |
| M9PIC9 | Spc105-related, isoform B OS=Drosophila melanogaster GN=Spc105R PE=1 SV=1 | -0.537424111957796 | -0.539519529959989 |
| P52654 | Transcription initiation factor IIA subunit 1 OS=Drosophila melanogaster GN=TfIIA-L PE=1 SV=2 | -0.446148031818874 | -0.41888982477445 |
| Q9VDQ3 | CG4936 OS=Drosophila melanogaster GN=CG4936 PE=1 SV=1 | -0.533242384273829 | -0.567040592723894 |
| Q9W088 | DNA polymerase delta small subunit OS=Drosophila melanogaster GN=CG12018 PE=2 SV=1 | -0.83650126771712 | -0.883635243308215 |
| A1Z6M6 | DNA polymerase interacting tpr containing protein of 47kD, isoform A OS=Drosophila melanogaster GN=Dpit47 PE=1 SV=1 | -0.606034724339757 | -0.573466861883327 |
| Q9VZ62 | CG11207-PA OS=Drosophila melanogaster GN=feo PE=1 SV=1 | -0.595096877854869 | -0.639354797539784 |
| Q9VWA8 | Protein FRG1 homolog OS=Drosophila melanogaster GN=FRG1 PE=2 SV=1 | -0.42081985187285 | -0.465938397578882 |
| Q24317 | DNA primase small subunit OS=Drosophila melanogaster GN=DNApol-alpha50 PE=2 SV=2 | -0.915935735211525 | -0.785875194647153 |
| A1Z987 | Cap-G, isoform F OS=Drosophila melanogaster GN=Cap-G PE=1 SV=2 | -0.694321256757713 | -0.710755714843358 |
| Q9VBH7 | CG14544 OS=Drosophila melanogaster GN=CG14544 PE=1 SV=1 | -0.477944250839036 | -0.588573754273535 |
| Q9VEX5 | Protein asunder OS=Drosophila melanogaster GN=Asun PE=1 SV=1 | -0.547931769776189 | -0.483984852996335 |

| | | | |
|---|---|---|---|
| M9MS48 | Rbp1-like, isoform B OS=Drosophila melanogaster GN=Rbp1-like PE=1 SV=1 | -0.500217879852688 | -0.510457064357526 |
| Q9VHT5 | LD31571p OS=Drosophila melanogaster GN=mRpL1 PE=1 SV=2 | -0.473931188332412 | -0.454031630894707 |
| O02002 | Caspase-1 OS=Drosophila melanogaster GN=Dcp-1 PE=1 SV=1 | -0.467932447710969 | -0.65744525452268 |
| D1FYH5 | Odorant-binding protein 99a OS=Drosophila melanogaster GN=Obp99a PE=4 SV=1 | -0.946193556304206 | -115.521.264.992.094 |
| Q9VVL6 | Mediator of RNA polymerase II transcription subunit 19 OS=Drosophila melanogaster GN=MED19 PE=2 SV=1 | -0.65744525452268 | -0.494109070270043 |
| Q9VUJ0 | 39S ribosomal protein L39, mitochondrial OS=Drosophila melanogaster GN=mRpL39 PE=1 SV=2 | -0.522840788813359 | -0.512513650651464 |
| P49846 | Transcription initiation factor TFIID subunit 5 OS=Drosophila melanogaster GN=Taf5 PE=1 SV=1 | -0.446148031818874 | -0.446148031818874 |
| Q9W1N3 | Levy, isoform A OS=Drosophila melanogaster GN=levy PE=1 SV=1 | -0.554273296650016 | -0.628162382669579 |
| Q9VGP7 | Mitochondrial ribosomal protein L40 OS=Drosophila melanogaster GN=mRpL40 PE=1 SV=1 | -0.554273296650016 | -0.481968507397831 |
| Q9VBP5 | Jing interacting gene regulatory 1, isoform A OS=Drosophila melanogaster GN=jigr1 PE=1 SV=1 | -0.625934281777462 | -0.603840510926846 |
| Q9VFB5 | IP02321p OS=Drosophila melanogaster GN=Rpb7 PE=1 SV=2 | -0.846843211938579 | -0.891642821909581 |
| Q9VIQ8 | CG10664-PA, isoform A OS=Drosophila melanogaster GN=COX4 PE=1 SV=1 | -0.678071905112638 | -0.666576266274808 |
| Q8IML6 | CG11876, isoform B OS=Drosophila melanogaster GN=CG11876 PE=1 SV=1 | -0.586405917590825 | -0.49817873457909 |
| Q9V3W1 | CG4599-PA, isoform A OS=Drosophila melanogaster GN=Tpr2 PE=1 SV=1 | -0.522840788813359 | -0.459972730742493 |
| A4V3G8 | Ballchen, isoform B OS=Drosophila melanogaster GN=ball PE=1 SV=1 | -0.550042516371997 | -0.601649629654035 |
| Q05783 | High mobility group protein D OS=Drosophila melanogaster GN=HmgD PE=1 SV=1 | -1 | -10.350.469.470.992 |
| Q9VHG5 | Insulator binding factor 1, isoform A OS=Drosophila melanogaster GN=Ibf1 PE=1 SV=1 | -108.008.791.132.269 | -116.165.326.347.877 |
| Q7JZD5 | Dorsal interacting protein 3 OS=Drosophila melanogaster GN=Dlip3 PE=1 SV=1 | -0.63262893435147 | -0.556393348524385 |
| Q9VWV8 | Nitric oxide synthase-interacting protein homolog OS=Drosophila melanogaster GN=CG6179 PE=3 SV=1 | -0.531156057025363 | -0.675765437729469 |
| Q9VAY7 | Protein FAM50 homolog OS=Drosophila melanogaster GN=CG12259 PE=2 SV=1 | -0.592919224549499 | -0.603840510926846 |
| Q9VJZ4 | AT12494p OS=Drosophila melanogaster GN=ND-B22 PE=1 SV=1 | -0.795859283219775 | -0.763660460831626 |
| Q9VQ35 | CG17642-PA OS=Drosophila melanogaster GN=mRpL48 PE=1 SV=1 | -0.601649629654035 | -0.564904848379903 |
| Q9VEL2 | Brf, isoform A OS=Drosophila melanogaster GN=Brf PE=1 SV=2 | -0.500217879852688 | -0.575615328461903 |
| Q9VRP5 | Ubiquitin carboxyl-terminal hydrolase 36 OS=Drosophila melanogaster GN=scny PE=1 SV=3 | -0.45205668870965 | -0.407363571393423 |
| Q9VX24 | CG8173-PA OS=Drosophila melanogaster GN=CG8173 PE=1 SV=1 | -0.673462651860048 | -0.72499295250013 |
| Q9VL16 | CG5676-PA OS=Drosophila melanogaster GN=CG5676 PE=1 SV=1 | -0.504304837375931 | -0.436353730515936 |
| Q9V3R3 | CG3704 OS=Drosophila melanogaster GN=EG:BACR7A4.17 PE=1 SV=1 | -0.432454552356253 | -0.496142467422571 |
| Q9VEX9 | Histone deacetylase complex subunit SAP18 OS=Drosophila melanogaster GN=Bin1 PE=1 SV=1 | -0.533242384273829 | -0.539519529959989 |
| Q9W3G1 | CG10555, isoform A OS=Drosophila melanogaster GN=CG10555 PE=1 SV=1 | -0.579921884020626 | -0.522840788813359 |
| Q7K2B0 | Ribosomal RNA-processing protein 8 OS=Drosophila melanogaster GN=CG7137 PE=1 SV=1 | -0.504304837375931 | -0.500217879852688 |
| Q9VSY1 | CG4022, isoform A OS=Drosophila melanogaster GN=CG4022 PE=2 SV=1 | -0.531156057025363 | -0.481968507397831 |
| M9PBC6 | CG17912, isoform F OS=Drosophila melanogaster GN=BuGZ PE=1 SV=1 | -0.488026018218199 | -0.547931769776189 |
| Q9VG76 | C-Myc-binding protein homolog OS=Drosophila melanogaster GN=CG17202 PE=2 SV=2 | -0.641603738043346 | -0.614845103115656 |
| Q9U6L4 | Protein tweety OS=Drosophila melanogaster GN=tty PE=2 SV=1 | -0.510457064357526 | -0.547931769776189 |
| Q9V3W2 | GM23292p OS=Drosophila melanogaster GN=ND-B17 PE=1 SV=1 | -0.706041020971306 | -0.687334826441606 |
| P27716 | Innexin inx1 OS=Drosophila melanogaster GN=ogre PE=1 SV=1 | -0.584241333477502 | -0.753895990116083 |
| P42124 | Histone-lysine N-methyltransferase E(z) OS=Drosophila melanogaster GN=E(z) PE=1 SV=2 | -0.469929257774916 | -0.488026018218199 |

| | | | |
|---|---|---|---|
| Q9VYY4 | Cytochrome P450 4g15 OS=Drosophila melanogaster GN=Cyp4g15 PE=2 SV=1 | -0.409278229990159 | -0.407363571393423 |
| M9PBW0 | Ubiquitin conjugating enzyme 12, isoform B OS=Drosophila melanogaster GN=UbcE2M PE=3 SV=1 | -0.816037165157405 | -0.741782610463982 |
| Q4V5H1 | Peptidyl-prolyl cis-trans isomerase OS=Drosophila melanogaster GN=CG17266 PE=1 SV=1 | -0.56277226108709 | -0.494109070270043 |
| Q9VDL1 | UPF0483 protein CG5412 OS=Drosophila melanogaster GN=CG5412 PE=2 SV=1 | -0.442222328605074 | -0.56277226108709 |
| Q9V444 | Chromatin accessibility complex 14kD protein OS=Drosophila melanogaster GN=Chrac-14 PE=1 SV=1 | -124.127.043.154.214 | -0.910501849160897 |
| A1Z9I6 | CG13344, isoform A OS=Drosophila melanogaster GN=CG13344 PE=1 SV=1 | -0.461958546666336 | -0.479954975960182 |
| Q7KRW4 | CG14516, isoform B OS=Drosophila melanogaster GN=CG14516 PE=1 SV=1 | -0.510457064357526 | -0.617056130431009 |
| A0A0B4KEU2 | Scribbler, isoform J OS=Drosophila melanogaster GN=sbb PE=1 SV=1 | -0.550042516371997 | -0.619270551496451 |
| P43332 | U1 small nuclear ribonucleoprotein A OS=Drosophila melanogaster GN=snf PE=1 SV=1 | -0.486004020632987 | -0.543719518489275 |
| Q9VYB1 | LD33040p OS=Drosophila melanogaster GN=Ndc80 PE=1 SV=1 | -0.422752464406849 | -0.428565884123491 |
| M9PBD6 | Lethal (2) 37Cg, isoform D OS=Drosophila melanogaster GN=l(2)37Cg PE=1 SV=1 | -0.450084446378045 | -0.424687669312563 |
| Q6NP72 | CG13220, isoform A OS=Drosophila melanogaster GN=CG13220 PE=1 SV=1 | -0.554273296650016 | -0.558516520417355 |
| Q24050 | Anon-i1 protein OS=Drosophila melanogaster GN=anon-i1 PE=1 SV=1 | -0.694321256757713 | -0.639354797539784 |
| Q9VHM3 | LD30467p OS=Drosophila melanogaster GN=M1BP PE=1 SV=1 | -0.744197163397282 | -0.708396441969435 |
| Q9VGS3 | RH44771p OS=Drosophila melanogaster GN=SdhC PE=1 SV=2 | -0.550042516371997 | -0.522840788813359 |
| Q9VUA5 | LD42058p OS=Drosophila melanogaster GN=ssp2 PE=1 SV=1 | -0.436353730515936 | -0.407363571393423 |
| X2J6S0 | Male-specific lethal 1, isoform B OS=Drosophila melanogaster GN=msl-1 PE=1 SV=1 | -0.428565884123491 | -0.535331732996556 |
| Q9V998 | Ubiquitin-like protein 5 OS=Drosophila melanogaster GN=ubl PE=3 SV=1 | -0.793356776016605 | -0.828793172581858 |
| Q9VQF5 | Cwc25 OS=Drosophila melanogaster GN=Cwc25 PE=1 SV=1 | -0.446148031818874 | -0.504304837375931 |
| Q8IQI3 | CG10984-PB, isoform B OS=Drosophila melanogaster GN=CG10984 PE=1 SV=2 | -0.529072742524873 | -0.675765437729469 |
| Q7KIN0 | Toll-7 OS=Drosophila melanogaster GN=Toll-7 PE=2 SV=1 | -0.56277226108709 | -0.564904848379903 |
| Q9W5P1 | Mediator of RNA polymerase II transcription subunit 21 OS=Drosophila melanogaster GN=MED21 PE=1 SV=1 | -0.831357964441161 | -0.83650126771712 |
| Q9VXE5 | Serine/threonine-protein kinase PAK mbt OS=Drosophila melanogaster GN=mbt PE=1 SV=2 | -0.603840510926846 | -0.628162382669579 |
| A4V3J9 | Darkener of apricot, isoform N OS=Drosophila melanogaster GN=Doa PE=1 SV=1 | -0.428565884123491 | -0.416962376203336 |
| Q27272 | Transcription initiation factor TFIID subunit 9 OS=Drosophila melanogaster GN=e(y)1 PE=1 SV=1 | -0.454031630894707 | -0.492078535042672 |
| Q9VIJ5 | LD03247p OS=Drosophila melanogaster GN=Pomp PE=1 SV=1 | -0.504304837375931 | -0.569179503480228 |
| Q9VTM2 | CG11652, isoform A OS=Drosophila melanogaster GN=CG11652 PE=1 SV=1 | -0.426625473554056 | -0.518701058452435 |
| Q960C5 | CG6860-PA, isoform A OS=Drosophila melanogaster GN=Lrch PE=1 SV=1 | -0.771027430239839 | -0.630393929968162 |
| A1Z8K9 | Superoxide dismutase [Cu-Zn] OS=Drosophila melanogaster GN=Sod3 PE=1 SV=1 | -0.416962376203336 | -0.461958546666336 |
| Q24423 | Zinc finger protein Noc OS=Drosophila melanogaster GN=noc PE=1 SV=1 | -0.560642821525743 | -0.701341684435485 |
| Q8SWX6 | CG11790, isoform A OS=Drosophila melanogaster GN=CG11790 PE=1 SV=1 | -0.490050853695689 | -0.415037499278844 |
| Q9VUX0 | CG5830, isoform A OS=Drosophila melanogaster GN=CG5830-RA PE=1 SV=2 | -0.518701058452435 | -0.496142467422571 |
| Q9VMB6 | CG31911-PA OS=Drosophila melanogaster GN=Ent2 PE=1 SV=1 | -0.687334826441606 | -0.628162382669579 |
| Q9VLF6 | UPF0585 protein CG18661 OS=Drosophila melanogaster GN=CG18661 PE=2 SV=2 | -0.839079811818897 | -0.736965594166206 |
| Q9VX94 | CG13001-PA OS=Drosophila melanogaster GN=CG13001 PE=1 SV=1 | -0.492078535042672 | -0.552156355637914 |
| Q9VXT5 | CWF19-like protein 2 homolog OS=Drosophila melanogaster GN=CG9213 PE=1 SV=2 | -0.436353730515936 | -0.413115187147815 |
| Q8MKJ6 | CG11777 OS=Drosophila melanogaster GN=CG11777 PE=1 SV=1 | -0.659722595233746 | -0.678071905112638 |

| | | | |
|---|---|---|---|
| P05552 | Transcription factor Adf-1 OS=Drosophila melanogaster GN=Adf1 PE=2 SV=2 | -0.446148031818874 | -0.438307278601691 |
| Q9VW53 | CG8025-PA OS=Drosophila melanogaster GN=Mtr3 PE=1 SV=1 | -0.407363571393423 | -0.518701058452435 |
| Q9V452 | CG15736-PA OS=Drosophila melanogaster GN=Chrac-16 PE=1 SV=1 | -0.888968687611256 | -0.965784284662087 |
| Q9W086 | Mitochondrial ribosomal protein L46 OS=Drosophila melanogaster GN=mRpL46 PE=1 SV=1 | -0.440263475567017 | -0.520769438793664 |
| A0A0B4K6C3 | CG9603, isoform B OS=Drosophila melanogaster GN=COX7A PE=1 SV=1 | -0.701341684435485 | -0.780908941753803 |
| Q8MT36 | Probable histone-lysine N-methyltransferase Mes-4 OS=Drosophila melanogaster GN=Mes-4 PE=1 SV=2 | -0.77349147019132 | -0.935117148415146 |
| Q9VUS0 | CG7372-PA OS=Drosophila melanogaster GN=CG7372 PE=1 SV=2 | -0.810966175609983 | -0.805912947883698 |
| M9PEA2 | CG18292, isoform B OS=Drosophila melanogaster GN=CDK2AP1 PE=1 SV=1 | -0.577766999316952 | -0.582079992188035 |
| Q9VH79 | RE01104p OS=Drosophila melanogaster GN=Rpt3R PE=1 SV=2 | -0.698997743967186 | -0.483984852996335 |
| Q9VKN7 | Aurora kinase B OS=Drosophila melanogaster GN=ial PE=1 SV=1 | -0.524915117051217 | -0.502259911390907 |
| Q9W3V9 | Nuclear factor Y-box C OS=Drosophila melanogaster GN=Nf-YC PE=2 SV=1 | -0.477944250839036 | -0.486004020632987 |
| Q0E8Q7 | CG31731, isoform B OS=Drosophila melanogaster GN=CG31731 PE=1 SV=1 | -0.422752464406849 | -0.473931188332412 |
| Q7JQG5 | Caskin, isoform B OS=Drosophila melanogaster GN=ckn PE=2 SV=1 | -0.409278229990159 | -0.438307278601691 |
| Q9VC49 | DNA-directed RNA polymerases I, II, and III subunit RPABC5 OS=Drosophila melanogaster GN=Rpb10 PE=3 SV=1 | -0.639354797539784 | -0.473931188332412 |
| Q9VPP5 | Ribonuclease H2 subunit A OS=Drosophila melanogaster GN=CG13690 PE=2 SV=1 | -0.646112163715093 | -0.564904848379903 |
| Q7JUZ6 | DnaJ-related co-chaperone MRJ OS=Drosophila melanogaster GN=mrj PE=1 SV=1 | -0.426625473554056 | -0.409278229990159 |
| Q9VG45 | Dpp target protein OS=Drosophila melanogaster GN=Dtg PE=1 SV=1 | -0.483984852996335 | -0.826232932263294 |
| A1ZBW7 | WASH complex subunit FAM21 homolog OS=Drosophila melanogaster GN=CG16742 PE=1 SV=1 | -0.512513650651464 | -0.477944250839036 |
| Q9I7M5 | CG17343-PA OS=Drosophila melanogaster GN=CG17343 PE=1 SV=1 | -0.630393929968162 | -0.648371670897218 |
| Q9VFJ2 | 39S ribosomal protein L11, mitochondrial OS=Drosophila melanogaster GN=mRpL11 PE=1 SV=1 | -0.524915117051217 | -0.646112163715093 |
| Q9V877 | Kinesin-like protein subito OS=Drosophila melanogaster GN=sub PE=1 SV=1 | -0.531156057025363 | -0.556393348524385 |
| O76857 | BCL7-like OS=Drosophila melanogaster GN=BCL7-like PE=1 SV=1 | -0.46394709975979 | -0.560642821525743 |
| Q9VDB7 | CG16791, isoform A OS=Drosophila melanogaster GN=CG16791 PE=1 SV=4 | -0.703689439291908 | -0.648371670897218 |
| Q24297 | Small nuclear ribonucleoprotein F OS=Drosophila melanogaster GN=SmF PE=1 SV=2 | -0.564904848379903 | -0.457989644463391 |
| Q9W4J5 | Ribosomal RNA small subunit methyltransferase NEP1 OS=Drosophila melanogaster GN=CG3527 PE=3 SV=2 | -0.516635639286651 | -0.514573172829758 |
| Q9VJQ4 | mRNA cap guanine-N7 methyltransferase OS=Drosophila melanogaster GN=l(2)35Bd PE=1 SV=2 | -0.63710935733414 | -0.573466861883327 |
| P13008 | 40S ribosomal protein S26 OS=Drosophila melanogaster GN=RpS26 PE=1 SV=1 | -0.486004020632987 | -0.597277823154395 |
| Q9VYW6 | Rhomboid-like protein OS=Drosophila melanogaster GN=rho-4 PE=1 SV=2 | -0.800877357986399 | -0.778432211461591 |
| Q9VAG5 | Claret, isoform A OS=Drosophila melanogaster GN=ca PE=1 SV=4 | -0.424687669312563 | -0.401634794676355 |
| Q7K332 | CG30159, isoform A OS=Drosophila melanogaster GN=CG3364 PE=1 SV=1 | -12.276.920.250.416 | -12.584.251.525.812 |
| E2QCK0 | Transport and golgi organization 11, isoform B OS=Drosophila melanogaster GN=Tango11 PE=1 SV=1 | -0.689659879387849 | -0.763660460831626 |
| Q9VJE4 | DNA-directed RNA polymerase II subunit RPB11 OS=Drosophila melanogaster GN=Rpb11 PE=3 SV=1 | -0.680382065799839 | -0.599462070416271 |
| Q9VUK9 | CG6854-PA, isoform A OS=Drosophila melanogaster GN=CTPsyn PE=2 SV=2 | -0.678071905112638 | -0.558516520417355 |
| Q9VTD6 | CG43693, isoform B OS=Drosophila melanogaster GN=CG6327 PE=1 SV=1 | -0.424687669312563 | -0.477944250839036 |
| Q8SWS7 | LD10447p OS=Drosophila melanogaster GN=vito PE=1 SV=1 | -0.526992432083826 | -0.510457064357526 |
| Q9V4C4 | Synaptotagmin 7, isoform A OS=Drosophila melanogaster GN=Syt7 PE=1 SV=4 | -0.666576266274808 | -0.488026018218199 |
| Q7K172 | LD04933p OS=Drosophila melanogaster GN=ste24a PE=1 SV=1 | -0.729770092762002 | -0.756330919033137 |

| | | | |
|---|---|---|---|
| Q9VJ62 | H/ACA ribonucleoprotein complex non-core subunit NAF1 OS=Drosophila melanogaster GN=CG10341 PE=1 SV=2 | -0.875671864997798 | -0.675765437729469 |
| Q5EAK6 | Serine/threonine-protein kinase ATM OS=Drosophila melanogaster GN=tefu PE=2 SV=1 | -0.524915117051217 | -0.510457064357526 |
| X2JHT7 | Sidekick, isoform G (Fragment) OS=Drosophila melanogaster GN=sdk PE=1 SV=1 | -0.488026018218199 | -0.446148031818874 |
| M9MSM5 | Dishevelled associated activator of morphogenesis, isoform D OS=Drosophila melanogaster GN=DAAM PE=1 SV=1 | -0.524915117051217 | -0.512513650651464 |
| Q9VPH2 | DNA primase large subunit OS=Drosophila melanogaster GN=DNApol-alpha60 PE=1 SV=2 | -0.45600928033515 | -0.434402824145775 |
| Q9VPB8 | CG4289-PA OS=Drosophila melanogaster GN=Pex14 PE=1 SV=1 | -0.434402824145775 | -0.45600928033515 |
| E1JIR3 | Cdc2c, isoform C OS=Drosophila melanogaster GN=Cdk2 PE=1 SV=1 | -0.880975896857725 | -0.915935735211525 |
| G2J5Z0 | CG44242, isoform B OS=Drosophila melanogaster GN=CG8446-RE PE=1 SV=1 | -0.584241333477502 | -0.62148837674627 |
| Q9VH69 | 40S ribosomal protein S29 OS=Drosophila melanogaster GN=RpS29 PE=1 SV=1 | -0.675765437729469 | -0.710755714843358 |
| Q7K284 | Protein CLP1 homolog OS=Drosophila melanogaster GN=cbc PE=2 SV=1 | -0.494109070270043 | -0.512513650651464 |
| P13368 | Protein sevenless OS=Drosophila melanogaster GN=sev PE=1 SV=2 | -0.486004020632987 | -0.599462070416271 |
| Q9XZT7 | Transcription initiation factor TFIID subunit 10b OS=Drosophila melanogaster GN=Taf10b PE=1 SV=1 | -0.432454552356253 | -0.639354797539784 |
| Q9W0M0 | CG13890 OS=Drosophila melanogaster GN=CG13890 PE=1 SV=1 | -0.758769964484555 | -0.788364746672851 |
| Q95T08 | CG9636, isoform C OS=Drosophila melanogaster GN=CG9636 PE=1 SV=1 | -0.422752464406849 | -0.573466861883327 |
| Q9VSJ5 | NADPH-dependent diflavin oxidoreductase 1 OS=Drosophila melanogaster GN=CG13667 PE=1 SV=1 | -0.465938397578882 | -0.508403405589552 |
| Q9VUR7 | Phosducin-like protein OS=Drosophila melanogaster GN=CG7650 PE=1 SV=2 | -0.407363571393423 | -0.41888982477445 |
| Q9XYF4 | Caspase Dronc OS=Drosophila melanogaster GN=Dronc PE=1 SV=1 | -0.409278229990159 | -0.473931188332412 |
| Q9VAN8 | CG11882, isoform A OS=Drosophila melanogaster GN=CG11882-RA PE=1 SV=1 | -0.56277226108709 | -0.758769964484555 |
| Q9VRQ7 | DNA polymerase epsilon subunit 2 OS=Drosophila melanogaster GN=DNApol-epsilon58 PE=1 SV=1 | -0.422752464406849 | -0.502259911390907 |
| Q9VL69 | CG5885-PA OS=Drosophila melanogaster GN=BEST:CK01296 PE=1 SV=1 | -0.547931769776189 | -0.471928835421265 |
| Q8SYK5 | Protein insensitive OS=Drosophila melanogaster GN=insv PE=1 SV=2 | -0.673462651860048 | -0.682695931638085 |
| Q9VU13 | CG42709, isoform A OS=Drosophila melanogaster GN=CG17667 PE=1 SV=3 | -0.436353730515936 | -0.49817873457909 |
| Q8IPV3 | CG3164, isoform C OS=Drosophila melanogaster GN=CG3164 PE=1 SV=1 | -0.405451450449646 | -0.547931769776189 |
| Q24152 | Cyclin-dependent kinases regulatory subunit OS=Drosophila melanogaster GN=Cks30A PE=3 SV=1 | -0.913216233857933 | -0.79836613883035 |
| Q500Y7 | CG14482, isoform A OS=Drosophila melanogaster GN=UQCR-6.4 PE=1 SV=1 | -0.428565884123491 | -0.42081985187285 |
| Q9W3C8 | CG11284, isoform B OS=Drosophila melanogaster GN=CG11284 PE=1 SV=1 | -0.520769438793664 | -0.556393348524385 |
| Q9W3E1 | GH13214p OS=Drosophila melanogaster GN=IntS4 PE=1 SV=1 | -0.744197163397282 | -0.603840510926846 |
| Q9VNG0 | Mediator of RNA polymerase II transcription subunit 27 OS=Drosophila melanogaster GN=MED27 PE=1 SV=1 | -0.428565884123491 | -0.446148031818874 |
| P13002 | Protein grainyhead OS=Drosophila melanogaster GN=grh PE=2 SV=3 | -0.994240730711315 | -0.921390165303633 |
| Q9VRM6 | Lethal (3) persistent salivary gland 2 OS=Drosophila melanogaster GN=l(3)psg2 PE=4 SV=2 | -0.426625473554056 | -0.416962376203336 |
| B7Z043 | Myocardin-related transcription factor, isoform H OS=Drosophila melanogaster GN=Mrtf PE=4 SV=3 | -0.469929257774916 | -0.461958546666336 |
| X2J9B3 | E2F transcription factor 2, isoform B OS=Drosophila melanogaster GN=E2f2 PE=1 SV=1 | -0.430508908041284 | -0.483984852996335 |
| Q9VH38 | Dimethyladenosine transferase 2, mitochondrial OS=Drosophila melanogaster GN=mtTFB2 PE=2 SV=2 | -0.590744853315162 | -0.586405917590825 |
| Q9VMS1 | CG14028-PA OS=Drosophila melanogaster GN=cype PE=1 SV=3 | -0.63262893435147 | -0.641603738043346 |
| Q9GYU7 | Mediator of RNA polymerase II transcription subunit 10 OS=Drosophila melanogaster GN=MED10 PE=1 SV=1 | -0.610433188237274 | -0.584241333477502 |
| P26017 | Polycomb group protein Pc OS=Drosophila melanogaster GN=Pc PE=1 SV=1 | -0.608232280044003 | -0.552156355637914 |
| Q7K4H1 | CG30467 OS=Drosophila melanogaster GN=CG8185 PE=1 SV=1 | -0.588573754273535 | -0.652901329377732 |

| | | | |
|---|---|---|---|
| A1Z7W1 | CG1868, isoform B OS=Drosophila melanogaster GN=Smyd4-1 PE=4 SV=1 | -0.488026018218199 | -0.516635639286651 |
| Q9VAD6 | Conserved oligomeric Golgi complex subunit 7 OS=Drosophila melanogaster GN=Cog7 PE=2 SV=2 | -0.467932447710969 | -0.413115187147815 |
| E1JIB4 | CG2162, isoform E OS=Drosophila melanogaster GN=CG2162 PE=4 SV=2 | -0.57132159005177 | -0.440263475567017 |
| A1Z898 | Caf1-105 OS=Drosophila melanogaster GN=Caf1-105 PE=1 SV=1 | -0.666576266274808 | -0.63262893435147 |
| Q9VY72 | Maternal gene required for meiosis, isoform H OS=Drosophila melanogaster GN=mamo PE=1 SV=4 | -0.98564470702293 | -0.994240730711315 |
| P00408 | Cytochrome c oxidase subunit 2 OS=Drosophila melanogaster GN=mt:CoII PE=3 SV=1 | -108.314.123.530.025 | -109.541.956.507.868 |
| Q7KBL8 | Mediator of RNA polymerase II transcription subunit 29 OS=Drosophila melanogaster GN=ix PE=1 SV=1 | -0.465938397578882 | -0.471928835421265 |
| E1JIP3 | WRN exonuclease, isoform B OS=Drosophila melanogaster GN=WRNexo PE=1 SV=1 | -0.932361283124637 | -0.746615764199925 |
| Q9W2I0 | CG42672, isoform P OS=Drosophila melanogaster GN=CG42672 PE=1 SV=5 | -0.401634794676355 | -0.403541860441014 |
| Q9W0P3 | Mediator of RNA polymerase II transcription subunit 30 OS=Drosophila melanogaster GN=MED30 PE=1 SV=1 | -0.457989644463391 | -0.473931188332412 |
| P49906 | Transcription initiation factor TFIID subunit 11 OS=Drosophila melanogaster GN=Taf11 PE=1 SV=1 | -130.044.836.747.691 | -12.584.251.525.812 |
| Q8T9H8 | CG31908, isoform B OS=Drosophila melanogaster GN=ade3 PE=2 SV=1 | -109.850.554.495.243 | -108.008.791.132.269 |
| Q9VD55 | Cytochrome c heme lyase, isoform A OS=Drosophila melanogaster GN=Cchl PE=1 SV=1 | -0.753895990116083 | -0.623709616662948 |
| Q9W554 | CG14814, isoform B OS=Drosophila melanogaster GN=CG14814 PE=1 SV=1 | -148.196.850.739.783 | -136.959.452.851.768 |
| P56175 | Probable RNA 3'-terminal phosphate cyclase-like protein OS=Drosophila melanogaster GN=Rtc1 PE=2 SV=3 | -0.41119543298445 | -0.428565884123491 |
| Q9W0S5 | Miple, isoform A OS=Drosophila melanogaster GN=miple1 PE=2 SV=2 | -0.682695931638085 | -0.713118852211838 |
| Q6IKC0 | CG42394, isoform A OS=Drosophila melanogaster GN=CG42394 PE=1 SV=1 | -0.440263475567017 | -0.403541860441014 |
| Q9W020 | Nucleolar MIF4G domain-containing protein 1 homolog OS=Drosophila melanogaster GN=CG9004 PE=2 SV=1 | -0.446148031818874 | -0.543719518489275 |
| Q6IDF5 | CG12859 OS=Drosophila melanogaster GN=ND-B15 PE=1 SV=1 | -0.488026018218199 | -0.533242384273829 |
| Q9W141 | Putative ATP synthase subunit f, mitochondrial OS=Drosophila melanogaster GN=CG4692 PE=1 SV=1 | -0.746615764199925 | -0.696657605512669 |
| Q9W547 | LD21404p OS=Drosophila melanogaster GN=mRpL16 PE=1 SV=1 | -0.518701058452435 | -0.486004020632987 |
| Q9VM17 | SOSS complex subunit B homolog OS=Drosophila melanogaster GN=CG5181 PE=2 SV=1 | -0.535331732996556 | -0.608232280044003 |
| A0A0B4KFV4 | CG15107, isoform B OS=Drosophila melanogaster GN=CG15107 PE=1 SV=1 | -0.810966175609983 | -0.778432211461591 |
| Q9VQY9 | Probable DNA replication complex GINS protein PSF2 OS=Drosophila melanogaster GN=Psf2 PE=2 SV=1 | -0.438307278601691 | -0.440263475567017 |
| Q9W282 | tRNA pseudouridine synthase OS=Drosophila melanogaster GN=CG3045 PE=1 SV=1 | -1,98850436116217 | -2,07704103576383 |
| Q9V4B6 | CG31998-PA OS=Drosophila melanogaster GN=CG31998 PE=1 SV=2 | -0.558516520417355 | -0.520769438793664 |
| Q9W3S4 | CG4617, isoform A OS=Drosophila melanogaster GN=CG4617 PE=1 SV=2 | -0.454031630894707 | -0.514573172829758 |
| A1ZAW5 | Methylosome subunit pICln OS=Drosophila melanogaster GN=icln PE=1 SV=1 | -0.543719518489275 | -0.481968507397831 |
| Q9VNP3 | Mesoderm-expressed 2, isoform B OS=Drosophila melanogaster GN=Mes2 PE=1 SV=1 | -0.703689439291908 | -0.727379545337008 |
| Q8SZR6 | CG31223 OS=Drosophila melanogaster GN=syndapin PE=1 SV=1 | -0.461958546666336 | -0.56277226108709 |
| Q7JZ53 | CG4866 OS=Drosophila melanogaster GN=CG4866 PE=1 SV=1 | -0.652901329377732 | -0.706041020971306 |
| Q9VAS7 | Innexin inx3 OS=Drosophila melanogaster GN=Inx3 PE=1 SV=1 | -0.935117148415146 | -0.862496476250065 |
| Q9VMJ5 | Beta galactosidase, isoform A OS=Drosophila melanogaster GN=Gal PE=1 SV=2 | -0.461958546666336 | -0.500217879852688 |
| Q9VVX0 | Protein Gemin2 OS=Drosophila melanogaster GN=Gem2 PE=1 SV=1 | -0.407363571393423 | -0.502259911390907 |
| Q07886 | Probable ATP-dependent RNA helicase Dbp45A OS=Drosophila melanogaster GN=Dbp45A PE=2 SV=2 | -0.63710935733414 | -0.560642821525743 |
| Q9VM45 | Nuf2 OS=Drosophila melanogaster GN=Nuf2 PE=4 SV=2 | -0.965784284662087 | -0.951763814347152 |
| Q9VCG0 | CG13599 OS=Drosophila melanogaster GN=CG13599 PE=1 SV=1 | -0.599462070416271 | -0.558516520417355 |

| | | | |
|---|---|---|---|
| Q9VMH4 | CG9175, isoform A OS=Drosophila melanogaster GN=CG9175-RA PE=1 SV=1 | -0.46394709975979 | -0.446148031818874 |
| Q9VIK0 | CG9319, isoform B OS=Drosophila melanogaster GN=CG9319-RA PE=1 SV=1 | -0.590744853315162 | -0.524915117051217 |
| A1ZAK1 | FI09622p OS=Drosophila melanogaster GN=IntS8 PE=1 SV=1 | -0.524915117051217 | -0.664288089679538 |
| Q9VLN0 | CG13390 OS=Drosophila melanogaster GN=CG13390 PE=4 SV=1 | -0.438307278601691 | -0.601649629654035 |
| Q9NK54 | Protein chiffon OS=Drosophila melanogaster GN=chif PE=1 SV=2 | -0.67116353577046 | -0.617056130431009 |
| O46079 | CG3587-PA OS=Drosophila melanogaster GN=EG:39E1.2 PE=1 SV=1 | -220.423.305.221.761 | -212.029.423.371.771 |
| Q8IPS4 | CG11109, isoform B OS=Drosophila melanogaster GN=CG11109-RB PE=1 SV=1 | -0.531156057025363 | -0.494109070270043 |
| Q8IMI5 | FI04474p OS=Drosophila melanogaster GN=spdo PE=2 SV=1 | -0.560642821525743 | -0.436353730515936 |
| Q2PDX2 | CG34001, isoform B OS=Drosophila melanogaster GN=CG34001 PE=1 SV=2 | -0.678071905112638 | -0.582079992188035 |
| O17468 | Protein HIRA homolog OS=Drosophila melanogaster GN=Hira PE=1 SV=2 | -0.968604803724466 | -0.749038426466781 |
| A0A0B4KEN4 | Van gogh, isoform B OS=Drosophila melanogaster GN=Vang PE=4 SV=1 | -0.448114896528275 | -0.592919224549499 |
| Q9VTY6 | Ubiquitin-conjugating enzyme E2 C OS=Drosophila melanogaster GN=vih PE=1 SV=1 | -0.477944250839036 | -0.508403405589552 |
| Q9VH45 | CG5359, isoform A OS=Drosophila melanogaster GN=Dlc90F PE=1 SV=1 | -0.680382065799839 | -0.816037165157405 |
| M9PFW3 | Frizzled 2, isoform F OS=Drosophila melanogaster GN=fz2 PE=4 SV=1 | -0.63710935733414 | -0.586405917590825 |
| Q8IP99 | Transcriptional adaptor 1-1, isoform A OS=Drosophila melanogaster GN=Ada1-1 PE=4 SV=1 | -0.558516520417355 | -0.430508908041284 |
| Q9W4L1 | 39S ribosomal protein L33, mitochondrial OS=Drosophila melanogaster GN=mRpL33 PE=3 SV=2 | -0.432454552356253 | -0.422752464406849 |
| Q9VSW5 | Kinesin-like protein OS=Drosophila melanogaster GN=Klp67A PE=1 SV=1 | -0.415037499278844 | -0.45205668870965 |
| M9NEL5 | Erect wing, isoform K OS=Drosophila melanogaster GN=ewg PE=4 SV=1 | -0.496142467422571 | -0.434402824145775 |
| Q7JW61 | CG8494 OS=Drosophila melanogaster GN=Usp20-33 PE=1 SV=1 | -0.514573172829758 | -0.45205668870965 |
| E1JIZ7 | CG42487 OS=Drosophila melanogaster GN=CG42487 PE=1 SV=1 | -0.768567591552035 | -0.826232932263294 |
| Q9VG07 | CG7488 OS=Drosophila melanogaster GN=CG7488 PE=1 SV=2 | -0.65744525452268 | -0.751465163861321 |
| A0A0B4JD21 | CG10253, isoform B OS=Drosophila melanogaster GN=CG10253 PE=1 SV=1 | -0.768567591552035 | -0.803392955905182 |
| Q7JWH6 | CG1888, isoform A OS=Drosophila melanogaster GN=CG1888 PE=1 SV=1 | -0.753895990116083 | -0.639354797539784 |
| Q9VAQ5 | Probable dimethyladenosine transferase OS=Drosophila melanogaster GN=CG11837 PE=2 SV=1 | -0.552156355637914 | -0.518701058452435 |
| Q9VHW6 | CG10903, isoform A OS=Drosophila melanogaster GN=CG10903 PE=1 SV=1 | -0.522840788813359 | -0.541617995843987 |
| M9PIU6 | Zeste, isoform C OS=Drosophila melanogaster GN=z PE=4 SV=1 | -114.880.066.140.671 | -105.889.368.905.357 |
| Q0KI97 | CG34117 OS=Drosophila melanogaster GN=CG34117 PE=1 SV=1 | -0.582079992188035 | -0.575615328461903 |
| Q9VRH6 | Translation factor waclaw, mitochondrial OS=Drosophila melanogaster GN=waw PE=3 SV=2 | -0.564904848379903 | -0.541617995843987 |
| C8VV95 | Daughterless, isoform B OS=Drosophila melanogaster GN=da PE=1 SV=1 | -0.516635639286651 | -0.617056130431009 |
| Q9XZ53 | Oligosaccharyl transferase 3 OS=Drosophila melanogaster GN=OstStt3 PE=1 SV=1 | -0.763660460831626 | -0.739372091873301 |
| Q9VL63 | UPF0430 protein CG31712 OS=Drosophila melanogaster GN=CG31712 PE=1 SV=3 | -0.520769438793664 | -0.403541860441014 |
| Q9W256 | IP14609p OS=Drosophila melanogaster GN=Mes4 PE=1 SV=1 | -0.554273296650016 | -0.502259911390907 |
| Q9VKC2 | Amino acid transporter protein JHI-21 OS=Drosophila melanogaster GN=JhI-21 PE=1 SV=1 | -0.524915117051217 | -0.510457064357526 |
| Q9V6K3 | Tyrosine-protein kinase transmembrane receptor Ror2 OS=Drosophila melanogaster GN=Nrk PE=1 SV=2 | -0.428565884123491 | -0.438307278601691 |
| Q9NJB5 | Homeobox protein onecut OS=Drosophila melanogaster GN=onecut PE=2 SV=2 | -0.543719518489275 | -0.481968507397831 |
| Q9Y171 | BcDNA.GH02220 OS=Drosophila melanogaster GN=BcDNA.GH02220 PE=1 SV=1 | -1 | -0.785875194647153 |
| Q9VKN8 | Palmitoyltransferase OS=Drosophila melanogaster GN=Dnz1 PE=1 SV=1 | -0.41119543298445 | -0.438307278601691 |

| | | | |
|---|---|---|---|
| Q9VBF0 | CG5447, isoform A OS=Drosophila melanogaster GN=CG5447 PE=1 SV=1 | -0.547931769776189 | -0.514573172829758 |
| Q6NL34 | AT03686p OS=Drosophila melanogaster GN=WDR79 PE=1 SV=1 | -0.727379545337008 | -0.63486740654747 |
